# Predicting functional constraints across evolutionary timescales with phylogeny-informed genomic language models

**DOI:** 10.1101/2025.09.21.677619

**Authors:** Chengzhong Ye, Gonzalo Benegas, Carlos Albors, Jianan Canal Li, Sebastian Prillo, Peter D. Fields, Brian Clarke, Yun S. Song

## Abstract

Genomic language models (gLMs) have emerged as a powerful approach for learning genome-wide functional constraints directly from DNA sequences. However, standard gLMs adapted from natural language processing often require extremely large model sizes and computational resources, yet still fall short of classical evolutionary models in predictive tasks. Here, we introduce GPN-Star (Genomic Pretrained Network with Species Tree and Alignment Representation), a biologically grounded gLM featuring a phylogeny-aware architecture that leverages whole-genome alignments and species trees to model evolutionary relationships explicitly. Trained on alignments spanning vertebrate, mammalian, and primate evolutionary timescales, GPN-Star achieves state-of-the-art performance across a wide range of variant effect prediction tasks in both coding and non-coding regions of the human genome. Analyses across timescales reveal task-dependent advantages of modeling more recent versus deeper evolution. To demonstrate its potential to advance human genetics, we show that GPN-Star substantially outperforms prior methods in prioritizing pathogenic and fine-mapped GWAS variants; yields unprecedented enrichments of complex trait heritability; and improves power in rare variant association testing. Extending beyond humans, we train GPN-Star for five model organisms – *Mus musculus, Gallus gallus, Drosophila melanogaster, Caenorhabditis elegans*, and *Arabidopsis thaliana* – demonstrating the robustness and generalizability of the framework. Taken together, these results position GPN-Star as a scalable, powerful, and flexible new tool for genome interpretation, well suited to leverage the growing abundance of comparative genomics data.

## Introduction

A fundamental challenge in biology is understanding the functional significance of genetic variants. Despite tremendous advances in experimental technologies in genomics, determining which variants influence phenotypes or contribute to diseases remains difficult. Evolution provides valuable insights into this question: deleterious mutations tend to be purged by natural selection, while tolerated or advantageous changes may accumulate. Leveraging millions of years of such natural experiments offers a genome-wide, *in vivo* record of functional constraint, providing a unique and highly informative perspective for variant interpretation. This idea of learning from evolution has deep roots, beginning with mathematical models of molecular evolution and early work on comparative sequence analysis [1, 2].

Genomic language models (gLMs) have recently emerged as a promising approach for extracting genome-wide evolutionary information directly from raw DNA sequences (see [3] and references therein). These are large-scale deep learning models trained with self-supervision objectives on unlabeled sequences – a paradigm that has driven recent breakthrough in AI and machine learning (ML), especially in natural language processing [4, 5]. By learning to predict masked nucleotides from sequence context, gLMs assign per-site likelihoods that quantitatively reflect the functional constraint for every allele, without requiring supervision on labeled variants. These likelihoods have been shown to be effective predictors of genome-wide variant effects [3, 6]. However, gLMs based on standard language modeling frameworks still underperform compared to much simpler classical phylogenetic models on certain variant interpretation tasks – particularly in complex eukaryotic genomes such as in humans and in distal regulatory elements such as enhancers [7] – even with massive model sizes [8, 9].

A long-standing approach to modeling evolutionary data is to construct multiple sequence alignments (MSAs). By algorithmically aligning homologous loci across biological sequences, MSAs reveal site-specific patterns of conservation and facilitate the inference of evolutionary preferences for variants at specific positions. Protein MSAs have been foundational to several prominent deep learning models for protein sequences, including AlphaFold [10], MSA Transformer [11], and EVE [12]. More recently, upon noticing diminishing returns from scaling protein language models, a new wave of methods based on MSAs (or, more generally, homology search) has emerged [13–20]. Extending this concept from proteins to genomes, whole-genome alignments (WGAs) of complete assemblies from tens to hundreds of species enable genome-wide studies of sequence evolution [21, 22]. Classical parametric phylogenetic models fitted on WGAs – such as GERP [23], PhastCons [24], and PhyloP [25] – have long been staples in variant interpretation for humans and other species. Reflecting the importance of these resources, several large consortia have recently invested in constructing high-quality, large-scale WGA datasets [26–28].

Therefore, combining the gLM framework with whole-genome alignment data presents a promising direction. We recently demonstrated this potential with GPN-MSA [29], a transformer model trained on a WGA of 90 vertebrate species. GPN-MSA achieves strong performance across several variant effect prediction tasks. However, it uses a generic architecture more suitable for unaligned sequences, which limits its ability to fully exploit the rich structure inherent in alignment data, particularly the evolutionary relationship between species. Furthermore, conditioning on species too closely related to humans was empirically shown to deteriorate performance in variant effect prediction, possibly because the model’s learned probability distribution becomes overly biased toward their genomes. This prompted us to exclude most primates from the training data – a suboptimal decision, since primates offer valuable insight into recent evolutionary constraints relevant to humans. Lastly, GPN-MSA is highly tuned for a particular alignment, which limits its adaptability to new data.

Here, we present GPN-Star (**G**enomic **P**retrained **N**etwork with **S**pecies **T**ree and **A**lignment **R**epresentations), a general and powerful gLM framework that better utilizes multispecies WGA information through a new model architecture with phylogeny-aware designs. This flexible framework enabled us to apply GPN-Star to three WGAs relative to the human genome, spanning vertebrate, mammal, and primate evolutionary timescales. GPN-Star achieves state-of-the-art performance across a broad range of variant interpretation benchmarks, spanning both coding and non-coding regions. The vertebrate model consistently outperforms GPN-MSA trained on the same alignment, while the mammal and primate models excel particularly at predicting non-coding variant effects. We further evaluate its utility in human genetics by analyzing the heritability of complex traits. Remarkably, the GPN-Star model trained on the primate alignment prioritizes variants with unprecedented levels of heritability enrichment across more than a hundred complex traits [30]. In addition, we find a striking connection between the most informative evolutionary timescale for constraint prediction and the effective polygenicity of traits.

To further demonstrate the generality of our approach, we apply GPN-Star to five model organisms – including mouse (*M. musculus*), fruit fly (*D. melanogaster*), chicken (*G. gallus*), *C. elegans*, and *A. thaliana* – with minimal tuning, and show its effectiveness in assessing variant effects in these species.

## Results

### Alignment- and phylogeny-informed genomic language models

GPN-Star learns the functional constraints of genetic variants in a target genome by leveraging evolutionary signals embedded in multispecies phylogeny and whole-genome alignment data. Inspired by classical evolutionary models, it aims to characterize how each genomic position evolves across species. We introduce a specialized transformer architecture designed to capture highly expressive functions that describe the evolutionary information, while effectively incorporating the genomic context of each site (Figure 1A).

**Figure 1:**
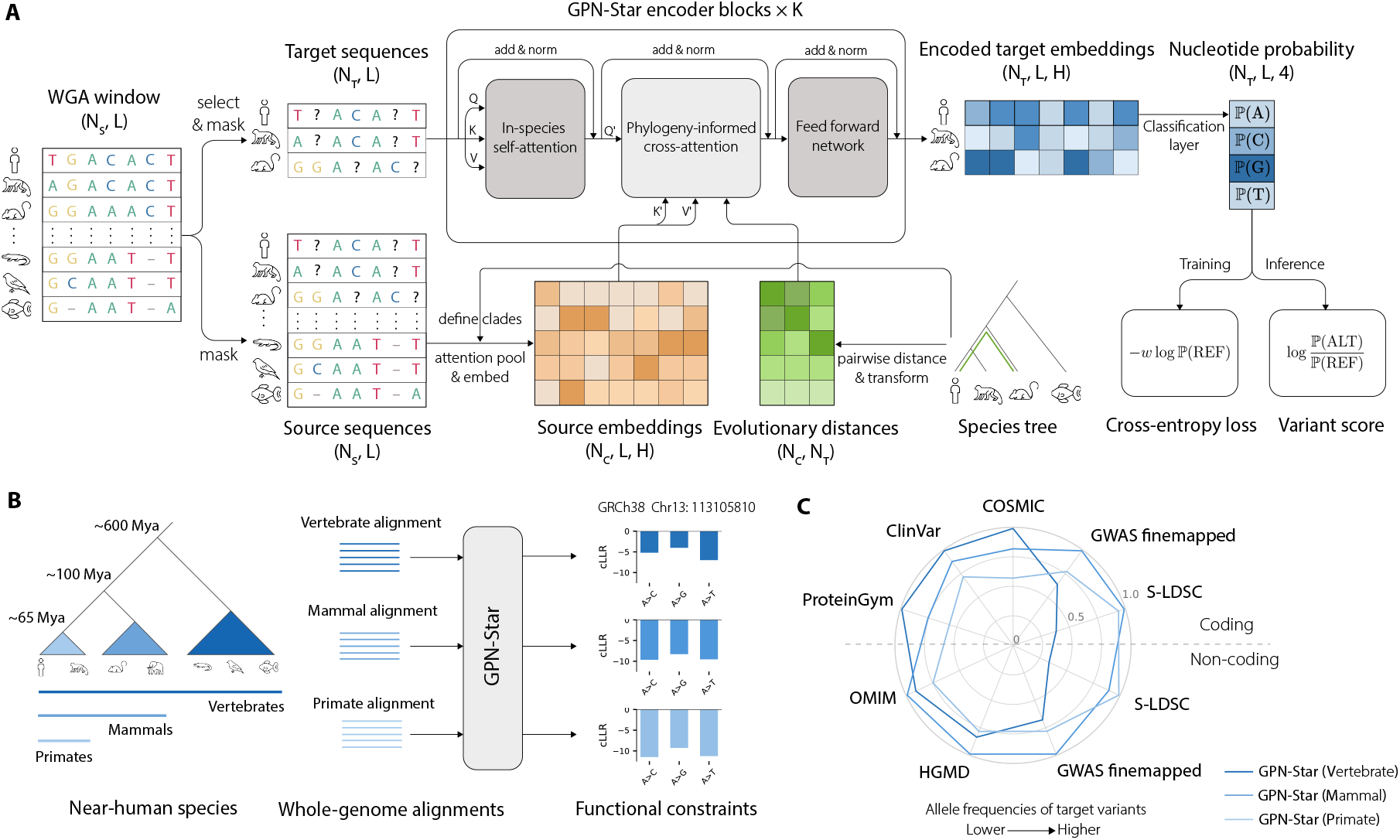
Overview of GPN-Star. **(A)** A diagram of the GPN-Star model architecture. The input to the model is a whole-genome alignment window. The target sequences and source sequences are constructed from the alignment window. The source sequences are compressed into clade-level embeddings via attention pooling following the species tree. The target sequences are encoded through a stack of GPN-Star encoder blocks, where the phylogeny-informed cross-attention module integrates information from the source sequences guided by evolutionary distances between species based on the species tree. Finally, a classification layer transforms the encoded embeddings to nucleotide probabilities at each locus in the target sequences, which are then used to compute the training loss or to make inferences on variant impact. A full description of the model is provided in Methods. **(B)** Application of GPN-Star to the human genome. Three models were trained on vertebrate, mammal, and primate alignments, respectively, learning functional constraints at different evolutionary timescales. Mya: million years ago. cLLR: calibrated log-likelihood ratio (see Methods). **(C)** A summary of performance across the downstream tasks of the vertebrate, mammal, and primate GPN-Star models. For ClinVar, COSMIC, OMIM, HGMD, and GWAS fine-mapped datasets, the performance metric was AUPRC. For ProteinGym, the metric was the mean Spearman’s *ρ* across assays. For S-LDSC, the metric was heritability enrichment. The performance metrics were scaled linearly to the range of 0 to 1, where in each task 0 was defined as the lowest metric among the three GPN-Star models and PhyloP and PhastCons fitted at the same three evolutionary timescales, and 1 was defined as the highest metric.

The model uses an encoder-only architecture trained with a masked language modeling (MLM) objective. Each input consists of a WGA window across multiple species and a corresponding species tree. From this alignment, we construct two sets of sequences: *target sequences* and *source sequences*, with each sequence corresponding to a species. The model learns to predict masked nucleotides in the target sequences, conditioned on their own sequence context and the evolutionary context from the source sequences. This is accomplished through a stack of encoder blocks, each comprising a sequence-wise self-attention module to encode intra-sequence context, a phylogeny-informed cross-attention module to encode evolutionary context from the source sequences, and a feed-forward network to integrate the information. At the core of the model, the cross-attention module adaptively weights source sequence contributions using evolutionary distances derived from the species tree via an attention mechanism. The resulting encoder embeddings for the target sequences integrate rich information from both intra-sequence context and evolutionary information, and are passed through an MLM head to produce the final per-nucleotide probability predictions used for training and downstream variant scoring. Full architectural and training details are provided in Methods.

The GPN-Star framework advances beyond GPN-MSA in three key aspects: First, whereas GPN-MSA masks only the human genome during training, GPN-Star leverages all genomes in the alignment by predicting masked nucleotides across multiple species, thereby substantially increasing both the amount and diversity of training data. Second, GPN-Star explicitly incorporates phylogenetic relationships among species through specialized attention modules, enabling more accurate and biologically grounded modeling. Third, it flexibly accommodates alignments of arbitrary composition and size, eliminating the need for manual curation, such as excluding closely related species, as was required in GPN-MSA [29]. Empirically, we observe consistent and substantial performance gains across extensive evaluations (Supplementary Figure 1).

An additional methodological innovation of this work is accounting for mutation rate variation for the first time in gLMs. We developed an effective calibration procedure that removes mutation rate effects from the raw model outputs, yielding more accurate quantification of genome-wide functional constraints (detailed below and in Methods).

The GPN-Star framework is general and flexible, designed to work with any alignment data from any species, requiring minimal hyperparameter tuning to achieve robust performance (Methods). We first applied our framework to the human genome, training three separate GPN-Star models using the largest available vertebrate, mammalian, and primate WGAs (Figure 1B, Methods). Throughout this study, we denote these models as GPN-Star (V), GPN-Star (M), and GPN-Star (P), respectively. We have experimented with different model sizes, but we focus here on the largest models, which have 200 million parameters and were trained for several days on 8 NVIDIA A100 GPUs (Supplementary Table 1), representing a significantly more efficient resource footprint than previous gLMs – such as Nucleotide Transformer (trained on 128 A100 GPUs for a month) [8] and Evo-2 (trained on over 2,000 H200 GPUs for several months) [9].

In contrast to previous gLMs trained on data from extremely broad evolutionary timescales (e.g., spanning prokaryotes to humans), GPN-Star focuses on narrower, more recent phylogenetic distances closely related to humans (Figure 1B). As we demonstrate below, modeling longer evolutionary histories is not always optimal, depending on the downstream application. Instead, capturing recent evolutionary constraints proves especially advantageous for interpreting certain classes of genetic variants (Figure 1C).

### GPN-Star shows strong predictive capability for pathogenic coding variants

A critical challenge in human genetics is understanding the impact of genetic variants on disease susceptibility – an essential step toward improving the diagnosis of genetic disorders, identifying drug targets, and realizing the promise of precision medicine. Numerous computational models have been developed to predict variant pathogenicity in the human genome. Here, we systematically evaluate the performance of GPN-Star across a comprehensive set of benchmarks.

Our evaluation of coding variants focused on missense variants, the most prevalent class. We first considered classifying pathogenic vs. benign variants in ClinVar, a widely-used clinical variant database containing expert-curated pathogenicity labels [31]. Among all major genome-wide variant effect predictors – including classical evolutionary methods (PhyloP, PhastCons), the ensemble model CADD, recent gLMs (Nucleotide Transformer 2.5B multispecies model, Evo-2 40B model, and GPN-MSA) – GPN-Star (V) achieved the highest area under the precision-recall curve (AUPRC), matching the performance of the protein language model ESM-1b (Figure 2A). For ease of visualization, only the best-performing GPN-Star model is shown in Figure 2; results for all three versions are available in Supplementary Figure 2.

**Figure 2:**
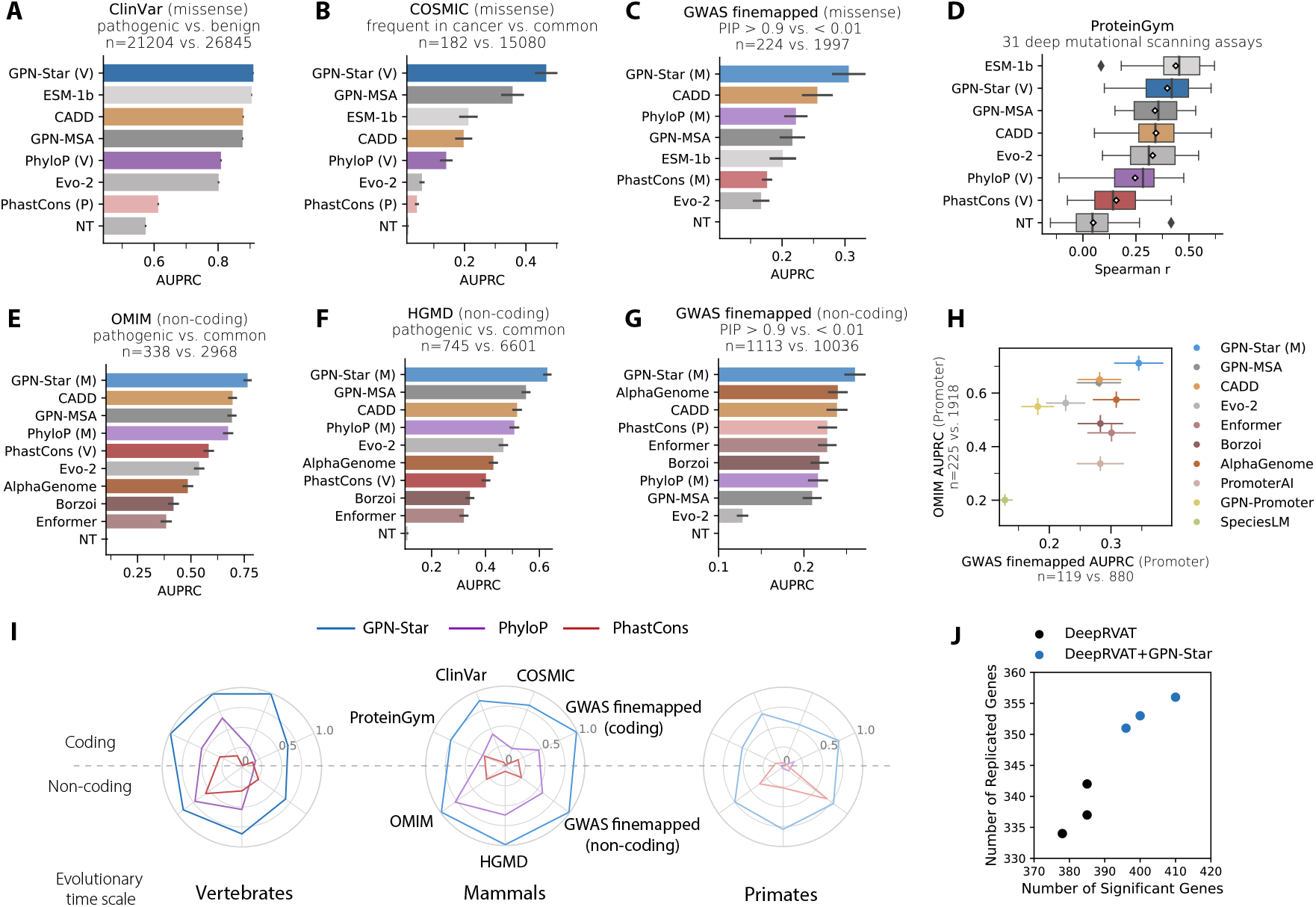
Performance of GPN-Star on human genome-wide variant effect prediction. **(A)** Classification of ClinVar pathogenic versus benign missense variants. **(B)** Classification of COSMIC high-frequency missense variants (*>*0.1%) versus gnomAD v3 common missense variants. **(C)** Classification of GWAS fine-mapped putatively causal (PIP*>*0.9) versus non-causal (PIP*<*0.01) missense variants across 66 traits in UK Biobank. **(D)** Boxplots showing Spearman correlations with deep mutational scanning fitness scores of 31 assays on human proteins in ProteinGym. **(E)** Classification of OMIM pathogenic versus gnomAD v3 common non-coding variants. **(F)** Classification of HGMD pathogenic versus gnomAD v3 common non-coding variants. **(G)** Classification of GWAS fine-mapped putatively causal (PIP*>*0.9) versus non-causal (PIP*<*0.01) non-coding variants across 84 traits in UK Biobank. **(H)** Classification of OMIM pathogenic versus gnomAD v3 common (*y* axis) and GWAS fine-mapped putatively GWAS fine-mapped causal versus non-causal (*x* axis) promoter variants. The performance metric used in (A)-(C) and (E)-(G) is the area under the precision-recall curve (AUPRC). The error bars represent the standard errors from 1000 bootstrap resamples. **(I)** Radar plots comparing performances of GPN-Star against PhyloP and PhastCons at the vertebrate, mammal, and primate timescales on the seven pathogenicity prediction benchmarks. The performance metrics were scaled linearly to the range of 0 to 1, where 0 was defined as the lowest metric across the nine models, and 1 was defined as the highest metric. **(J)** Rare variant association testing on 34 quantitative traits in UK Biobank with WES data from 161,822 unrelated individuals of European ancestry, comparing original DeepRVAT versus DeepRVAT with the three GPN-Star predictions (V, M, and P) on the number of significant genes (*x* axis) and the number of significant genes that replicated discoveries from previous studies with large sample sizes (*y* axis).

We next considered somatic cancer variants from the COSMIC database [32], constructing a benchmark to distinguish missense variants frequently observed in tumors from common missense variants in the general population (gnomAD [33]). As shown in Figure 2B, GPN-Star (V) substantially outperformed all competing models, demonstrating strong predictive capability for pathogeniticy beyond germline variants.

Beyond clinical variants, we also assessed performance on deep mutational scanning (DMS) data, functional assays that measure the fitness effects of all possible missense mutations in a given protein. Across 31 human DMS datasets from ProteinGym [34], GPN-Star (V) outperformed all genome-wide models, though it slightly lagged behind the protein-specific model ESM-1b (Figure 2D).

We further compared GPN-Star with two recent missense variant effect predictors, AlphaMissense [35] and PrimateAI-3D [36]. These models were supervised on population allele frequency data, a highly informative signal for pathogenicity prediction that is also used to define labels in databases such ClinVar [37]. Interestingly, although both AlphaMissense and PrimateAI-3D achieved top performance on ProteinGym and ClinVar, a simple post hoc adjustment of GPN-Star (V) predictions using gnomAD allele frequencies boosted its performance on ClinVar to surpass PrimateAI-3D and approach that of AlphaMissense (Supplementary Figure 3).

### GPN-Star is the state-of-the-art predictor of pathogenic non-coding variants

We next considered tasks regarding non-coding variants, which are known to be especially challenging for predictive modeling. Here, we demonstrate that GPN-Star is a powerful tool for identifying pathogenic non-coding variants in the human genome.

In addition to previous genome-wide evolutionary models, we included three prominent sequence-to-function models – Enformer [38], Borzoi [39], and AlphaGenome [40] – which are trained on extensive functional genomics data and have been widely applied to non-coding variant interpretation. We evaluated the models on classifying non-coding pathogenic variants from the OMIM [41] and HGMD [42] databases, both of which contain expert-curated annotations of human disease-associated variants. GPN-Star (M) achieved the best performance on both benchmarks (Figure 2E, F). Notably, the sequence-to-function models performed substantially worse than evolutionary models on this task, consistent with recent findings [7, 43].

Given the critical role of promoter regions in transcription initiation and gene regulation, many specialized models have been developed to predict the impact of promoter variants. We therefore evaluated GPN-Star on promoter variants in OMIM and compared its performance with other methods, including three promoter-specific models: PromoterAI [44], SpeciesLM [45], and GPN-Promoter [7]. As shown in Figure 2H, GPN-Star (M) demonstrated superior predictive performance compared to all competing models. In particular, the margin of improvement over the promoter-specific models is rather substantial.

### GPN-Star is able to identify fine-mapped putatively causal variants in GWAS

Genome-wide association studies (GWAS) have been instrumental in identifying variants that contribute to genetic disease susceptibility. To further assess the utility of GPN-Star, we evaluated its performance in classifying putatively causal vs. non-causal missense variants resulting from fine-mapping of GWAS variants across 65 traits from the UK Biobank [46]. Among all competing models, GPN-Star (M) achieved the highest predictive performance on these fine-mapped missense variants (Figure 2C). Notably, despite leveraging population allele frequency information, both AlphaMissense and PrimateAI-3D were substantially outperformed by GPN-Star in this task (Supplementary Figure 3).

We next evaluated the models on fine-mapped non-coding GWAS variants across 83 traits from the UK Biobank [46], again assessing their ability to distinguish putatively causal vs. non-causal variants. GPN-Star (M) continued to outperform all other models on this benchmark (Figure 2G). While sequence-to-function models (Enformer, Borzoi, and AlphaGenome) exhibited moderate performance, Evo-2 showed relatively limited predictive value, as was previously observed [7]. For fine-mapped variants located in promoter regions, GPN-Star (M) again outperformed all models, including the promoter-specific models PromoterAI, SpeciesLM, and GPN-Promoter (Figure 2H).

### Relevant evolutionary timescales for different variant effect prediction tasks

The above results demonstrate that GPN-Star is a powerful and versatile framework for genome-wide variant interpretation. Moreover, our analysis underscores the importance of the timescale represented in the training data of evolutionary models. Across benchmarks, we observed a consistent pattern: Coding variants and lower-frequency variants with larger effect sizes tended to be better predicted by models trained on deeper evolutionary timescales. Such variants are enriched in the ClinVar, COSMIC, and ProteinGym benchmarks. In contrast, non-coding variants and higher-frequency variants with smaller effect sizes were, in general, more accurately predicted by models trained on shallower evolutionary timescales, as seen in the OMIM, HGMD, and fine-mapped GWAS benchmarks. A similar trend was also observed with PhyloP and PhastCons scores at the three evolutionary timescales, although both were consistently outperformed by GPN-Star at each timescale (Figure 2I). We further discuss potential explanations of this phenomenon in the Discussion section.

### GPN-Star improves rare variant association testing

Having observed the strong performance of GPN-Star in predicting pathogenic and fine-mapped variants, we further explored its utility in rare variant association testing (RVAT), an important but challenging task in statistical genetics. The great interest in RVAT stems from the fact that most large-effect variants tend to be rare, as they are subject to negative selection pressure [47]. To overcome statistical power issues with low-frequency variants, RVAT is typically carried out at the gene level by aggregating variants within each gene [48]. These testing procedures often rely on variant annotations that reflect functional importance to prioritize variants in the aggregation. DeepRVAT [49] is a recent method that employs a deep set network to integrate such variant annotations for rare variant association testing. By combining an expressive deep learning framework with a powerful set of variant annotations, it demonstrated improved statistical power and computational efficiency compared with previous methods in extensive evaluations on UK Biobank whole-exome sequencing (WES) data [49].

We conducted RVAT experiments using DeepRVAT, enhancing the annotations used in the published version with all three GPN-Star predictions during both the training and association testing phases. Following the benchmarking procedure of [49], we performed RVAT experiments using DeepRVAT on 34 quantitative traits in UK Biobank with the WES data from 161,822 unrelated individuals of European ancestry, retaining variants with minor allele frequency (MAF) *<* 0.1% (Methods). To account for differences in stochastic model initialization, this procedure was run three times with different random seeds. As shown in Figure 2J, adding the predictions from the GPN-Star models as variant annotations into DeepRVAT resulted in an increase in the number of discovered genes at a family-wise error rate *<* 0.05 (on average across runs, 402 genes compared with 383 from the original method). Among the significant discoveries, there were also more gene-phenotype associations that replicated in conventional RVAT studies on UK Biobank with larger sample sizes (Methods) [50, 51] (on average 353 compared with 338), indicating high robustness of the additional discoveries. Notably, the original DeepRVAT already includes state-of-the-art missense variant effect predictors such as AlphaMissense and PrimateAI, as well as the sequence-to-function model DeepSEA, in the annotations. Nonetheless, GPN-Star appeared to offer complementary information and further boosted the power of the tests.

### GPN-Star predictions are highly informative for complex trait heritability

To further investigate the prediction of causal variants for complex trait GWAS, we turned to stratified LD score regression (S-LDSC) [52]. S-LDSC is a principled way to estimate the informativeness of an annotation for complex trait heritability while leveraging the signal from all SNPs, including those not confidently fine-mapped or genome-wide significant. It also serves as a backbone for functionally-informed fine-mapping [53] and polygenic risk scores [54]. Because S-LDSC requires scoring ∼10 million variants, it was feasible to evaluate only the most scalable models (or those with precomputed scores). We binarized model scores to select a specific fraction of common variants. In our main analysis, we used the top 0.1% most constrained variants. We ran S-LDSC separately for each model score, while conditioning on 96 baseline features, and meta-analyzed the results across 106 independent traits [30].

Since the early work on S-LDSC, conservation scores have been found to be the most enriched annotation for complex trait heritability [52]. More recently, primate-specific conservation – particularly PhastCons (P) – has emerged as the state-of-the-art for complex trait heritability enrichment [55, 56]. Notably, GPN-Star (P) significantly improves on this result, followed by GPN-Star (M) (Figure 3A). These improvements are even more striking when examining the heritability coefficient *τ*^⋆^ (Figure 3A), which quantifies the unique contribution of an annotation to heritability after adjusting for baseline features (Methods). While per-trait enrichment estimates are noisier, GPN-Star (P) or GPN-Star (M) consistently ranks at the top (Supplementary Figure 4). This advance is particularly notable given the long-standing role of complex trait heritability enrichment as a meaningful benchmark in human genetics.

**Figure 3:**
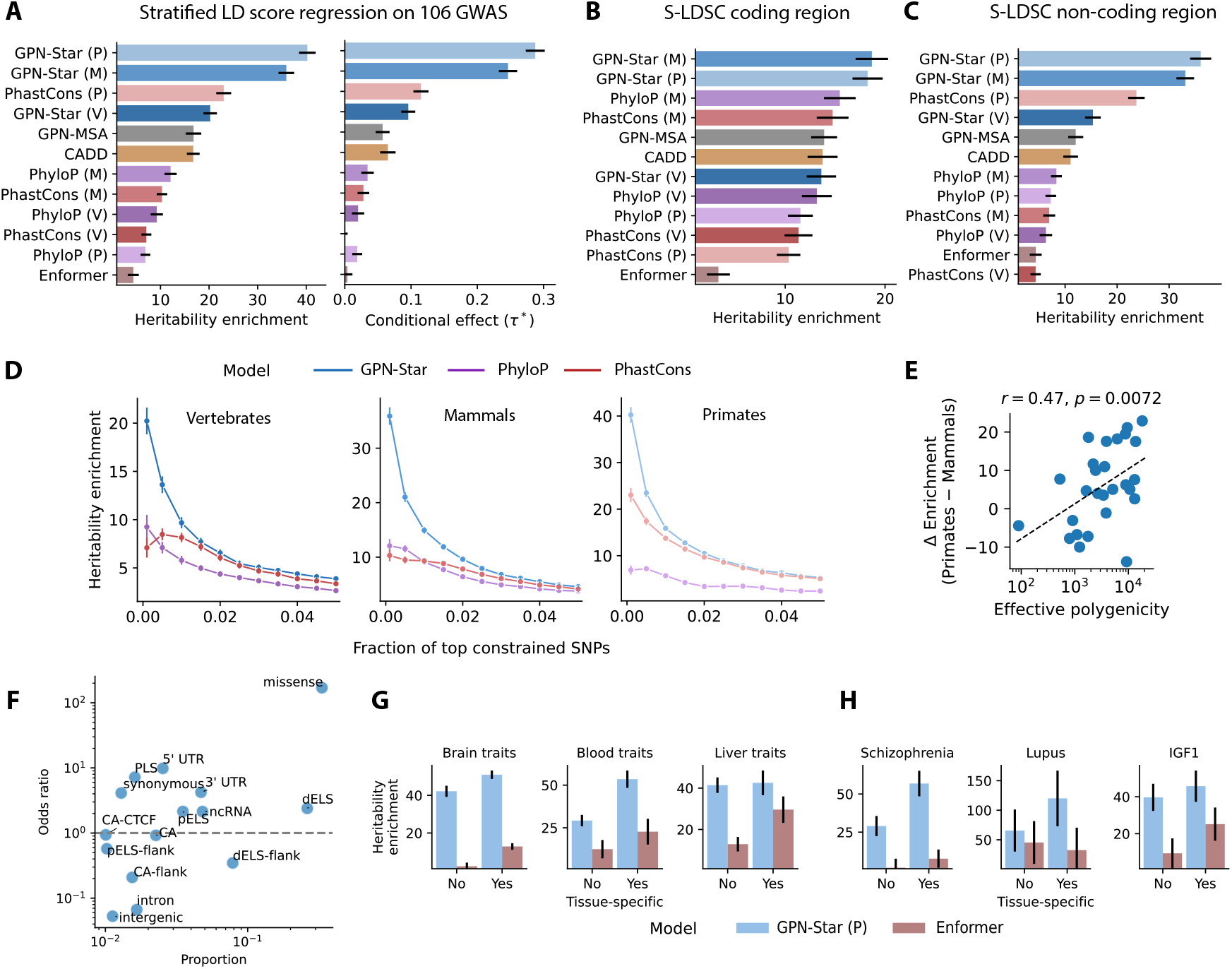
SNP heritability analyses in human complex traits. **(A)** Informativeness of different models for complex trait heritability. We binarized model scores to select the top 0.1% most constrained common variants. We ran S-LDSC separately for each model annotation, while conditioning on 96 baseline features, and meta-analyzed the results across 106 independent traits. Heritability enrichment is the proportion of heritability explained by an annotation divided by the size of the annotation. The conditional effect (*τ*^⋆^) measures the unique contribution to heritability that is not explained by existing annotations. The error bars represent standard errors. **(B)** Performance restricting model annotations to coding regions. **(C)** Performance restricting model annotations to non-coding regions. **(D)** Comparison as the fraction of top constrained SNPs is varied. **(E)** Difference in enrichment between GPN-Star (P) and (M) as a function of estimated effective polygenicity, for 27 traits. *p*-value is one-sided. Dashed line is ordinary least squares fit. **(F)** Type of common variants prioritized by GPN-Star (P). Variants are annotated by a combination of Ensembl consequences and ENCODE SCREEN candidate *cis*-regulatory elements. Only types with proportion above 1% are displayed. Odds ratio is with respect to the bottom 99.9% of common variants. **(G)** Comparison of tissue-agnostic and tissue-specific adaptations of GPN-Star (P) and Enformer across grouped traits. **(H)** Comparison of tissue-agnostic and tissue-specific adaptations of GPN-Star (P) and Enformer across individual traits.

The improvement of GPN-Star persists when restricting to top-ranked variants in coding and non-coding regions separately (Figure 3B, C), with stronger gains in non-coding regions. Moreover, across a range of binarization thresholds and all three evolutionary timescales, GPN-Star consistently outperforms previous conservation scores (Figure 3D).

### Evolutionary timescales and the genetic architecture of complex traits

We then analyzed how GPN-Star models trained on different evolutionary timescales prioritize variants. Among common variants prioritized by GPN-Star (P), 39% are shared with GPN-Star (M) and (V). Additionally, 59% are shared only with GPN-Star (M), and 36.6% are unique to GPN-Star (P) (Supplementary Figure 5, see Supplementary Figure 6 for all S-LDSC variants). Motivated by the observation that GPN-Star (M) performs better on Mendelian traits while GPN-Star (P) excels on polygenic traits (Figure 1C, Figure 2I), we inspected their relative heritability enrichment performance against a recently estimated spectrum of trait polygenicity [57]. Across 27 traits with available polygenicity estimates, we found a moderate correlation (Pearson’s *r* = 0.47) between the (log) effective polygenicity and the heritability enrichment difference between GPN-Star (P) and GPN-Star (M) (Figure 3E, Supplementary Figure 7). For example, GPN-Star (M) attains a greater enrichment for LDL cholesterol (effective polygenicity = 89), while GPN-Star (P) attains a higher enrichment for schizophrenia (effective polygenicity = 13, 069). This suggests a meaningful connection between evolutionary timescale and the genetic architecture of traits, and strengthens the evidence for the special role of primate-specific evolutionary constraint acting on human complex trait variation.

To better understand the kinds of variants prioritized by the top model GPN-Star (P), we analyzed Ensembl variant consequences [58] and ENCODE SCREEN candidate cis-regulatory elements (cCREs, Supplementary Table 2) [59]. Results for common variants are summarized in Figure 3F, focusing on consequence classes each comprising *>* 1% of prioritized variants. Full results for common variants and results including low-frequency variants are in Supplementary Tables 3-4 and Supplementary Figures 8-9.

Missense variants represent the majority (33%, 170-fold enriched), followed by variants in regions with distal enhancer-like signatures (dELS) (26%, 2.4-fold enriched) and their flanks (8%, 0.35-fold depleted). The importance of distal enhancer variants for complex trait heritability should not be understated, as they remain a major weakness of current sequence-to-function models [60] (including the latest AlphaGenome [43]), as well as alignment-free gLMs (including the largest Evo-2 [7]).

There are interesting differences in the variant types prioritized by the three GPN-Star models. There are only slight differences between GPN-Star (P) and (M), with the former enriched 1.14-fold in missense variants (Supplementary Table 5). Not surprisingly, there are far more differences between GPN-Star (P) and (V) (Supplementary Table 6). GPN-Star (P) is generally enriched in exonic variants, such as missense (1.52-fold) and even synonymous (3.89-fold), while it is depleted in non-exonic variants such as intergenic.

### Tissue-specific predictions further improve heritability enrichment

As the final analysis of complex trait heritability, we investigated the performance of tissue-specific scores. Sequence-to-function models such as Enformer [38] do not provide a single variant effect score, but instead predict changes in activity across thousands of functional genomics tracks representing different assays and tissues. In our comparisons, we leveraged careful aggregations of Enformer scores across all tracks (tissue-agnostic) as well as within nine specific tissues, recently generated by Fabiha *et al*. [61]. Given the low enrichment of tissue-agnostic Enformer when meta-analyzed across all traits (Figure 3A), we investigated the performance of tissue-specific Enformer scores. We meta-analyzed these scores only across traits where we expected the tissue to be relevant (Supplementary Table 7). Unsurprisingly, tissue-specific Enformer score consistently outperforms tissue-agnostic Enformer score (Figure 3G, Supplementary Figure 10).

Motivated by this observation, we devised a simple approach to incorporate tissue-specificity into GPN-Star by only considering top-scoring variants located near tissue-specific genes (inspired by LDSC-SEG [62]). While this improved performance for some tissues – such as brain and blood/immune – it did not consistently surpass the performance of tissue-agnostic GPN-Star annotations, possibly because the way we are incorporating the tissue-specificity of a trait is not optimal (Figure 3G, Supplementary Figure 10). Nevertheless, GPN-Star outperformed Enformer across all nine tissues.

Zooming in on example traits, brain-specific GPN-Star showed the highest enrichment for schizophrenia, blood/immune-specific GPN-Star performed best for lupus, and both tissue-agnostic and liver-specific GPN-Star yielded similar results for IGF1 (Figure 3H). These findings highlight the importance of incorporating tissue-specific information in variant effect prediction.

### GPN-Star learns functional elements in the genome and their dependencies

gLMs can learn powerful sequence representations by predicting masked nucleotides, as local nucleotide distributions in a region are heavily influenced by the region’s function. While previous work has shown that gLM embeddings can distinguish genomic regions without supervision [6], it has remained unclear whether alignment-based models could do the same. To investigate this, we visualized embeddings of genomic windows from gene regions, cCREs, and background regions (Figure 4A). We highlight the GPN-Star (M) model with 85 M parameters here and include the remaining models in the supplement.

**Figure 4:**
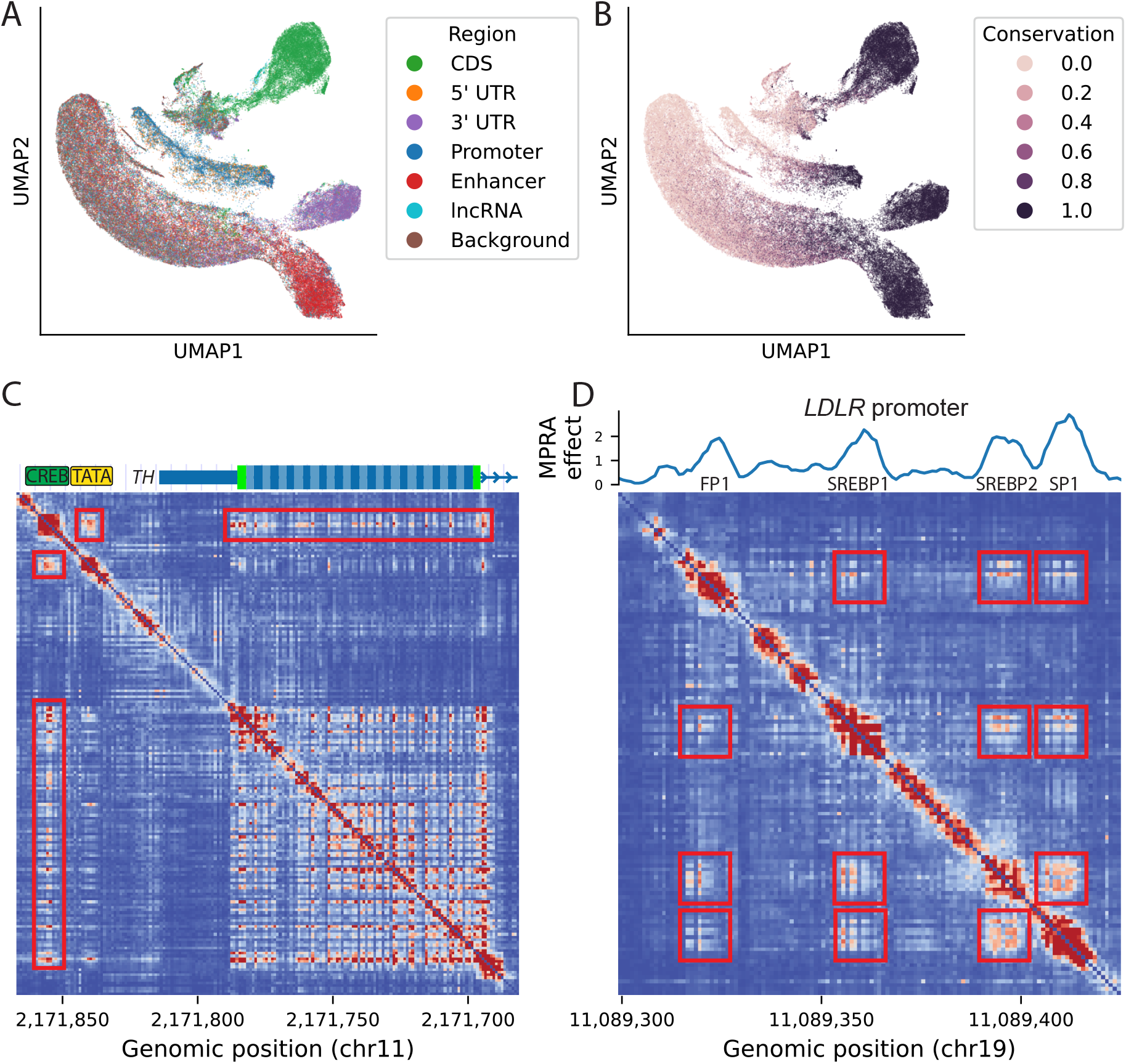
Interpretation of GPN-Star reveals functional elements and their dependencies. **(A)** Visualization of embeddings of different genomic windows, colored by their annotated region. **(B)** Visualization of embeddings of different genomic windows, colored by conservation, defined as the 75^th^ percentile of PhastCons (P) in the window. **(C)** Nucleotide dependency map in *TH* promoter. The intensity of the heatmap at position *i, j* corresponds to the influence of the nucleotide at position *i* on the predicted nucleotide probabilities at position *j*. **(D)** Nucleotide dependency map in *LDLR* promoter. MPRA effect refers to smoothed absolute log fold change.

The resulting embeddings largely segregate by genomic region (Figure 4A, Supplementary Figure 11). For example, coding sequences (CDS) and enhancers form distinct clusters. On the other hand, promoter and 5’ UTR windows are difficult to distinguish, perhaps because 5’ UTRs embeddings encode a mix of transcriptional and post-transcriptional signals. Embeddings for conserved windows show stronger clustering by functional region compared to those for non-conserved windows (Figure 4B). These results suggest that GPN-Star is aware of core functional elements of the genome when making predictions.

Site-independent models such as phyloP cannot, by definition, capture dependencies among nucleotides. In contrast, alignment-free gLMs have been shown to learn dependencies among wellknown interacting elements, e.g., within a transcription factor binding site (TFBS) motif or between splice donors and acceptors [45]. To further probe GPN-Star’s understanding of genomic syntax, we analyzed learned nucleotide dependencies by systematically mutating each position and quantifying the resulting probability changes at other positions in the sequence [45, 63]. In the promoter and first exon of *TH*, which encodes the enzyme tyrosine hydroxylase, this analysis revealed a strong interaction block within the coding region and another at a binding site of the transcription factor CREB, where mutations are known to cause tyrosine hydroxylase deficiency and dystonia [44, 64–67] (Figure 4C, Supplementary Figure 12). The model predicts that CREB depends on both the TATA box and, interestingly, the coding region.

We also observed predicted dependencies across exons in *HBA1* (Supplementary Figure 13), a gene with exceptionally small introns that fit within the model’s context size. Notably, dependencies between splice donor and acceptor regions were especially strong, consistent with previous findings [45].

We next examined the *LDLR* promoter, implicated in familial hypercholesterolemia, which contains well-known TFBS and has been studied using massively parallel reporter assays (MPRA) [55, 68, 69]. TFBS locations can be predicted well from the block structure in the nucleotide dependency map (Figure 4D, Supplementary Figure 14) [45]. Furthermore, the model also predicts dependencies between TFBS, including the well-known interaction between SREBP2 and SP1 [70].

Finally, we analyzed an accessible region hypothesized to be under primate-specific constraint [56]. Supporting this hypothesis, the GPN-Star (P) model showed the highest level of dependency around a putative TEAD4 binding site (Supplementary Figure 15).

These results demonstrate that GPN-Star can leverage co-evolutionary signals to learn meaningful nucleotide dependencies that align with known functional dependencies. This represents a notable advance over traditional conservation scores such as PhyloP and PhastCons.

### GPN-Star learns genome-wide evolutionary constraints

To more directly assess the connection between model predictions and evolutionary constraints in the genome, we leveraged allele frequency data from gnomAD v3.1.2, which aggregates whole-genome sequencing samples from 76,156 human individuals [33]. Allele frequencies in the human population serve as informative indicators of selective constraint: more deleterious alleles tend to have lower frequencies due to purifying selection [6, 29, 33].

In this evaluation, we focused on comparison with PhyloP and PhastCons, which also learn evolutionary constraints from WGA data. To assess how well each model captures the relationship between allele frequency and constraint, we considered the vertebrate, mammal, and primate versions of each model, and obtained predictions for all gnomAD v3 variants on chromosome 22, the held-out chromosome not used in training GPN-Star. We then compared the mean allele frequencies in several quantile bins defined by different models. As shown in Figure 5A, across all three evolutionary timescales, variants in lower GPN-Star quantile bins have consistently lower average allele frequencies compared to those in corresponding PhyloP and PhastCons bins, suggesting that GPN-Star more accurately captures selective constraints in the human genome.

**Figure 5:**
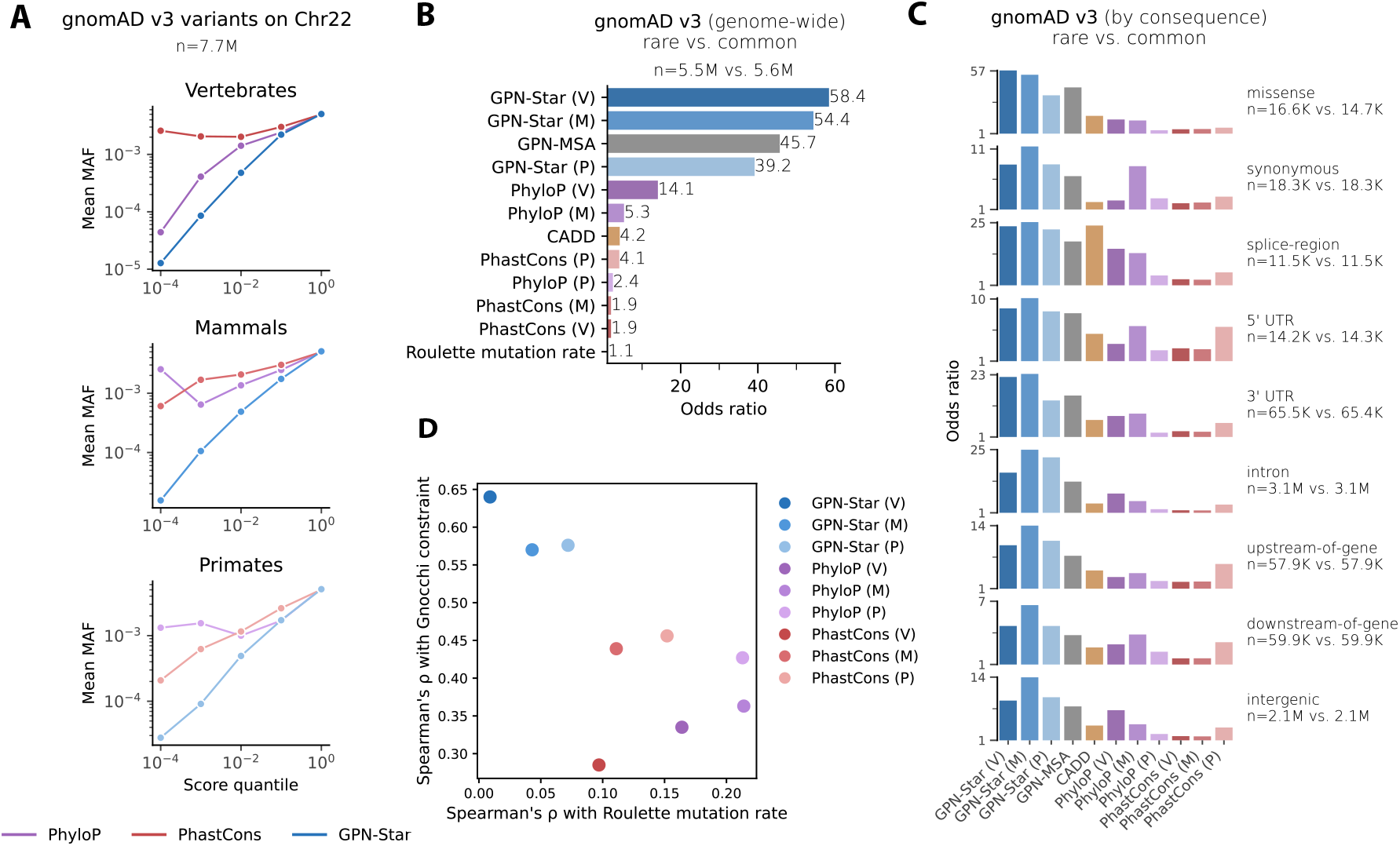
GPN-Star scores reflect evolutionary constraints on the human genome. **(A)** Mean MAF for quantile bins ([0, 10^4^), (10^4^, 10^3^], …, (10^1^, 1]) defined by GPN-Star, PhyloP, and PhastCons based on the vertebrate, mammal, and primate alignments at the gnomAD biallelic sites on chromosome 22. **(B and C)** Enrichment of rare (singletons) versus common (MAF*>*5%) gnomAD variants in the tail of deleterious scores (the threshold was chosen such that each score made 30 false discoveries). (B) shows the genome-wide enrichment, and (C) shows the enrichments stratified by molecular consequences of the variants. The rare variants were downsampled to match the number of common variants in each category. **(D)** Performance comparison of GPN-Star with PhyloP and PhastCons at the three evolutionary timescales on correlation with Roulette mutation rate estimates on chromosome 22 (*x* axis) and correlation with Gnocchi constraint estimates (*y* axis).

We next performed a more quantitative evaluation focusing on the most deleterious tails of the model score distributions, a regime especially important in many human genetics applications. We quantified the enrichment of rare variants relative to common variants in the most constrained tail predicted by each model. Here, we defined rare variants as singletons and common variants as those with allele frequency *>* 5%. Because rare variants are, on average, more deleterious, a model with more accurate constraint predictions should show higher enrichment of singletons in its most constrained tail. As shown in Figure 5B, all three GPN-Star models yielded substantially higher enrichment of rare variants than PhyloP, PhastCons, or CADD. Among the GPN-Star models, the vertebrate model showed the strongest enrichment overall and outperformed GPN-MSA (also trained on vertebrate genomes). When stratifying variants by molecular consequences, GPN-Star again achieved the highest enrichment in every category (Figure 5C). Notably, GPN-Star (V) performed the best for missense variants, whereas GPN-Star (M) led for the synonymous and non-coding categories, mirroring trends observed in previous benchmarks.

### Controlling for mutation rate variation improves variant effect prediction

We studied the influence of context-dependent mutation rate variation on model predictions. Because GPN-Star is trained on genomic sequences observed in nature, its predictions inherently reflect both mutation and selection processes. As expected, the raw model scores showed moderate correlations with mutation rate estimates from Roulette [71] (Spearman’s *ρ* = 0.31-0.34 across the models for chromosome 22). To isolate the impact of selection, we designed a simple yet effective calibration procedure to remove the influence of mutation rate variation from our model scores (Methods). This calibration procedure reduced the correlation with mutation rate estimates to negligible levels, lower than those observed for PhyloP and PhastCons at the same evolutionary timescales, and improved performance across most downstream benchmarks (Figure 5D, Supplementary Figure 16).

We further validated our calibration procedure by comparing our GPN-Star scores to Gnocchi, a genome-wide constraint score estimated from gnomAD v3 that explicitly accounts for mutation rate variation. Calibrated GPN-Star scores showed higher concordance with Gnocchi than either PhyloP or PhastCons (Figure 5D).

Accordingly, all analyses presented in this article used the calibrated scores. As a final sanity check, we confirmed that Roulette mutation rates themselves exhibit negligible enrichment of singletons in their low-rate tails (Figure 5B) and near-baseline performance on pathogenicity prediction tasks (Supplementary Figure 2). This confirms that our rare variant enrichment and pathogenicity benchmarks primarily assess selective constraint rather than mutational bias.

These findings highlight the critical importance of addressing mutation rate variation in future development of gLMs, particularly for applications involving functional constraint prediction.

### GPN-Star is a general and flexible framework for learning evolutionary constraints across diverse species and alignments

Major experimental efforts to understand genetic variation have focused primarily on the human genome [72, 73]. In contrast, researchers have had limited means to investigate the functional impact of variants at the genome-wide scale in other species. We propose GPN-Star to be a powerful and general framework for deciphering genetic variation across diverse species. GPN-Star requires only a multispecies WGA to obtain genome-wide variant effect predictions, and such data are increasingly available for many species [26–28].

As a demonstration, we trained GPN-Star models on five important model organisms: *M. musculus, G. gallus, D. melanogaster, C. elegans*, and *A. thaliana*. In each case, we show that the learned constraints are highly informative for interpreting genetic variation in that species. For each of the five species, we collected a WGA dataset comprising aligned genomes from 18 to 135 species and applied training procedures similar to those used for the human models, with only minor adjustments (Supplementary Table 1).

Due to the scarcity of large-scale curated variant datasets with functional annotations in non-human species, we used population genetic data for evaluation. Specifically, we gathered population-level variant data for each of the five species (Methods) and examined the enrichment of rare variants in the most deleterious tail of the prediction distributions – mirroring the gnomAD-based analysis used for the human models. Across all five species, GPN-Star scores exhibited substantially higher rare variant enrichment compared to PhyloP and PhastCons (Figure 6A), indicating more accurate predictions of genome-wide evolutionary constraints. Across different variant categories based on molecular consequences, GPN-Star consistently showed the highest enrichment in almost all cases (Supplementary Figure 17). In *A. thaliana*, the performance gap between GPN-Star and other models was smaller compared to the results for the other species, which could be due to either the small alignment or the known lower quality of WGAs in plants [74]

**Figure 6:**
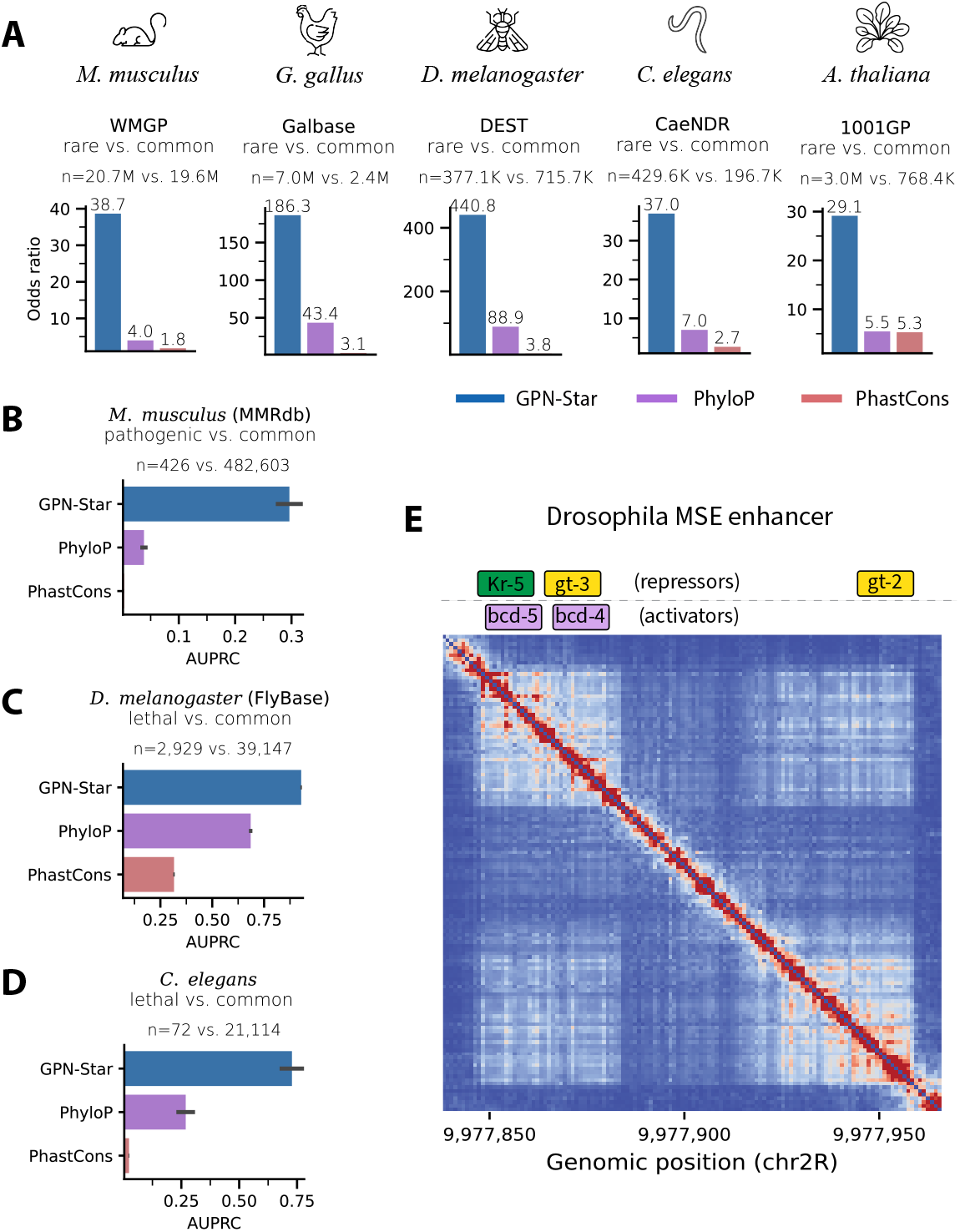
Application of GPN-Star to non-human species. **(A)** Evaluation of GPN-Star models for five non-human species on enrichment of rare versus common variants (thresholds in Methods) in five corresponding population genome databases in the tail of deleterious scores (the threshold was chosen such that each score made 30 false discoveries), compared with PhyloP and PhastCons fitted to the same alignments. Classification of MMrdb pathogenic variants versus WMGP common variants in *M. musculus*. **(C)** Classification of FlyBase lethal variants versus DEST common variants in *D. melanogaster*. **(D)** Classification of *C. elegans* lethal variants versus CaeNDR common variants. The performance metric used in (B)-(D) is the area under the precision-recall curve (AUPRC). The error bars represent the standard errors from 1000 bootstrap resamples. **(E)** Nucleotide dependency map of the *D. melanogaster* model at a locus within the MSE enhancer annotated with known TFBS.

For three of the species, we were able to gather curated pathogenic variants to carry out further evaluation. For the *M. musculus* model, we used pathogenic variants from the MMRdb database (Methods). GPN-Star outperformed both PhyloP and PhastCons in distinguishing these pathogenic variants from common variants in the population, both genome-wide (Figure 6B) and within each variant category (Supplementary Figure 17). Similarly, we collected experimentally validated lethal variants for *D. melanogaster* (from FlyBase [75]) and for *C. elegans* (from an experimental study [76]), and found that GPN-Star consistently outperformed the other models on both datasets (Figure 6C, D and Supplementary Figure 17).

Beyond variant prioritization, GPN-Star also enables exploration of learned functional elements and their co-evolution through nucleotide dependency analysis. As an example, we highlight learned nucleotide dependencies between TFBS at the well-characterized MSE enhancer locus in *D. melanogaster* (Figure 6E). The coordinated activity of these transcription factors drives precise spatial patterning across the fly embryo [77, 78]. This showcases how GPN-Star can serve as an efficient and low-cost tool for investigating functional elements and their dependencies across genomes, complementing and guiding experimental studies.

## Discussion

GPN-Star represents a novel genomic language modeling framework, equipped with powerful inductive biases grounded in evolutionary biology. It consistently outperforms existing methods in predicting deleterious variants across the entire human genome. Furthermore, it achieves top performance in classifying fine-mapped GWAS variants, as well as improving variant prioritization in rare variant association testing. Evaluated across 106 complex traits, GPN-Star substantially advances the state-of-the-art in heritability enrichment, and our analysis offers new insights into the importance of primate-specific constraint and tissue-specificity. Model interpretation further reveals that GPN-Star learns to recognize a wide range of functional elements in the genome – from enhancers to individual TFBS and their co-evolutionary dependencies – all without any supervision. GPN-Star is a general and scalable framework, requiring only aligned genome sequences and relatively modest GPU resources to train an effective model. Classical WGA-based models such as PhastCons [24] and PhyloP [25] have been indispensable tools to biologists and clinicians for over two decades. Our results suggest that GPN-Star can complement and extend these tools, offering improved performance across diverse species and alignments.

A central insight from our results is the strong dependence of learned constraint on the evolutionary timescale represented in the WGA training data. Broadly, the observed patterns align with established views of genome evolution. Coding variants and rarer, larger-effect variants are more accurately assessed by models trained on deeper timescales. These variants tend to occur at highly constrained loci, where evolutionary change is slower, allowing homologous sequences from distantly related species to retain informative genomic syntax. A parallel example is seen in protein language models, which benefit from extremely diverse training sequences spanning from prokaryotes to humans [79].

In contrast, non-coding and more common, smaller-effect variants are better captured at shallower timescales. These typically occur in faster-evolving loci, where sequence context shifts more rapidly. Some regulatory elements, for instance, have been shown to be constrained only within primates [56]. Taken together, our findings emphasize the evolutionary timescale as a critical consideration in the development and application of gLMs.

GPN-Star achieves state-of-the-art performance with substantially smaller model and context sizes compared to existing single-sequence gLMs. This efficiency likely stems from the explicit homology information provided by the alignment. Notably, alignment-based models for proteins have similarly outperformed single-sequence models with much fewer parameters (e.g., AlphaFold [10] vs. ESMFold [80], or MSA Transformer [11] vs. ESM-1b [79]). Although we observed modest performance gains from increased model sizes, improvements from expanding context sizes were relatively small (Supplementary Figure 18). A plausible explanation is that WGAs are constructed relative to a reference species and often consist of small, highly fragmented synteny blocks. As a result, when context size increases, nucleotides within a window may not be contiguous in the actual genomes of non-reference species, introducing misleading context information to the model. Exploring how to leverage larger context sizes in WGA data is a promising direction for future research.

While this study focuses on variant interpretation, gLMs have broader potential applications, including genome annotation and functional element discovery [3], which remain to be explored. Our model interpretation results suggest GPN-Star’s potential in these tasks. Extending the context size might be particularly useful in this scenario and warrants further investigation.

Our results on pathogenicity prediction and complex trait heritability demonstrate GPN-Star’s utility in human genetics. Initial experiments in rare variant association testing showed noticeable improvements, though current analyses were limited to exome data. We anticipate even greater benefits in non-coding regions, where variant prioritization has historically been more difficult. GPN-Star predictions could also enhance other downstream applications such as functionally informed fine-mapping, polygenic risk scoring, and phenotype prediction. We are actively exploring these directions in collaboration with domain experts.

Beyond identifying evolutionary constraints and deleterious variants, an important future direction is to study human-specific adaptations. The current GPN-Star models focus on capturing evolutionary signals across species, which lack the resolution for studying selection within human populations. We anticipate this to be an important direction for future work and would require learning from even more localized evolutionary data, such as human population genomes and archaic human genomes.

It is remarkable how much can be learned from unlabeled DNA sequences alone. Functional genomics and sequence-to-function models offer a complementary lens on genetic variation. Although they underperformed our evolutionary approach in pathogenicity and heritability analyses, they remain indispensable for studying gene regulatory mechanisms, tissue and cell specificity, and molecular phenotypes such as gene expression. In our tissue-specific heritability analysis, GTEx-derived tissue-specific genes were used to construct tissue-specific GPN-Star annotations, which achieved unprecedented enrichment across traits. A promising direction for future research is integrating evolutionary and functional genomics data to develop more powerful multi-modal gLMs.

Large-scale efforts to sequence and align genomes across the tree of life are accelerating [27, 28, 81, 82]. GPN-Star is well positioned to take advantage of this growth, as it scales readily to new alignments and may gain further capacity from increasingly diverse training data. In this age of rapidly expanding genomic data, we expect GPN-Star to be a valuable tool for leveraging these data to advance our understanding of genetic variation.

## Acknowledgements

This research is supported in part by National Institutes of Health grants R35-GM134922 and 3P40-OD011102-24S1 7772. In Figure 1 and Figure 6, animal icons were obtained from Flaticon (http://www.flaticon.com/).

## Methods

### Training Data

#### General workflow

The current training workflow requires:

- A multispecies whole -genome alignment (e.g., obtained with MULTIZ [21] or Cactus [22]).
- A phylogenetic tree.
- PhastCons [24] and PhyloP [25] conservation scores.

MULTIZ alignments (MAF format) were first transformed into FASTA format using maf2fasta [21]. As in GPN-MSA [29], we remove columns with gaps in the target sequence. Finally, we transform the alignment into Zarr format [83], which allows fast multi-threaded random access to genomic windows.

Similar to Benegas et al. [29], we further select training regions from the target genomes based on conservation. For every model, a sliding window equal to the model’s context length is moved across unmasked genomic regions with a stride of half the window size. We define the conservation of a window as the 75th percentile of PhastCons scores in the window. We retain the top 5% most conserved windows and a random 0.1% of the remaining windows as the training data for the model. The windows are split by chromosome into training and validation sets as detailed in Supplementary Table 1. For the selected windows, sequences on both the forward and reverse strands are included in the training data.

#### Data sources

MULTIZ alignments and associated data are readily available at the UCSC Genome Browser [84] for many of the species covered in this paper (Supplementary Table 8). The human alignments with primates and mammals [56, 85] were done with Cactus [22] and processed by Albors et al. [86]. The MULTIZ alignment for *A. thaliana* (TAIR10 reference) and 17 other Brassicales, as well as conservation scores, was downloaded from PlantRegMap [87].

#### Additional phylogenetic tree construction step for Brassicales alignment

Since no phylogenetic tree was publicly available for the species in the Brassicales alignment, we generated one with the PHAST package [88] based on fourfold degenerate sites, as described in the original publication [87]. We used the Ensembl [58] release 60 annotation. Since `msa_view`, part of the pipeline, only works with one chromosome at a time, we picked chromosome 1 arbitrarily.

~~~
msa_view../maf/tair10_multiz18way/1.maf --4d --features chr1.gff > 4d-codons.ss
msa_view 4d-codons.ss --in-format SS --out-format SS --tuple-size 1 > 4d-sites.ss
phyloFit --tree “(Carica_papaya,(Tarenaya_hassleriana,(Aethionema_arabicum,
((Arabis_alpina,(Eutrema_salsugineum,(Thellungiella_parvula,(Sisymbrium_irio,
(((Brassica_oleracea,Brassica_napus),Brassica_rapa),Raphanus_sativus))))),
((Boechera_stricta,(Camelina_sativa,(Capsella_grandiflora,Capsella_rubella))),
(Arabidopsis_thaliana,(Arabidopsis_halleri,Arabidopsis_lyrata)))))))”
--msa-format SS --out-root nonconserved-4d --EM --precision MED 4d-sites.ss
~~~

### Model Architecture

The GPN-Star model employs a novel transformer-based architecture operating on blocks of multispecies whole-genome alignments. We denote an input alignment block as 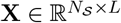, where *N*_𝒮_ is the number of species and *L* is the length of the alignment block. Each entry, or token, denoted as 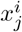, is represented as integer values encoding the nucleotide identity of the sequence of the *i*th species at position *j*.

#### Target and source sequences

The primary objective of GPN-Star is to learn a conditional distribution of nucleotides that models the evolutionary processes between sequences from different species in the alignment. Specifically, for each token in a sequence on which we want to predict the distribution, denoted as a target sequence, GPN-Star learns:

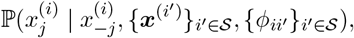

where 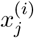 is the token at position *j* in the target sequence of the species *i*,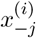 denotes the remaining tokens in the sequence *i*,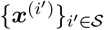 are sequences from a set of species 𝒮 that provides evolutionary context, which we call source sequences, and {*ϕ*_*ii*_*′*}_*i*′∈𝒮_ are the evolutionary distances between the target sequence and each source sequence.

- Therefore, as the first step in GPN-Star, we compile two distinct sets of sequences from each alignment block:
- Target sequences, {***x***^(*t*)^}_*t*∈𝒯_, where ***x***^(*t*)^ ∈ ℝ^*L*^, |𝒯 | = *N*_𝒯_: sequences used for loss calculation and inference. In training and inference, a subset of *N*_𝒯_ species is chosen. The first target species is designated as the primary target species, denoted as *t*_0_, which the whole genome alignment is stitched according to and thus provides contiguous genomic coordinates to anchor positional information. It is also the primary species for making inferences with the trained model.
- Source sequences, {***x***^(*s*)^}_*s*∈𝒮_, where ***x***^(*s*)^ ∈ ℝ^*L*^, |𝒮| = *N*_𝒯_: sequences of all species in the alignment. It provides the evolutionary context for inferring the distribution of the target sequences.

#### Grouping species into clades

To efficiently represent phylogenetic relationships among species, we group them into discrete clades based on pairwise phylogenetic distances. Concretely, species are partitioned into clades such that any two species belonging to distinct clades have a pairwise phylogenetic distance greater than a predefined model-specific threshold *ϕ*_clade_, which is effectively single-linkage clustering. This criterion ensures in-clade phylogenetic coherence while ensuring species in distinct clades are sufficiently distant evolutionarily. The number of clades is denoted by *N*_*C*_.

#### Embedding target and source sequences

For target sequences, each input token is directly embedded into *H*-dimension vectors:

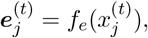

where *f*_*e*_: ℝ^*V*^ → ℝ^*H*^ is a learned embedding function and *V* is the size of nucleotide vocabulary. 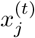 is the one-hot encoded nucleotide at position *j* of the target sequence *t*. This results in target sequence embeddings {***E***^(*t*)^}_*t*∈𝒯_, where ***E***^(*t*)^ ∈ ℝ^*L×H*^.

For embedding source sequences, the sequences of species within each clade are condensed into a single representative embedding per alignment column via an attention pooling module. For each clade *c*, tokens from member species at column *i* are combined in each attention head as follows:

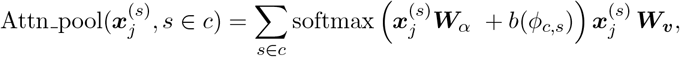

where ***W***_*α*_ ∈ ℝ^*V*×1^, ***W***_*v*_ ∈ ℝ^*V*×*D*^ are learnable weight matrices and *b*: ℝ → ℝ is the evolutionary distance encoding function (see the section Phylogeny-informed cross-attention for detailed description), and *ϕ*_*c,s*_ is the phylogenetic distance between species *s* and the Most Recent Common Ancestor (MRCA) of all species in clade *c*. 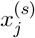 is the one-hot encoded nucleotide at position *j* of the source sequence *t*. The output embeddings across *A* attention heads are concatenated and then transformed to an *H*-dimension embedding by a linear layer, which is the final output of the module. This attention pooling procedure aggregates evolutionary signals while reducing computational complexity downstream. If there is only a single species in a clade, the attention pooling module is reduced to a single embedding module, similar to that for the target sequences. This results in clade-level source sequence embeddings {***E***^(*c*)^}_*c*∈*C*_, where ***E***^(*c*)^ ∈ ℝ^*L*×*H*^.

#### The GPN-Star encoder

GPN-Star uses an encoder-only architecture, and the encoder consists of a stack of *K* encoder blocks. Each block sequentially comprises:

1. A sequence-wise self-attention module.
2. A phylogeny-informed cross-attention module.
3. A feed-forward network.

Sequence embeddings are passed through the encoder blocks sequentially. The first and second attention modules each have *A* attention heads. Standard residual connections are applied in each module, and the embeddings are normalized with LayerNorm before each module, with dropout (probability 0.1) applied throughout. The third module is the same as in standard transformers. It is a two-layer fully-connected network that accepts the concatenated embeddings from the *A* attention heads and outputs an embedding of dimension *H*, with an intermediate layer of dimension *F* and the GELU activation. Here, we describe the first two specialized attention modules in further detail.

#### Sequence-wise self-attention

This module captures the local nucleotide context along the genome and is applied to each of the target sequences. Due to extensive rearrangements and fragmentation among the genomes of the species in the alignments, positional context is most reliably defined in the primary target species. Therefore, queries and keys in the self-attention are calculated exclusively from the embeddings of the primary target species, then the attention score is applied to all the value tensors from the embeddings of all target species. For each target sequence embedding, the output of one attention head is:

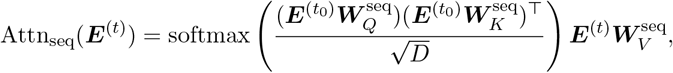

where 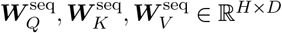 are learnable key, query and value weight matrices of the attention head, ***E***^(*t*)^ ∈ ℝ^*L*×*H*^ is the sequence embedding of the target species *t* from the previous layer, and 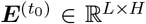 is the sequence embedding of the primary target species *t*_0_ from the previous layer. We employed the Rotary Positional Encoding (RoPE) [89] in the attention module to embed genomic positions with respect to the primary target species genome, which is omitted from the above formula for ease of description. As an additional benefit, sharing a single attention map across the *N*_𝒯_sequences significantly reduces memory footprint from *O*(*N*_𝒯_ *L*^2^) to *O*(*L*^2^), similar to another approach introduced in MSA Transformer [11].

#### Phylogeny-informed cross-attention

This module explicitly models evolutionary processes from the source sequences to the target sequences, while integrating phylogenetic information directly into attention calculations. Cross-attention is computed between target sequence embeddings (used to compute query tensors) and pooled source clade embeddings (used to compute the key and value tensors). The output of one attention head is:

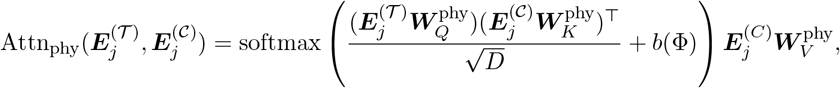

where 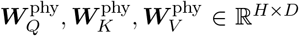 are learnable key, query and value weight matrices of the attention head, 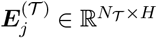 is target embeddings at the *j*th alignment column from the previous module, 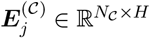 is the clade-level source embeddings at the *j*th column, and 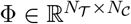 is the average phylogenetic distances between each target species and the source species in each clade. In order to encode the notion of evolutionary distance between each target and source species, we replaced the positional encoding in conventional attention modules with an evolutionary distance encoding, which is the bias term *b*(Φ) in the above formula. We adapted the Functional Interpolation for Relative Positional Encoding (FIRE) method [90] for encoding evolutionary distances:

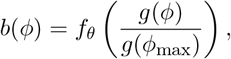

where *g*: *x* ↦ log(*cx* + 1) and *c* ∈ ℝ^+^ is a learned scaling factor, *ϕ*_max_ is the maximum distance among all species pairs in the alignment, and *f*_*θ*_: ℝ → ℝ is a two-layer perceptron 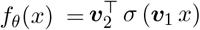, where *θ* = {***v***_1_, ***v***_2_ ∈ ℝ^*R*^} are learnable parameters and *σ* is the SiLU activation function. This evolutionary distance encoding module is shared across attention heads in the same layer.

There are two main reasons for the choice of this distance encoding scheme: first, unlike distances between positions on a sequence, the phylogenetic distances between species follow tree branching paths and are not additive along a single line. Therefore, the relative positional encoding scheme offers much convenience by simply adding bias terms to the attention score based on the pairwise distances; second, we would like the attention scores to have flexible dependencies on the evolutionary distances between species and the adapted FIRE method achieves this purpose by using a two-layer perceptron to learn a flexible mapping from the original evolutionary distance to the bias term in the attention calculation.

As mentioned, species within the same clade are evolutionarily close and have very high sequence similarity. Therefore, for the model to learn from meaningful evolutionary processes, we apply in-clade attention masks in this cross-attention module to prevent tokens in the target species from attending to the clade embeddings that the target species belongs to.

#### Output layer

Following the stack of *K* GPN-Star encoder blocks, the output layer is a standard masked language modeling head to convert hidden embeddings into per-token nucleotide probabilities:

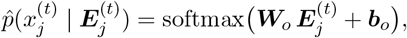

where *W*_*o*_ ∈ ℝ^*H×V*^ and *b*_*o*_ ∈ ℝ^*V*^ are the learnable weight matrix and bias term in the module, and 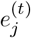 is the embedding from the final encoder block for species *t* at position *j*. The predicted nucleotide probabilities are then used to compute training loss or make inferences.

### Training Setup

The model is trained on a masked language modeling objective. For each training sample, we construct the target sequences from the alignment block as follows. The primary target species will be included in every sample. Then we select 19 target species by stratified random sampling to account for phylogenetic bias: first, we sample 19 clades out of all the clades, with replacement if the number of clades is fewer than 19 and without replacement otherwise; second, we sample one species from each selected clade. This results in 20 target species per training sample. As for source sequences, all species will be used as aforementioned.

We next apply masking to the input sequences. In each training sample, we randomly select 15% of the non-gap positions in each target sequence, ensuring that the same positions are selected in species that belong to the same clade. We denote the set of indices of the selected tokens as ℳ. Then, randomly, 90% of the selected tokens are replaced with a `[MASK]` token and the remaining 10% are left as is, resulting in a set of masked target sequences 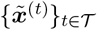. The source sequences are also masked in a way that all sequences from the same clade are masked at the same positions between the target and source sequences. The masked source sequences are denoted as 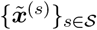. The masked sequences are used as input. And the model is trained with a weighted cross-entropy loss on the selected tokens of original target sequences after data augmentation 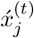 (detailed below):

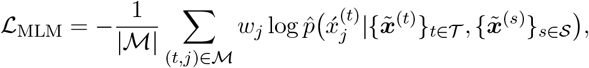

The same loss weighting scheme as Benegas *et al*. is applied [29]. The purpose is to downweight repeats and upweight conserved elements so that incorrect predictions at neutral or non-functional regions are penalized less during training. PhyloP and PhastCons are used as conservation metrics to calculate the weights. For PhastCons, we create a smoothed version, PhastCons_*M*_, that takes the maximum PhastCons score in the local 7-bp window. The loss weight of a position is defined as:

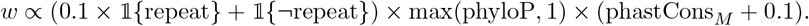

The positions in all the target sequences are defined with respect the the primary target sequence. And there is no explicit species-wise loss weighting applied. However, since we only compute loss on non-gap positions, there is an implicit downweighting of species that are more distant from the primary target, because the fraction of gap tokens tends to increase in the alignment as the species diverges further from the primary target species.

We also applied the data augmentation technique from Benegas *et al*. [29]. Before computing the loss, each position in the original target sequences is replaced by a random nucleotide with a certain probability *q*:

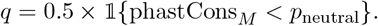

With this procedure, the high-confidence neutral sites defined by PhastCons will be randomly perturbed across training samples, so that the models learn from more diverse plausible sequences. The threshold *p*_neutral_ is set as 0.1 in all models except for the hg38 primate models, where the fraction of the defined neutral sites is much smaller than the others, and we thus raise the threshold to 0.2 in this regard.

All models are trained on a cluster of 8 NVIDIA A100 GPUs. For each alignment, we select a small fraction of chromosomes as the validation set, and the best model is selected based on the lowest validation loss over a predefined number of training steps. The full training setup of different models is described in Supplementary Table 1. The architecture hyperparameters at different model sizes are listed in Supplementary Table 9.

The hyperparameters for all the pretrained models of the same size are identical except for two: the context size and the clade-defining phylogenetic distance *ϕ*_clade_. For these two hyperparameters, we used two sets of choices for the models based on the features of the alignments. Essentially, in alignment where the species are sufficiently distant (where the maximum phylogenetic distance is greater than 1), we use a larger threshold *ϕ*_clade_ of 0.2 to define clades, and a smaller context size of 128, since these alignments are highly fragmented; in other alignments where the species are all relatively close (where the maximum phylogenetic distance is below 1), we use a smaller *ϕ*_clade_ of 0.05 to model the more nuanced evolutionary signals, and also a larger context size of 256 because there are less intense rearrangements between the genomes. The models under the first setting include: `hg38-vertebrate, mm39, galGal6, dm6`, and `ce11`. The models under the second setting are: `hg38-mammal, hg38-primate`, and `tair10`.

### Inference

When making inferences with the trained model, typically, only one target sequence is used as input, i.e., the primary target sequence. And all sequences are used as source sequences. When computing the entropy or log-likelihood ratio for a given site, we select an alignment window centered around the site with a length equal to the model training context size. The central token is masked in the input target sequence, then the model predicts nucleotide probabilities at the site. We compute the log-likelihood ratio (LLR) for a variant as follows:

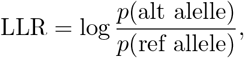

and the entropy as follows:

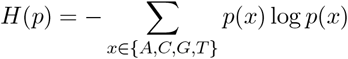

Following previous methodologies [7], we used LLR as the scoring function in most of the pathogenicity prediction and the rare variant association testing analysis. In the GWAS fine-mapped datasets, we used the absolute values of LLRs, since the studies prioritized variants based on the absolute effect sizes. For the complex trait heritability analysis, we initially started with absolute LLR, but later found that the entropy was more informative, which is surprising given that it loses allele specificity. One example where the entropy is more informative than the LLR is `rs149514089`, a putative causal variant for mean corpuscular volume and hemoglobin, located near a distal candidate enhancer [59] (Supplementary Figure 19). In this variant, both the reference and alternate allele have low frequencies in the alignment, so the LLR is close to 0.

To get the embedding of a genomic window of the target sequences, the aligned sequences of all species are used as source sequences. The input target sequence is unmasked in this case, and the hidden embeddings from the last encoder block are extracted as the output.

For the variant scores and embeddings, we average predictions from the forward and reverse strands.

### Mutation Rate Calibration

Our model is trained on genomic sequences observed in nature. These sequences are shaped by various forces in the highly complex evolutionary processes. Apart from selection, the mutation rate is also an essential determinant of the observed sequences. To more accurately estimate genome-wide selective constraints using our model, we designed a simple yet effective calibration approach to remove mutation rate variation from our model predictions. Similar to conventional phylogenetic models like PhyloP and PhastCons, the basic idea is to estimate neutral model scores at high-confidence neutral sites and then use them to adjust the model scores when making predictions.

Our procedures are described as follows. First, we gather a set of high-confidence neutral sites. For vertebrate genomes, we find ancestral repeats in the target species by lifting over the repeats in the genome of a moderately distant species and overlapping those with the repeats in the target genome. Specifically, for the human models, we lifted over the repeats from the mouse; for the mouse model, we lifted over repeats from the human; and for the chicken model, we lifted over repeats from the Zebra Finch. Next, we further select the most confidently neutral sites by selecting sites within ancestral repeats with a PhyloP score from −0.1 to 0.1, and a PhastCons score of 0. For non-vertebrate species, because the ancestral repeats provide too few sites to reliably estimate the neutral scores, we skipped the first step and directly used genome-wide neutral sites defined by PhyloP (−0.05 to 0.05) and PhastCons (0). Second, we generate the model scores at the neutral sites, including the LLR and entropy. To get baseline model scores that describe site-specific mutation rates, we bin the neutral sites by their central pentanucleotide contexts, as it is known that the central k-mer context is a major determinant of mutation rates [91]. Finally, we compute the mean model scores of variants with each pentanucleotide context, which is the baseline model score for a particular mutation rate bin determined by the pentanucleotide context and the mutation. We call them the neutral scores (i.e., LLR_*neutral*_ and *H*_*neutral*_). Then the neutral scores of the corresponding pentanucleotide bin are used to adjust the model predictions as:

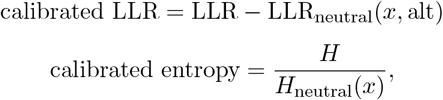

where *x* is the central pentanucleotide context of the target site.

As shown in Supplementary Figure 16 and Supplementary Figure 20, this mutation rate calibration procedure reduced the correlations between the GPN-Star scores and Roulette mutation rate estimates to a negligible level, improved constraint estimation, and the performance on most benchmarks to different extents.

### Visualizing Sequence Embeddings

We sought to visualize embeddings of a relatively balanced set of functional and non-functional genomics regions from human autosomes. We chose a resolution of 100-bp which is close to the median coding exon length in humans. We excluded all repeats from the analysis as these can use up a lot of the representational space [6] and difficult visualization of the remaining, more interesting, genomic regions. We downloaded RepeatMasker [92] repeat annotation from the UCSC Genome Browser [84]. We utilized the gene annotation from Ensembl release 113 [58] and the cCRE catalog from ENCODE SCREEN v4 [59]. For each genomic element type (e.g. lncRNA or promoter) we filtered out regions overlapping other element types, using Bioframe [93]. We defined background regions as those not overlapping exons nor cCREs. For cCREs we focused on promoters and distal enhancers. We labeled windows as “conserved” when the 75^th^ percentile of primate PhastCons [55] was 100%. For each functional element type, we sampled up to 10,000 “conserved” and 10,000 “non-conserved” regions. We sampled 20,000 background regions at random.

Model embeddings of 100-bp windows were obtained by adding flanks up to 256-bp, obtaining embeddings per position, and averaging across the center 100 positions. Embeddings from the forward and reverse strand were averaged and then standardized. UMAP [94] was run with default parameters.

### Nucleotide Dependency Analysis

Our approach is based on ref. [45]. Namely, we compute the dependency between positions *i* and *j* as:

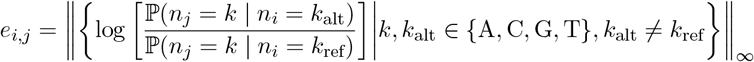

where *n*_*i*_ is the nucleotide at position *i*. No nucleotides were masked when computing this score. Additionally, we symmetrize by averaging *e*_*i,j*_ and *e*_*j,i*_.

The LDLR MPRA data [68] was downloaded in processed form from ref. [95]. We applied the most stringent filter described in the paper: variants with at least 10 barcodes and *p*-value *<* 10^−5^. To obtain a measure of effect size per position, we averaged the effect size of different alleles. The effect size for positions with no significant alleles was defined as 0. Finally, the effect size was smoothed by averaging over 5 consecutive positions.

### Evaluation Benchmarks

#### Homo sapiens

##### Datasets

- ClinVar [31]: downloaded release 20220924. The goal is to classify missense variants labeled as “Pathogenic” vs. “Benign” and the metric is the area under the receiving operating characteristic curve (AUROC).
- COSMIC [32]: downloaded data processed by ref. [29]. The goal is to classify COSMIC frequent vs. gnomAD [33] common missense variants and the metric is the area under the precision-recall curve (AUPRC).
- ProteinGym [34] downloaded the 31 human protein data in ProteinGym v0.1. We processed the dataset according to the procedures described in Benegas et al. [29].
- OMIM [41]: downloaded processed Mendelian traits dataset from TraitGym [7]. The goal is to classify pathogenic vs. common regulatory variants and the metric is the AUPRC.
- HGMD [42]: downloaded curated data from ref. [96] and processed it similarly to the Mendelian traits dataset in TraitGym [7]. The goal is to classify pathogenic vs. common regulatory variants and the metric is the AUPRC.
- GWAS fine-mapping [46, 97]: for non-coding variants, we downloaded processed complex traits dataset from TraitGym [7]. We processed coding variants from the same source [46, 97], similar to TraitGym [7] but only matching MAF. The goal is to classify variants with high vs. low posterior inclusion probability (PIP) and the metric is the AUPRC.
- gnomAD [33]. The gnomAD v3.1.2 allele frequency data were downloaded from their official website https://gnomad.broadinstitute.org/data. We follow the evaluation protocol of GPN-MSA paper [29]. Enrichment of rare (singletons) vs. common (MAF*>*5%) gnomAD variants in the tail of deleterious scores (the threshold defined by allowing a small number of false discoveries, here we use 30). We grouped certain variant consequences with small samples sizes, as follows. “splice-region” groups Ensembl categories `splice donor, splice acceptor, splice donor 5th base, splice donor region, splice region`, and `splice polypyrimidine tract`. “start-or-stop” groups Ensembl categories `start lost, stop gained or stop lost`. To reduce the number of variants for scoring, we subsampled the rare variants in each category to the same number of common variants.

We also used the Gnocchi [98] constraint scores derived from gnomAD v3 data. The precomputed genome-wide z-scores for 1kb windows that passed quality control were downloaded from the same website. The range of the scores is from -10 to 10. We downsampled the windows to 100 per score bin of length 1, resulting in 2,000 windows. In our analysis, we compared the mean GPN-Star, PhyloP, and PhastCons scores across all possible variants in each window against the Gnocchi z-score.

##### Compared models

- AlphaGenome [40]: obtained scores through the official API, following the Enformer/Borzoi protocol in TraitGym [7] but replacing the L2 aggregation across tracks with a max operation, as recommended [40].
- Borzoi [39]: downloaded precomputed scores for OMIM and GWAS fine-mapping datasets [7], and followed a similar approach for HGMD.
- CADD [95]: downloaded precomputed CADD v1.7 raw scores.
- ESM-1b [79]: log-likelihood ratio (LLR) precomputed by ref. [99] were downloaded in processed form from ref. [29].
- Enformer [38]: downloaded precomputed scores for OMIM and GWAS fine-mapping datasets [7], and followed a similar approach for HGMD. For LDSC, we downloaded precomputed scores from ref. [61], in particular tissue-agnostic feature `Enformer.Enformer_all_all` and tissue-specific features `Enformer.Enformer_<tissue>_all`.
- Evo-2 [9]: computed LLR using the 40B and 7B models with a context size of 8192-bp as in the paper.
- GPN-MSA [29]: downloaded precomputed LLR from https://huggingface.co/datasets/songlab/gpn-msa-hg38-scores.
- Nucleotide Transformer [8]: computed LLR using the 2.5B multispecies model.
- PhastCons (V): based on 100 vertebrates, downloaded from https://hgdownload.soe.ucsc.edu/goldenPath/hg38/phastCons100way/hg38.phastCons100way.bw.
- PhastCons (M): based on 470 mammals, downloaded from https://hgdownload.soe.ucsc.edu/goldenPath/hg38/phastCons470way/hg38.phastCons470way.bw
- PhastCons (P) [55]: based on 43 primates, downloaded from https://cgl.gi.ucsc.edu/data/cactus/zoonomia-2021-track-hub/hg38/phyloPPrimates.bigWig.
- PhyloP (V): based on 100 vertebrates, downloaded from https://hgdownload.soe.ucsc.edu/goldenPath/hg38/phyloP100way/hg38.phyloP100way.bw.
- PhyloP (M) [55]: based on 447 mammals, downloaded from https://hgdownload.soe.ucsc.edu/goldenPath/hg38/phyloP447way/hg38.phyloP447way.bw.
- PhyloP (P): based on 243 primates, downloaded from https://hgdownload.soe.ucsc.edu/goldenPath/hg38/phyloP447way/hg38.phyloP447wayPrimates.bw
- AlphaMissense [35]: downloaded precomputed scores for genomic variants on hg38.
- PrimateAI-3D [36]: obtained an academic license to download the scores from Illumina.
- SpeciesLM [45]: computed LLR with the model trained on animal genomes.
- PromoterAI [44]: obtained an academic license to download the scores from Illumina. We first took the absolute value of the score (to capture unsigned changes in expression rather than only over or under-expression). If a variant was predicted to affect multiple genes, we then took the maximum change in expression across all genes.
- GPN-Promoter [7]: computed LLR with the pretrained model from https://github.com/songlab-cal/gpn.
- Roulette [71]: precomputed genome-wide mutation rate estimates were downloaded from the GitHub repository https://github.com/vseplyarskiy/Roulette.
- For all scores we computed, we averaged predictions from the forward and reverse strands.

#### Mus musculus

##### Datasets

- Wild Mouse Genome Project (WMGP): population allele frequency data were derived from a set of previously published datasets [100–107] under WMGP https://andrewparkermorgan.github.io/wmgp/. Raw data were adapter- and quality-trimmed using fastp [108]. The resultant reads were aligned to a minigraph-cactus [109] constructed pangenome composed of the GRCm39 mouse reference and PacBio Continuous Long Read (CLR) assemblies of seven additional founder genomes of the Collaborative Cross mapping panel, plus PacBio CLR assemblies of BALB/cByJ, BALB/cJ, C3H/HeJ, C3H/HeOuJ, C57BL/6NJ, DBA/2J, PWD/PhJ [110] with VG giraffe [111]. The resultant alignments were surjected to the GRCm39 coordinate space in BAM format using VG [112]. Whole-genome variants were identified using DeepVariant [113] with the WGS model for each sample. Individual gVCF files were merged, and joint variant calling was conducted using GLnexus [114]. In our enrichment analysis, we defined singletons as rare variants and variants with allele frequencies above 0.2 as common variants. The molecular consequences were annotated following the same procedure as in the gnomAD dataset. To reduce the number of variants for scoring, we also followed the same procedure to subsample the rare variants.
- Mouse Mutant Resource Database (MMRdb): curated pathogenic variants were gathered from the latest MMrdb. MMRdb includes variant calls from more than 300 laboratory mouse strains with spontaneous disease phenotypes that exhibit Mendelian inheritance under cultivation at the Jackson Laboratory. A set of criteria for identifying likely variants associated with Mendelian disease phenotypes previously described in Fairfield *et al*. [115], based upon analysis of both exome and whole genome data sets, includes the following: variants will be rare in the database (*<*3%), the observed allele count in the sample will match the expected genotype (one allele for heterozygotes and two for homozygotes), and the chromosomal location of the variant will be consistent with existing linkage data. A subset of variants was derived from an earlier version of MMRdb [116], while the remainder was derived from a v.2.0.0 of the MMRdb, which can be accessed at https://mmrdb.jax.org/. All variants in the dataset were recorded in the GRCm38 reference genome coordinate space. In order to be compatible with the GRCm39 reference space used in the present model, we used the UCSC liftOver tool and associated chain files [117]. We filtered out variants that are not SNVs or with reference alleles different from the GRCm39 reference genome. We only retained variants in the following categories where there is a sufficient number of variants: missense, synonymous, nonsense, 3’ UTR. 5’ UTR and splicing. This curated dataset comprises 470 pathogenic or putatively pathogenic variants in total. Of the total dataset, 89 pathogenic variants have been confirmed via PCR validation (58 in Fairfield et al. 2015; 31 in version 2.0.0 of MMRdb), while the remaining 381 variants are classified as putatively pathogenic. We used the common variants from WMGP in the corresponding categories as the negative control in the benchmark.

##### Compared models

- PhastCons: based on 35 vertebrates, downloaded from https://hgdownload.soe.ucsc.edu/goldenPath/mm39/phastCons35way/mm39.phastCons35way.bw.
- PhyloP: based on 35 vertebrates, downloaded from https://hgdownload.soe.ucsc.edu/goldenPath/mm39/phyloP35way/mm39.phyloP35way.bw.

#### Drosophila melanogaster

##### Datasets

- Drosophila Evolution in Space and Time (DEST) [118]: downloaded allele frequency data from https://berglandlab.pods.uvarc.io/vcf/dest.all.PoolSNP.001.50.24Aug2024.ann.vcf.gz. In our enrichment analysis, we defined variants with allele frequency below 0.002 as rare variants and those with allele frequency above 0.2 as common variants, as there are no singletons in the dataset. We applied the same procedure as in the gnomAD dataset to annotate the molecular consequences of the variants.
- Flybase [75]: gathered experimentally validated lethal mutations from FlyBase https://flybase.org/ (FB2025 02, released April 17, 2025). We queried the database with the keywords “lethal”, “point mutation” and “na change” to obtain annotated SNVs that lead to the lethal phenotypes. We found the vast majority (*>*99%) of the variants are either missense, nonsense, or splicing; we therefore restricted to these three categories. After matching the reference allele with the dm6 reference genome, we got 2,929 lethal variants in total. We used the common variants from DEST in the corresponding categories as the negative control in the benchmark.

##### Compared models

- PhastCons: based on 124 insects, downloaded from https://hgdownload.soe.ucsc.edu/goldenPath/dm6/phastCons124way/dm6.phastCons124way.bw.
- PhyloP: based on 124 insects, downloaded from https://hgdownload.soe.ucsc.edu/goldenPath/dm6/phyloP124way/dm6.phyloP124way.bw.

#### Caenorhabditis elegans

##### Datasets

- Caenorhabditis elegans Natural Diversity Resource (CaeNDR) [119]: downloaded allele frequency data from https://storage.googleapis.com/caendr-site-public-bucket/dataset_release/c_elegans/20231213/variation/WI.20231213.hard-filter.isotype.vcf.gz. In our enrichment analysis, we defined variants with allele count 2 (one homozygous strain) as rare variants and those with allele frequency above 0.2 as common variants. We applied the same procedure as in the gnomAD dataset to annotate the molecular consequences of the variants.
- Experiment-validated lethal variants: obtained from the Supplementary data of Qin *et al*. [76]. We used the 72 SNVs in this dataset. We used the common variants from CaeNDR in the corresponding categories as the negative control in the benchmark.

##### Compared models

- PhastCons: based on 135 nematodes, downloaded from https://hgdownload.soe.ucsc.edu/goldenPath/ce11/phastCons135way/ce11.phastCons135way.bw.
- PhyloP: based on 135 nematodes, downloaded from https://hgdownload.soe.ucsc.edu/goldenPath/ce11/phyloP135way/ce11.phyloP135way.bw.

#### Gallus gallus

##### Datasets

- Galbase [120]: downloaded allele frequency data from http://animal.omics.pro/code/source/download/Chicken/variation/GRCg6a_SNPs.anno.tab.gz. In our enrichment analysis, we defined variants with allele frequency 0.001 (the minimum value in the dataset) as rare variants and those with allele frequency above 0.2 as common variants. We applied the same procedure as in the gnomAD dataset to annotate the molecular consequences of the variants.

##### Compared models

- PhastCons: based on 77 vertebrates, downloaded from https://hgdownload.soe.ucsc.edu/goldenPath/galGal6/phastCons77way/galGal6.phastCons77way.bw.
- PhyloP: based on 77 vertebrates, downloaded from https://hgdownload.soe.ucsc.edu/goldenPath/galGal6/phyloP77way/galGal6.phyloP77way.bw.

#### Arabidopsis thaliana

##### Datasets

- 1001 Genome Project [121]: downloaded allele frequency data processed in GPN paper [6] from https://huggingface.co/datasets/gonzalobenegas/processed-data-arabidopsis/resolve/main/variants/all/variants.parquet. In our enrichment analysis, we singletons as rare variants and those with allele frequency above 0.2 as common variants. We applied the same procedure as in the gnomAD dataset to annotate the molecular consequences of the variants.

##### Compared models

- PhastCons: based on 18 Brassicaceae, downloaded from PlantRegMap [87] http://plantregmap.gao-lab.org/download_ftp.php?filepath=08-download/Arabidopsis_thaliana/sequence_conservation/Ath_PhastCons.bedGraph.gz.
- PhyloP: based on 18 Brassicaceae, downloaded from from PlantRegMap [87] http://plantregmap.gao-lab.org/download_ftp.php?filepath=08-download/Arabidopsis_thaliana/sequence_conservation/Ath_PhyloP.bedGraph.gz.

### Heritability Enrichment Analysis

#### Background on stratified LD score regression (S-LDSC)

Following is a basic introduction to S-LDSC [52].

##### Notation

We begin by introducing some notation:

- *N* – GWAS sample size (individuals).
- *M* – number of SNPs analysed.
- **X** ∈ ℝ^*N×M*^ – standardised genotype matrix.
- **y** ∈ ℝ^*N*^ – standardised phenotype.
- ***β*** ∈ ℝ^*M*^ – true additive effect (with *β*_*j*_ denoting the effect of SNP *j*).
- *a*_*jc*_ – value of annotation *c* at SNP *j* (binary, continuous or probabilistic).
- *τ*_*c*_ – per-SNP contribution of one *unit* of annotation *c* to heritability, conditional on all other annotations.
- ***ε*** – environmental residual vector with independent 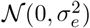 entries.

##### Model

Under the additive polygenic assumption the phenotype can be written as:

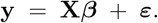

We assume that each SNP effect is an independent draw from a *zero-mean* Gaussian distribution whose variance depends linearly on the functional annotations:

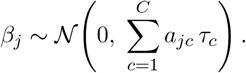

The total SNP heritability follows as:

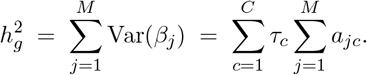

##### Inference

Parameter estimates 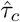 are obtained by regressing genome-wide association chi-square statistics on the stratified LD scores using (generalised) least squares. The main statistical difficulty is the strong covariance among SNP-level statistics induced by linkage disequilibrium; S-LDSC sidesteps this by collapsing LD into per-SNP scores and by computing robust standard errors with a block jackknife over genomic windows.

##### Outputs

For each annotation *c*, S-LDSC reports two *estimated* statistics that summarise its relationship with heritability:

- Coefficient 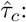 The estimated conditional per-SNP contribution to heritability associated with a one-unit increase in the annotation, obtained while adjusting for all other annotations. A scale-free version of the coefficient [122],

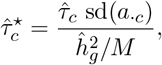

represents the proportionate change in per-SNP heritability produced by a one-standard-deviation increase in the annotation. The normalization by the mean per-SNP heritability facilitates comparison across traits.
- Enrichment (binary annotations only):

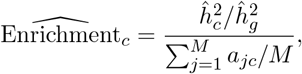

where 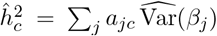 is the heritability at SNPs bearing the annotation (including contributions mediated through any overlap with other annotations). Enrichment compares the *proportion of heritability* to the *proportion of SNPs*; it is defined when *a*_*jc*_ ∈ {0, 1}.

#### S-LDSC analysis

Reference files, baseline annotations and summary statistics were downloaded from ref. [122]. We used `baselineLD_v2.2` as baseline annotations. Of the 107 independent traits with summary statistics, we kept the 106 that were available without restrictions. To evaluate a model, we ran S-LDSC including the model score and also the baseline annotations. Running S-LDSC requires computing scores for around 10M variants, prohibiting this analysis for all but the most scalable models. To be precise, S-LDSC uses 9 997 231 reference variants with allele count ≥ 5 (MAF ≥ 0.52%) in 479 European individuals from 1000 Genomes Phase 3 [123]. Of these, 5 961 159 are common (MAF ≥ 5%) and have a central role when estimating heritability (please see [52] for further details). We binarized scores based on a quantile threshold. Since enrichment is only calculated on common variants, only these were used to determine the threshold so that all model annotations have the same proportion of variants. We perform a random-effects meta-analysis of enrichments and coefficients across traits. To assess significance, we perform a one-sided Wald test. The tissue-specific analysis reuses annotations from LDSC-SEG [62] (gs://broad-alkesgroup-public-requester-pays/LDSCORE/LDSC_SEG_ldscores/Multi_tissue_gene_expr_1000Gv3_ldscores.tgz). In this study, regions specific to each GTEx [73] tissue were defined by first finding the top 10% most differentially over-expressed genes in that tissue and then adding a 100 kb window around the genes. To better match the Enformer tissue aggregation [61], we grouped certain tissues by taking the union over their windows (Supplementary Table 10). Finally, tissue-specific GPN-Star scores where defined by only picking top variants within tissue-specific regions. When comparing tissue-agnostic and tissue-specific GPN-Star annotations, we made sure they both contained the same number of variants. One thing to note is that PhastCons (P) used a much smaller alignment (43 primates) compared to GPN-Star (P) presented in the results (243 primates). We confirmed the advantage of GPN-Star (P) is not mainly driven by the increased number of species by retraining it with only the 36 intersecting species between the two sets (Supplementary Figure 21).

### Rare Variant Association Testing with DeepRVAT

Benchmarking of DeepRVAT with the published annotation set and enhanced with GPN-Star was carried out according to the same benchmarking procedure as for Fig. 4a in [49]. We describe the details of this in brief here.

UK Biobank WES data was processed and quality-controlled according to the procedure of the “UKBB 200k unrelated European ancestry dataset” from [49]. This dataset was derived from the UK Biobank 200K interim WES by including only individuals of European genetic ancestry and excluding those related to third degree or closer, resulting in a cohort of 161,822 individuals. This restriction was done for benchmarking in order to remove confounding effects of population stratification. Sequencing data processing, quality control, and annotation was carried out following [49], and variants with MAF *<* 1% were retained for training and variants with MAF *<* 0.1% were retained for association testing.

DeepRVAT models (package v1.1) with and without GPN-Star variant annotations were trained using stochastic parameter initialization for three different random seeds and the same 21 training phenotypes as in [49]. Burden testing was carried out as in Fig. 4a of [49], using REGENIE with DeepRVAT gene scores on the same 34 quantitative phenotypes (21 used during training and 13 not).

DeepRVAT discoveries were those with a Bonferroni-corrected *p*-value *<* 0.05. Replication was carried out by comparing significant gene-phenotype associations to reported discoveries from two studies [50, 51] on larger WES cohorts from UK Biobank (394,841 and 454,787 individuals, respectively).

## Data Availability

The pretrained models, training datasets and benchmark datasets are available on Hugging Face (https://huggingface.co/collections/songlab/gpn-star-68c0c055acc2ee51d5c4f129). Predictions for all possible single nucleotide variants in the human genome and the five model organisms will be made publicly available upon publication.

## Code Availability

Code to train the model, run inference, and reproduce the main analyses is available on GitHub (https://github.com/songlab-cal/gpn).

## Supplementary Tables

**Supplementary Table 1:**
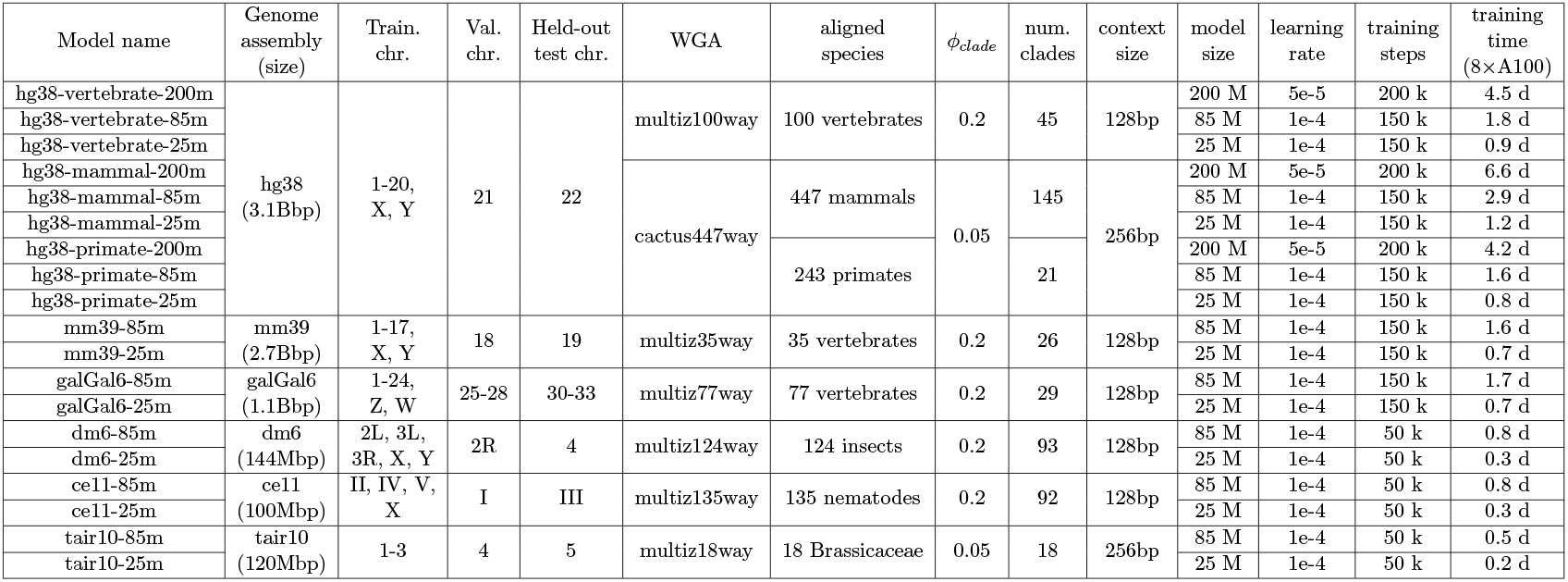
A summary of the training setups of all GPN-Star models in this work.

**Supplementary Table 2:** ENCODE SCREEN cCRE classes.

|  | TSS dist. | DNase/ATAC | H3K4me3 | H3K27ac | CTCF | TF |
| --- | --- | --- | --- | --- | --- | --- |
| promoter (PLS) | < 200 | ✓ | ✓ |  |  |  |
| proximal enhancer (pELS) | < 2 000 | ✓ |  | ✓ |  |  |
| distal enhancer (dELS) | > 2 000 | ✓ |  | ✓ |  |  |
| chromatin accesibility w/ H3K4me3 (CA-H3K4me3) |  | ✓ | ✓ | × |  |  |
| chromatin accesibility w/ CTCF (CA-CTCF) |  | ✓ | × | × | ✓ |  |
| chromatin accesibility w/ TF (CA-TF) |  | ✓ | × | × | × | ✓ |
| chromatin accesibility (CA) |  | ✓ | × | × | × | × |
| transcription factor (TF) |  | × | × | × | × | ✓ |

**Supplementary Table 3:** Consequence distribution for the top 0.1% most constrained variants according to GPN-Star (P) (common variants only). Odds ratio and *p*-value were calculated using a two-sided Fisher’s exact test. Bold: significant under a FDR *<* 5%.

| consequence | count | proportion | odds ratio |
| --- | --- | --- | --- |
| missense | 1983 | 0.33 | **170.27** |
| dELS | 1564 | 0.26 | **2.39** |
| dELS-flank | 468 | 0.08 | **0.35** |
| ncRNA | 287 | 0.05 | **2.14** |
| 3' UTR | 280 | 0.05 | **4.23** |
| pELS | 209 | 0.04 | **2.14** |
| 5' UTR | 151 | 0.03 | **9.83** |
| CA | 135 | 0.02 | 0.92 |
| intron | 99 | 0.02 | **0.07** |
| PLS | 96 | 0.02 | **7.18** |
| CA-flank | 92 | 0.02 | **0.21** |
| synonymous | 77 | 0.01 | **4.11** |
| intergenic | 67 | 0.01 | **0.05** |
| pELS-flank | 61 | 0.01 | **0.57** |
| CA-CTCF | 60 | 0.01 | 0.94 |
| CA-H3K4me3 | 49 | 8e-03 | 1.07 |
| splice-region | 40 | 7e-03 | **7.52** |
| CA-CTCF-flank | 36 | 6e-03 | **0.19** |
| splice-donor | 28 | 5e-03 | **50.38** |
| splice-donor-region | 27 | 5e-03 | **23.04** |
| TF-flank | 24 | 4e-03 | **0.14** |
| splice-donor-5th-base | 22 | 4e-03 | **52.98** |
| CA-TF | 20 | 3e-03 | 1.30 |
| splice-polypyrimidine-tract | 18 | 3e-03 | **4.19** |
| CA-H3K4me3-flank | 15 | 3e-03 | **0.14** |
| splice-acceptor | 11 | 2e-03 | **30.65** |
| CA-TF-flank | 10 | 2e-03 | **0.26** |
| stop-gained | 10 | 2e-03 | **41.91** |
| TF | 9 | 2e-03 | **0.15** |
| start-lost | 4 | 7e-04 | **68.72** |
| downstream-gene | 3 | 5e-04 | **0.15** |
| mature-miRNA | 3 | 5e-04 | **48.21** |
| stop-lost | 2 | 3e-04 | **22.64** |
| PLS-flank | 1 | 2e-04 | **0.17** |

**Supplementary Table 4:**
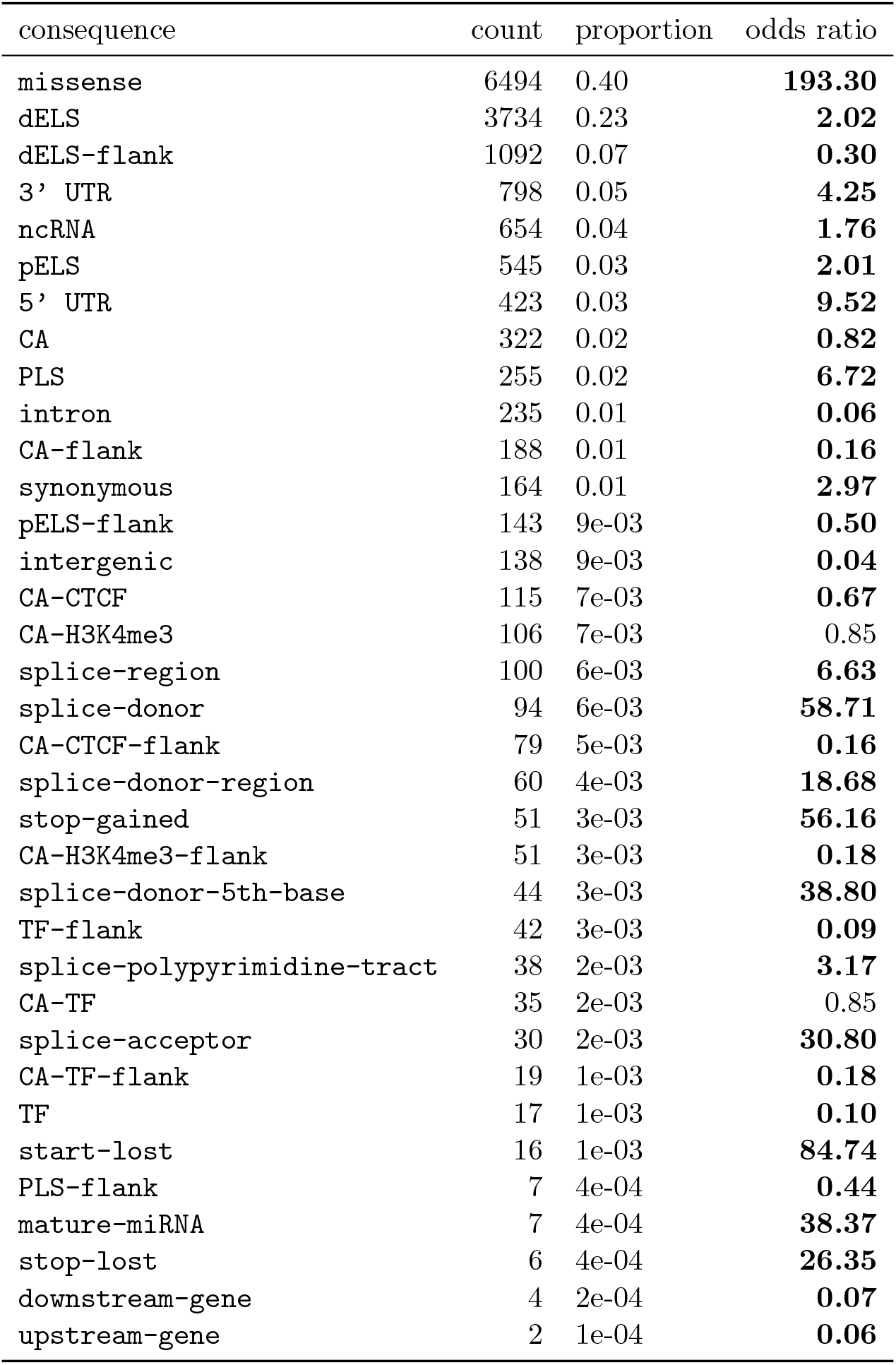
Consequence distribution for the top 0.1% most constrained variants according to GPN-Star (P) (all S-LDSC variants, common and low-frequency). Odds ratio and *p*-value were calculated using a two-sided Fisher’s exact test. Bold: significant under a FDR *<* 5%.

| consequence | count | proportion | odds ratio |
| --- | --- | --- | --- |
| missense | 6494 | 0.40 | **193.30** |
| dELS | 3734 | 0.23 | **2.02** |
| dELS-flank | 1092 | 0.07 | **0.30** |
| 3' UTR | 798 | 0.05 | **4.25** |
| ncRNA | 654 | 0.04 | **1.76** |
| pELS | 545 | 0.03 | **2.01** |
| 5' UTR | 423 | 0.03 | **9.52** |
| CA | 322 | 0.02 | **0.82** |
| PLS | 255 | 0.02 | **6.72** |
| intron | 235 | 0.01 | **0.06** |
| CA-flank | 188 | 0.01 | **0.16** |
| synonymous | 164 | 0.01 | **2.97** |
| pELS-flank | 143 | 9e-03 | **0.50** |
| intergenic | 138 | 9e-03 | **0.04** |
| CA-CTCF | 115 | 7e-03 | **0.67** |
| CA-H3K4me3 | 106 | 7e-03 | 0.85 |
| splice-region | 100 | 6e-03 | **6.63** |
| splice-donor | 94 | 6e-03 | **58.71** |
| CA-CTCF-flank | 79 | 5e-03 | **0.16** |
| splice-donor-region | 60 | 4e-03 | **18.68** |
| stop-gained | 51 | 3e-03 | **56.16** |
| CA-H3K4me3-flank | 51 | 3e-03 | **0.18** |
| splice-donor-5th-base | 44 | 3e-03 | **38.80** |
| TF-flank | 42 | 3e-03 | **0.09** |
| splice-polypyrimidine-tract | 38 | 2e-03 | **3.17** |
| CA-TF | 35 | 2e-03 | 0.85 |
| splice-acceptor | 30 | 2e-03 | **30.80** |
| CA-TF-flank | 19 | 1e-03 | **0.18** |
| TF | 17 | 1e-03 | **0.10** |
| start-lost | 16 | 1e-03 | **84.74** |
| PLS-flank | 7 | 4e-04 | **0.44** |
| mature-miRNA | 7 | 4e-04 | **38.37** |
| stop-lost | 6 | 4e-04 | **26.35** |
| downstream-gene | 4 | 2e-04 | **0.07** |
| upstream-gene | 2 | 1e-04 | **0.06** |

**Supplementary Table 5:**
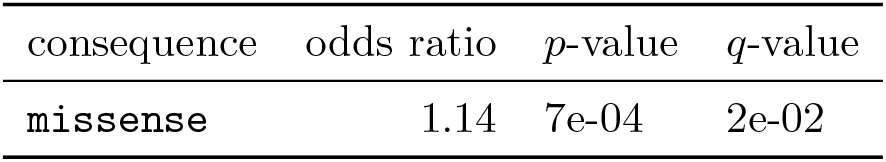
Consequences significantly enriched at the top common variants according to GPN-Star (P) vs. GPN-Star (M). consequence odds ratio *p*-value *q*-value missense 1.14 7e-04 2e-02.

**Supplementary Table 6:**
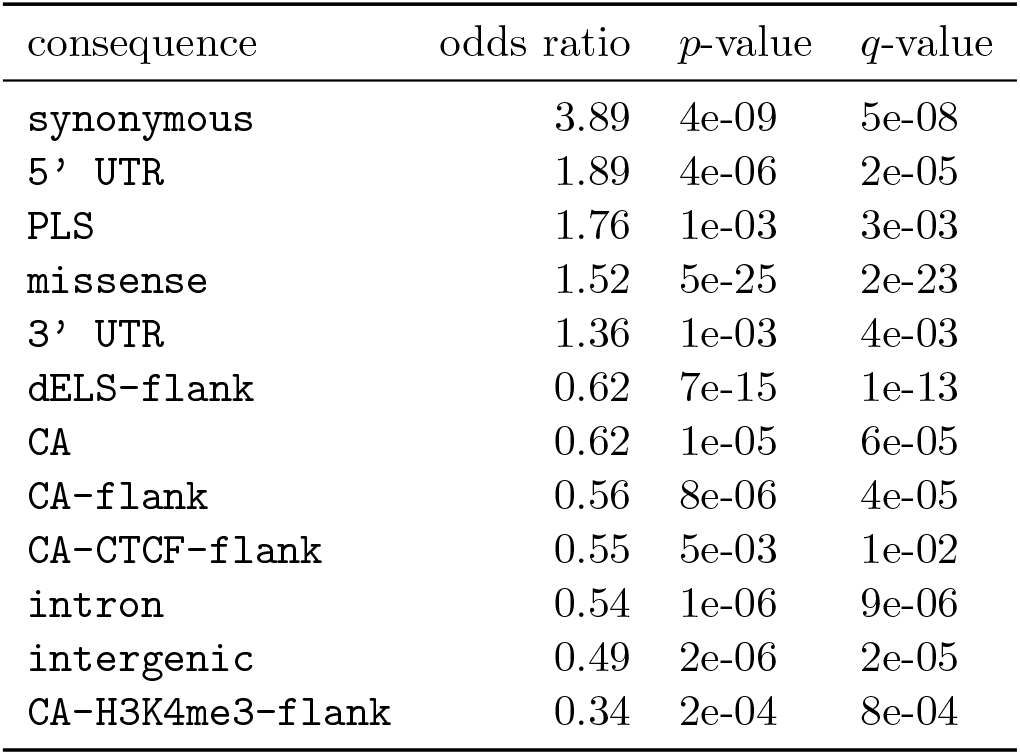
Consequences significantly enriched at the top common variants according to GPN-Star (P) vs. GPN-Star (V).

| consequence | odds ratio | *p*-value | *q*-value |
| --- | --- | --- | --- |
| **synonymous** | 3.89 | 4e-09 | 5e-08 |
| **5' UTR** | 1.89 | 4e-06 | 2e-05 |
| **PLS** | 1.76 | 1e-03 | 3e-03 |
| **missense** | 1.52 | 5e-25 | 2e-23 |
| **3' UTR** | 1.36 | 1e-03 | 4e-03 |
| **dELS-flank** | 0.62 | 7e-15 | 1e-13 |
| **CA** | 0.62 | 1e-05 | 6e-05 |
| **CA-flank** | 0.56 | 8e-06 | 4e-05 |
| **CA-CTCF-flank** | 0.55 | 5e-03 | 1e-02 |
| **intron** | 0.54 | 1e-06 | 9e-06 |
| **intergenic** | 0.49 | 2e-06 | 2e-05 |
| **CA-H3K4me3-flank** | 0.34 | 2e-04 | 8e-04 |

**Supplementary Table 7:**
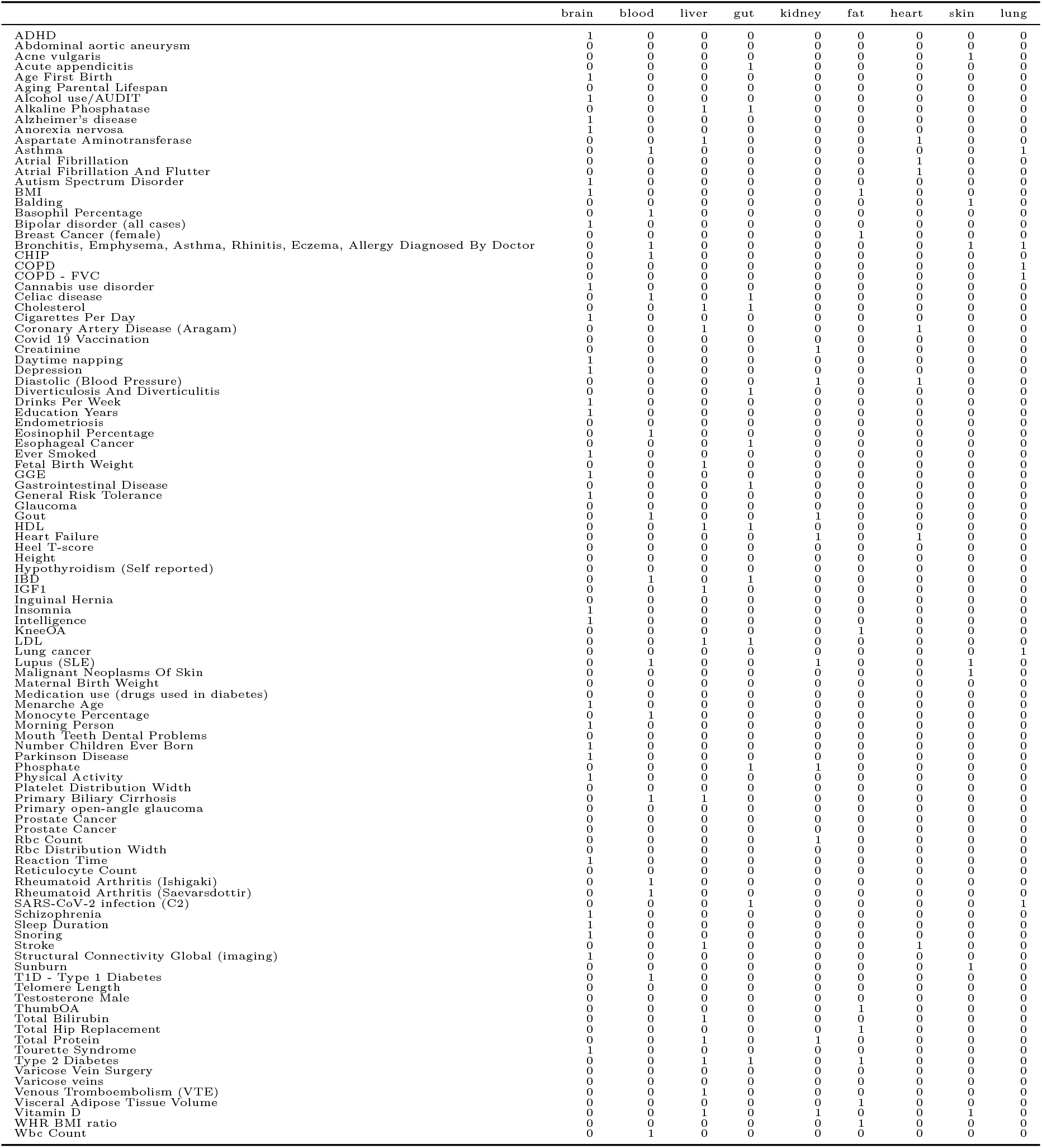
Traits and tissues we expect them to be related to.

**Supplementary Table 8:** Alignments obtained from the UCSC Genome Browser.

|  | Reference | Alignment |
| --- | --- | --- |
| *H. sapiens* | hg38 | multiz100way (100 vertebrates) |
| *M. musculus* | mm39 | multiz35way (35 vertebrates) |
| *G. gallus* | galGal6 | multiz77way (77 vertebrates) |
| *D. melanogaster* | dm6 | multiz124way (124 insects) |
| *C. elegans* | ce11 | multiz135way (135 nematodes) |

**Supplementary Table 9:** Hyperparameters of the GPN-Star architecture at different model sizes.

| Model size | 25M | 85M | 200M |
| --- | --- | --- | --- |
| Num encoder blocks (K) | 8 | 12 | 16 |
| Hidden dim (H) | 512 | 768 | 1024 |
| Intermediate size (F) | 2048 | 3072 | 4096 |
| Num self-attn heads (A) | 4 | 6 | 8 |
| Num cross-attn heads (A) | 4 | 6 | 8 |
| Attn head dim (D) | 64 | 64 | 64 |
| Dropout rate | 0.1 | 0.1 | 0.1 |

**Supplementary Table 10:** Grouping of GTEx tissues. brain blood/immune.

|  | GTEx tissue |
| --- | --- |
| brain | Brain Substantia nigra |
|  | Brain Spinal cord (cervical c-1) |
|  | Brain Amygdala |
|  | Brain Anterior cingulate cortex (BA24) |
|  | Brain Hippocampus |
|  | Brain Hypothalamus |
|  | Brain Putamen (basal ganglia) |
|  | Brain Cerebellar Hemisphere |
|  | Brain Frontal Cortex (BA9) |
|  | Brain Nucleus accumbens (basal ganglia) |
|  | Brain Cortex |
|  | Brain Caudate (basal ganglia) |
|  | Brain Cerebellum |
| blood/immune | Spleen |
|  | Cells EBV-transformed lymphocytes |
|  | Whole Blood |
| liver | Liver |
| gut | Small Intestine Terminal Ileum |
|  | Colon Sigmoid |
|  | Esophagus Gastroesophageal Junction |
|  | Stomach |
|  | Colon Transverse |
|  | Esophagus Muscularis |
|  | Esophagus Mucosa |
| kidney | Kidney Cortex |
| fat | Adipose Visceral (Omentum) |
|  | Adipose Subcutaneous |
| heart | Heart Atrial Appendage |
|  | Heart Left Ventricle |
| skin | Skin Not Sun Exposed (Suprapubic) |
|  | Cells Transformed fibroblasts |
|  | Skin Sun Exposed (Lower leg) |
| lung | Lung |

## Supplementary Figures

**Supplementary Figure 1:**
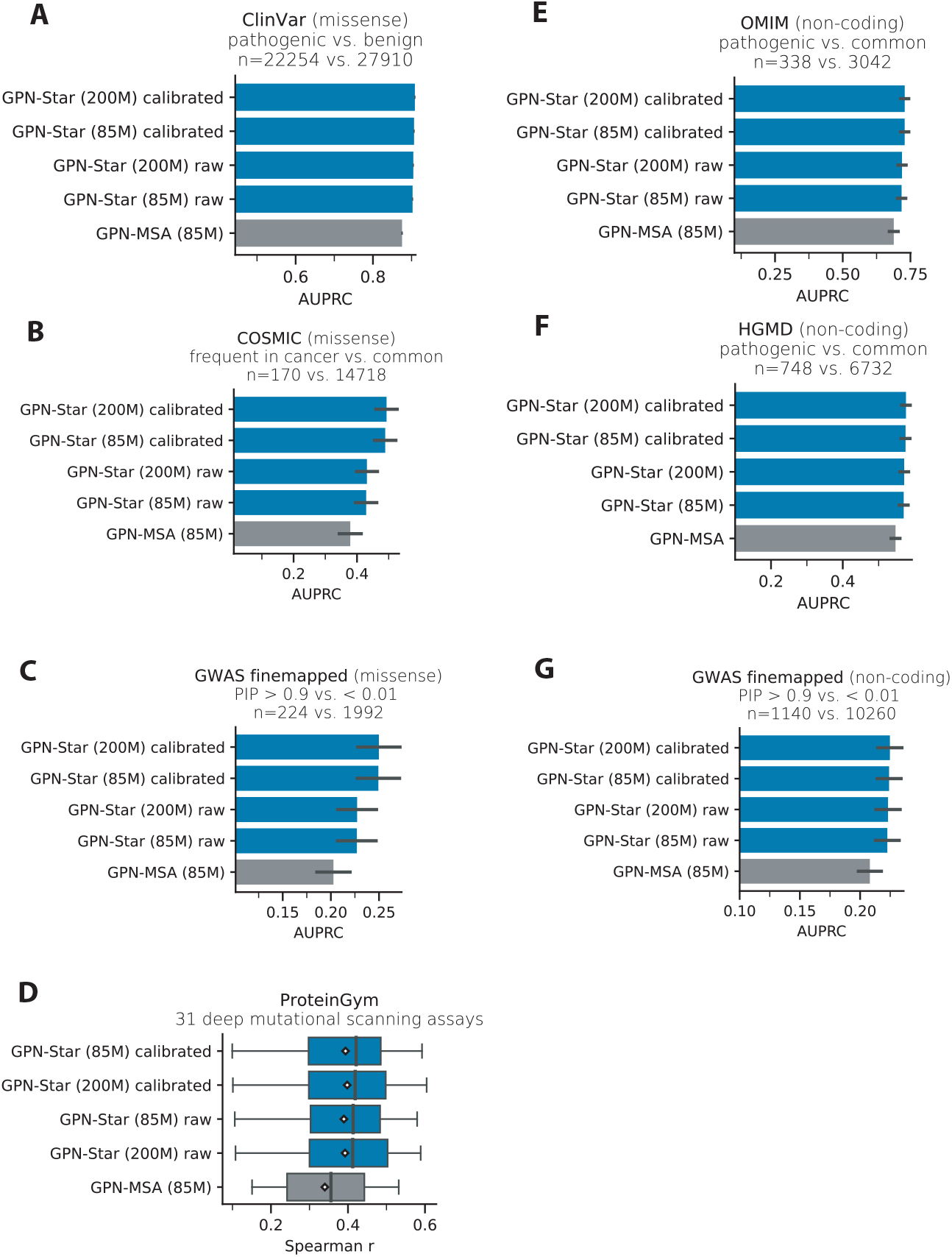
Head-to-head comparison with GPN-MSA on variant effect prediction. The benchmarks are the same as in Figure 2. For fair comparison, we showed the performance of GPN-Star with the same model size (85M) and the uncalibrated raw scores.

**Supplementary Figure 2:**
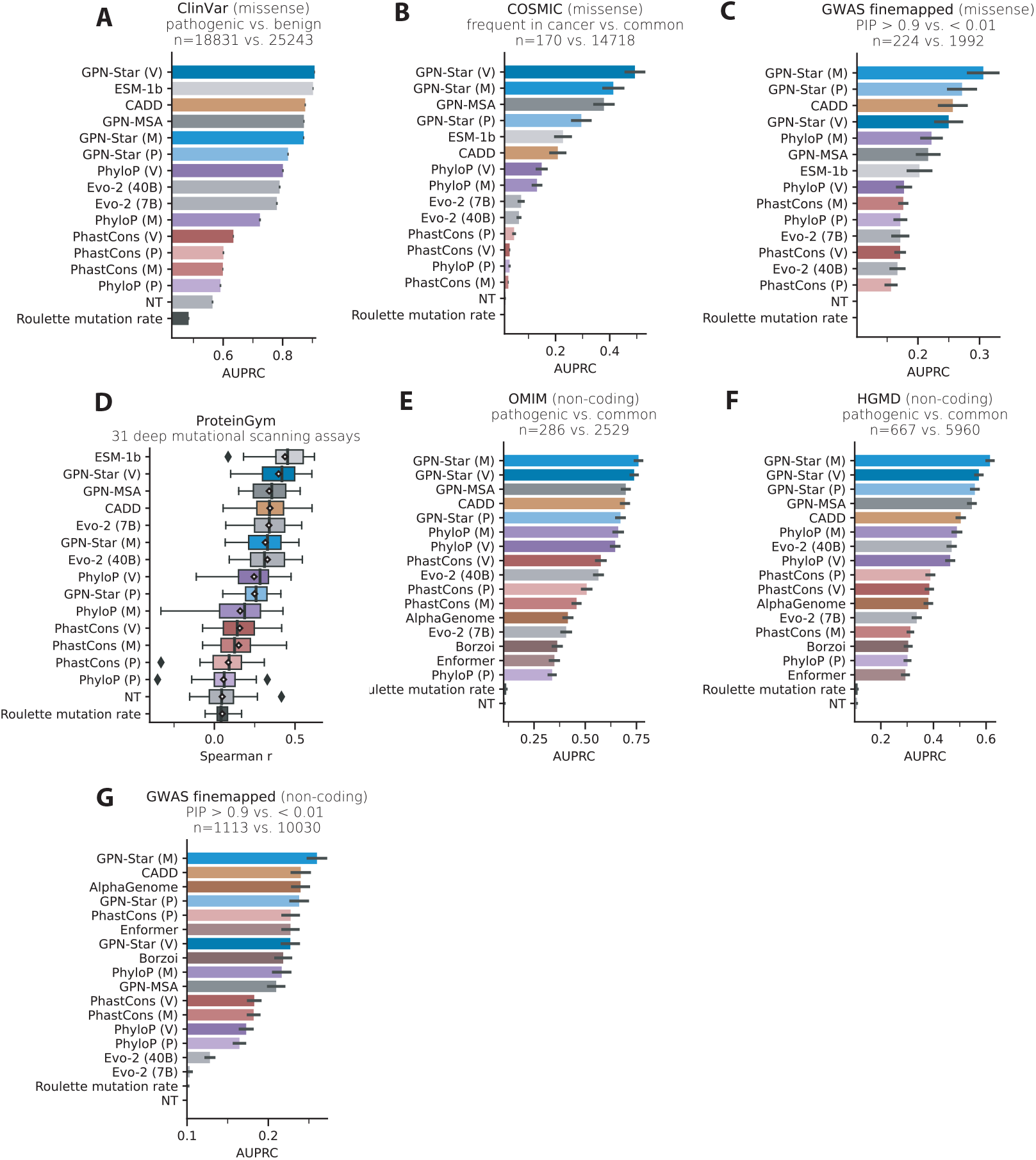
Variant effect prediction benchmarks with all competing models. The benchmarks are the same as in Figure 2, while including GPN-Star, PhyloP and PhastCons with all three evolutionary time scales, Evo-2 with two parameter sizes and Roulette mutation rate estimates in the comparison.

**Supplementary Figure 3:**
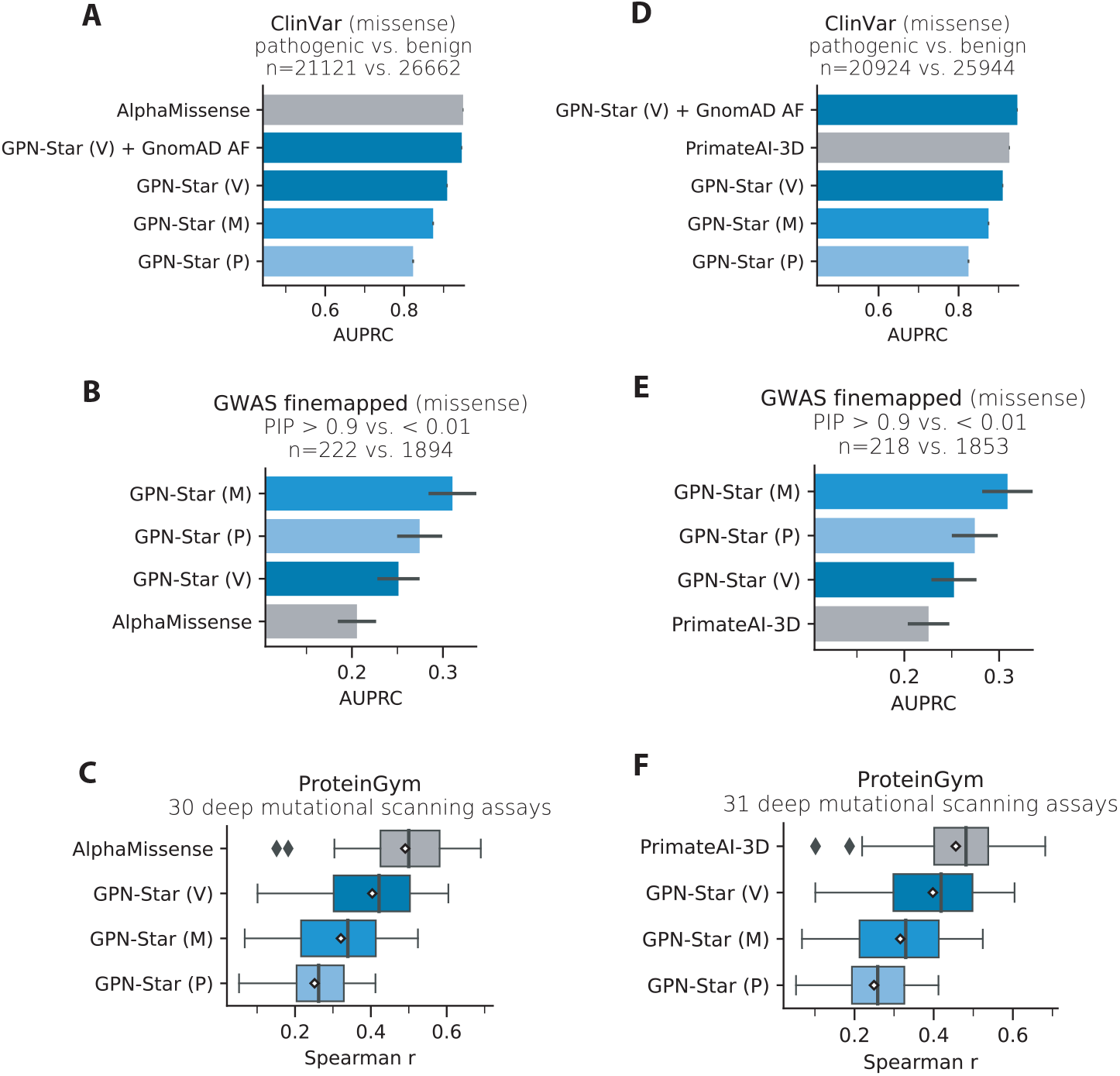
Head-to-head comparisons with AlphaMissense and PrimateAI-3D on missense variant effect prediction. This includes the coding benchmarks in Figure 2 except for the COSMIC dataset, since its negative set is entirely within the training data of the two models. The sample sizes in certain comparisons are smaller here since the precomputed scores from the two methods do not cover all the variants. In the ClinVar benchmarks, the method GPN-Star (V) + gnomAD AF is adjusting GPN-Star (V) predictions by assigning the most benign score to those variants with allele frequency *>* 2*e* − 4 in gnomAD v3. The threshold is chosen according to the definition of benign variants in AlphaMissense training data.

**Supplementary Figure 4:**
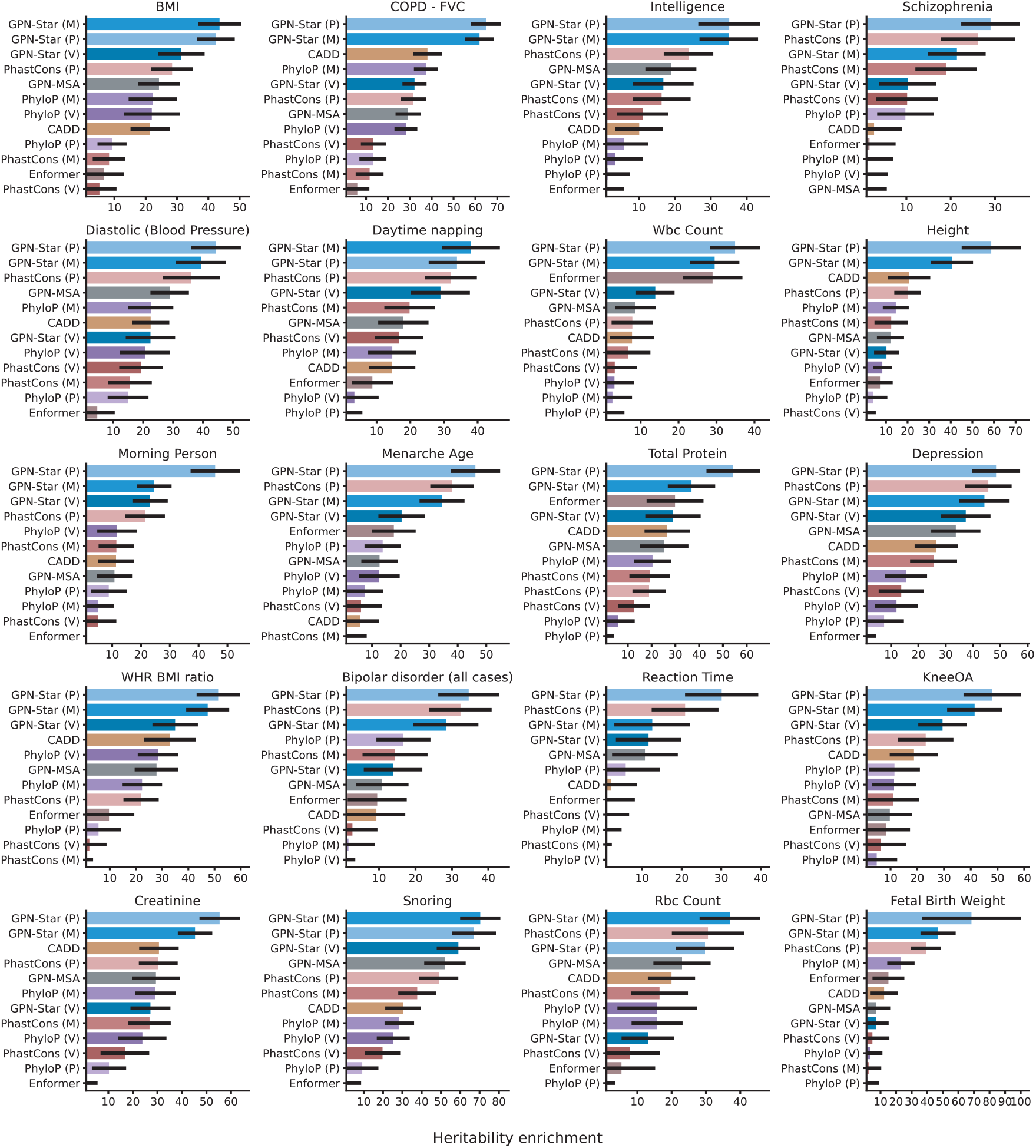
Heritability enrichment by model predictions of the 20 most heritable traits. Same as in Figure 3, here we showed the enrichment among the top 0.1% most constraint variants defined by each model.

**Supplementary Figure 5:**
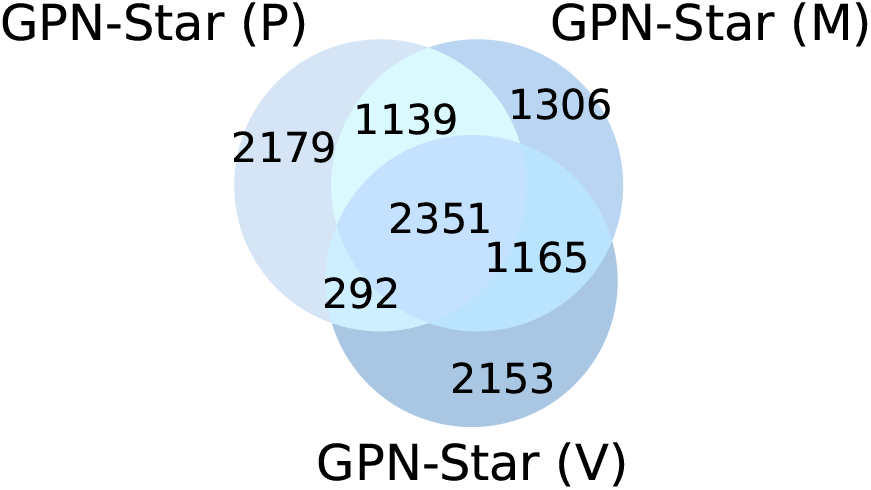
Overlap between top 0.1% most constrained variants according to the three models (common variants only).

**Supplementary Figure 6:**
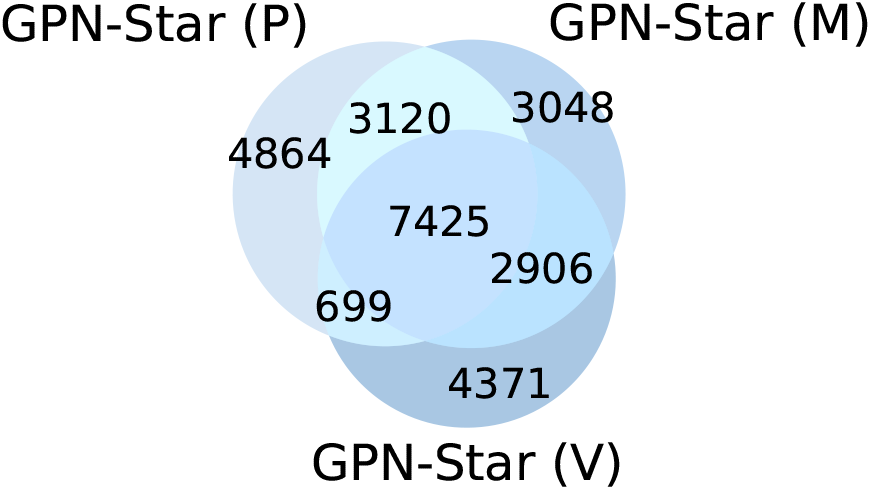
Overlap between top 0.1% most constrained variants according to the three models (all S-LDSC variants, common and low-frequency).

**Supplementary Figure 7:**
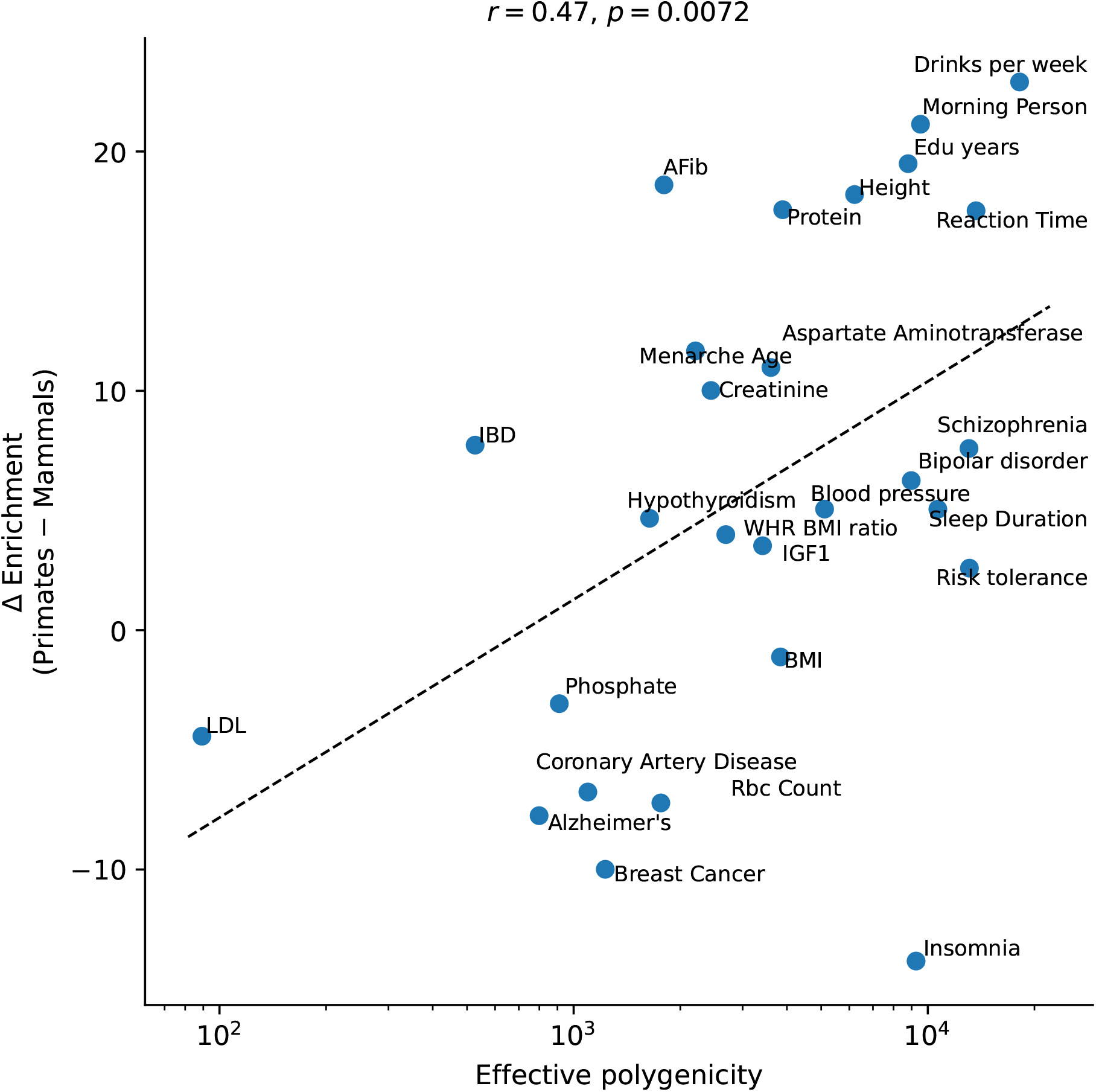
Difference in enrichment between GPN-Star (P) and (M) as a function of estimated effective polygenicity. *p*-value is one-sided. Dashed line is ordinary least squares fit.

**Supplementary Figure 8:**
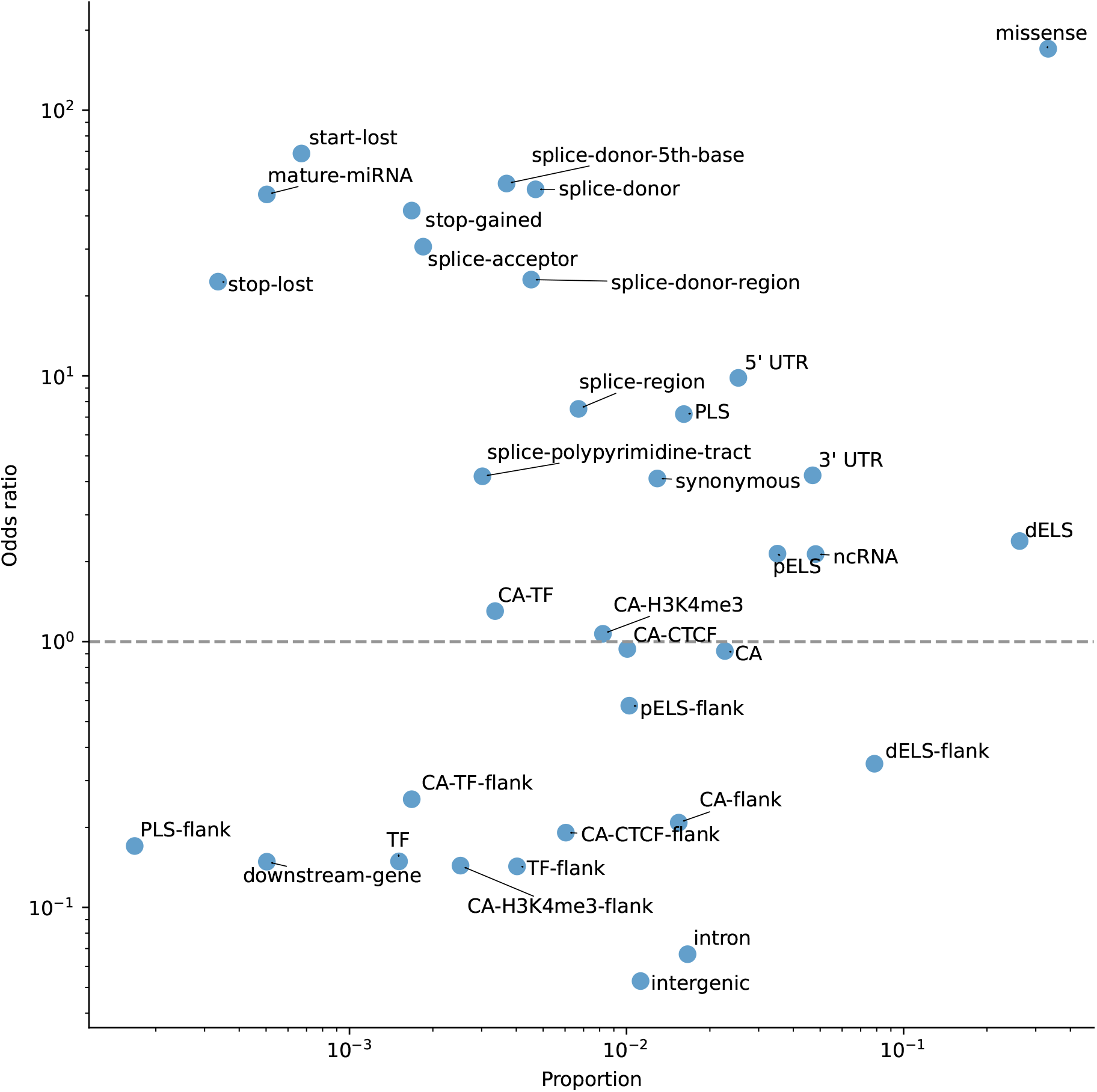
Proportion of variant consequence among the top 0.1% most constrained variants according to GPN-Star (P) and odds ratio with respect to the 99.9% least constrained variants (common variants only).

**Supplementary Figure 9:**
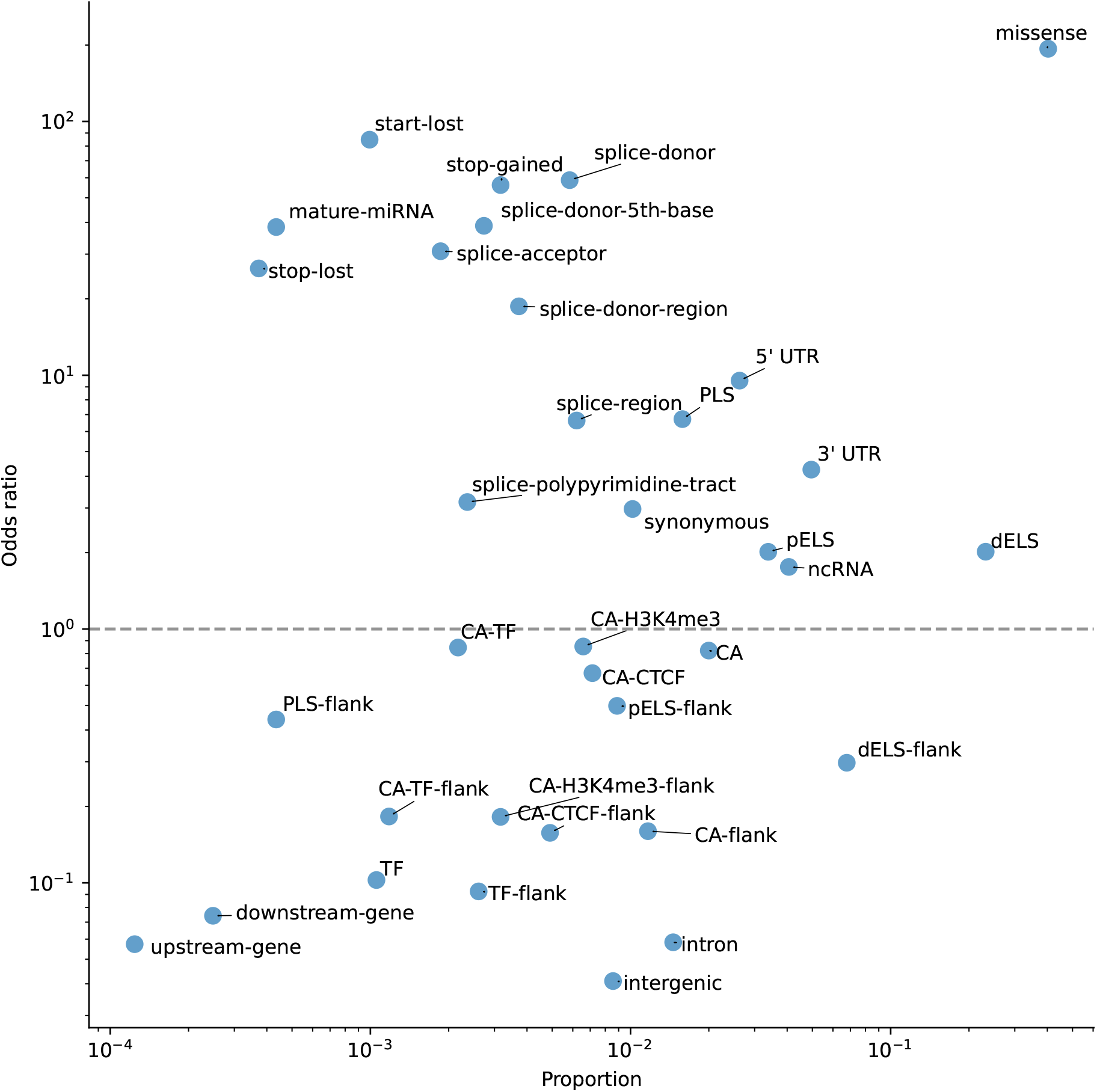
Proportion of variant consequence among the top 0.1% most constrained variants according to GPN-Star (P) and odds ratio with respect to the 99.9% least constrained variants (all S-LDSC variants, common and low-frequency.

**Supplementary Figure 10:**
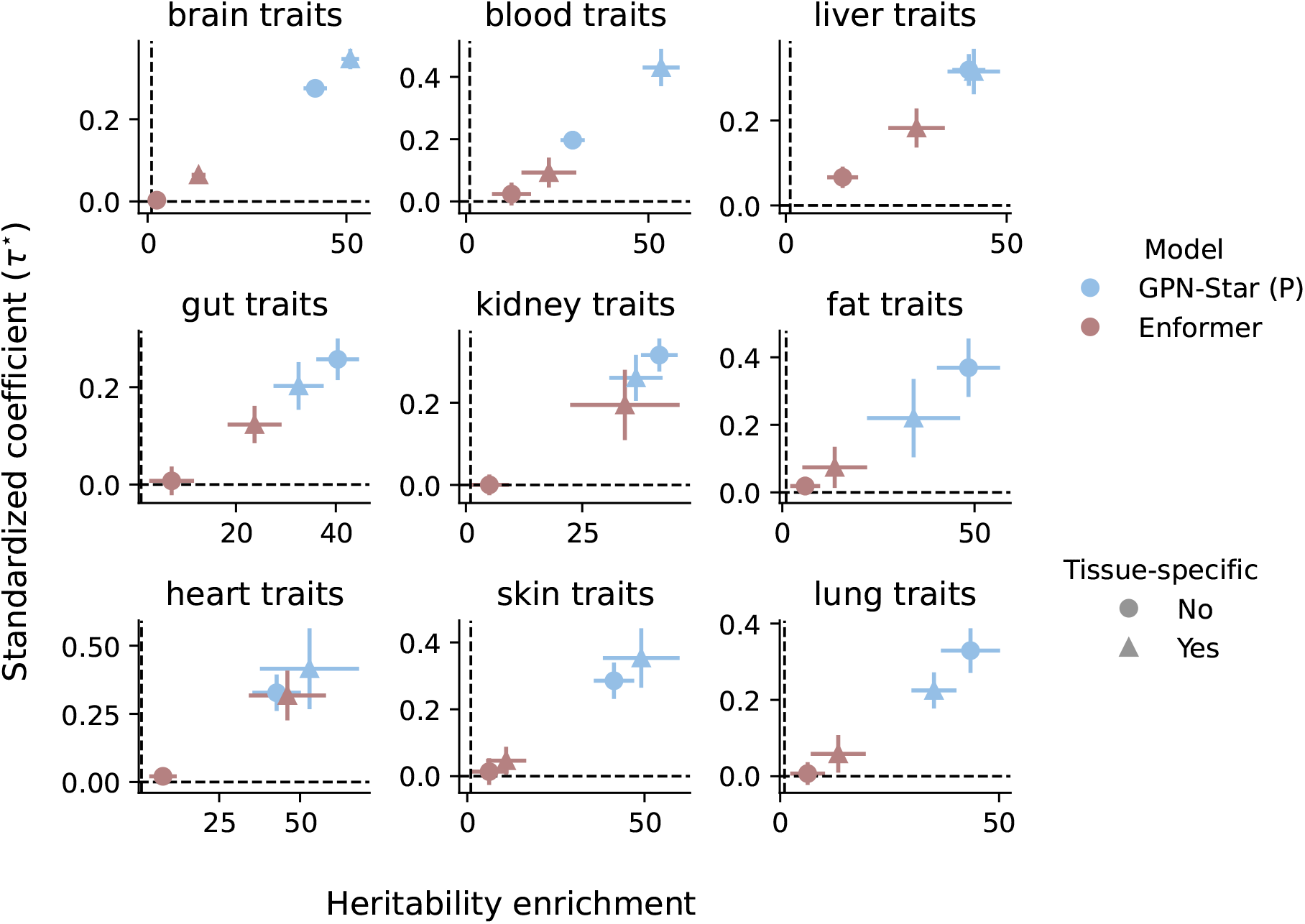
Heritability enrichment and standardized coefficient for tissue-specific analysis.

**Supplementary Figure 11:**
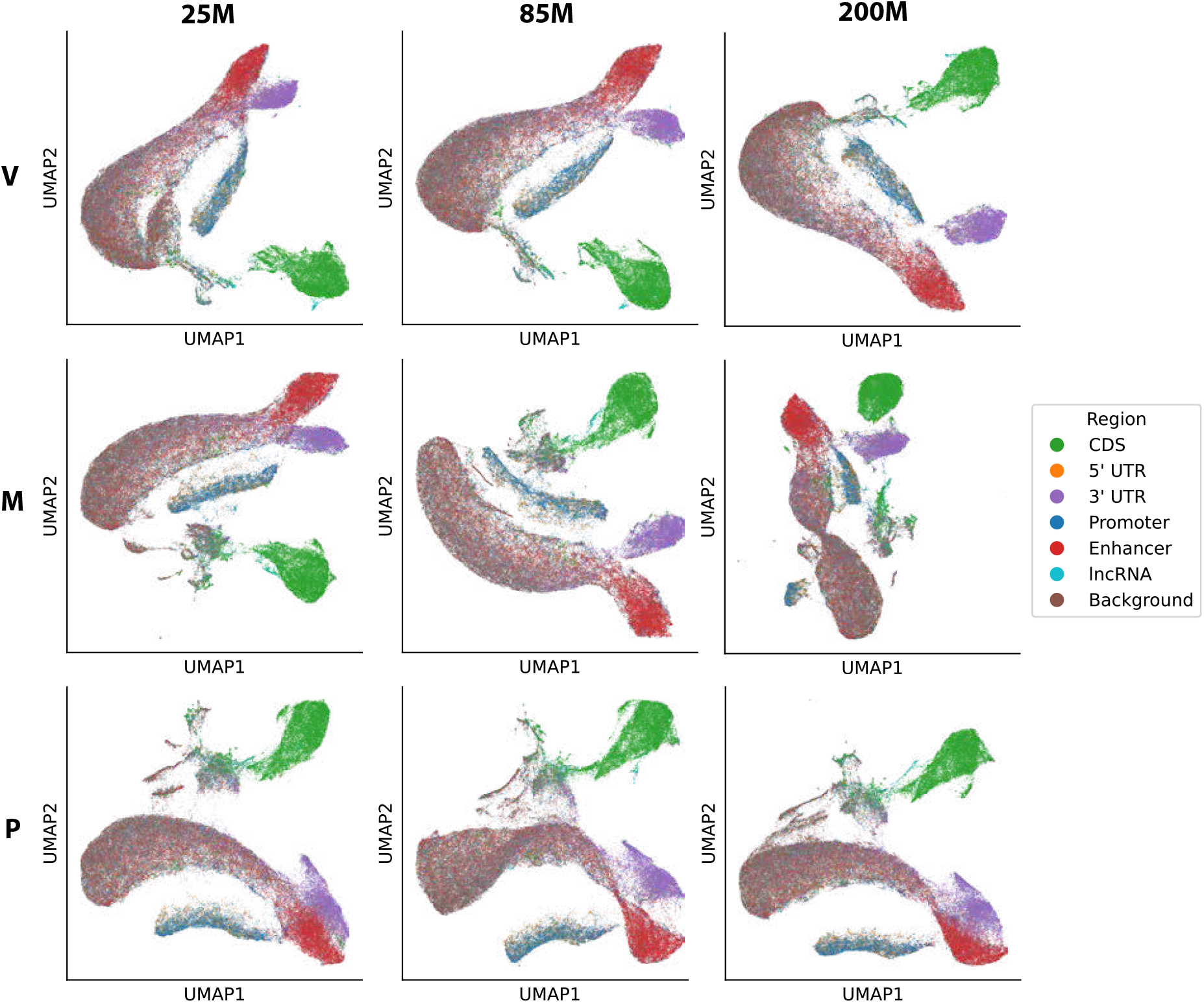
Visualization of window embeddings (all models). The rows represent models trained on different evolutionary timescales (V: vertebrates, M: mammals, P: primates) and the columns represent different model sizes (number of parameters).

**Supplementary Figure 12:**
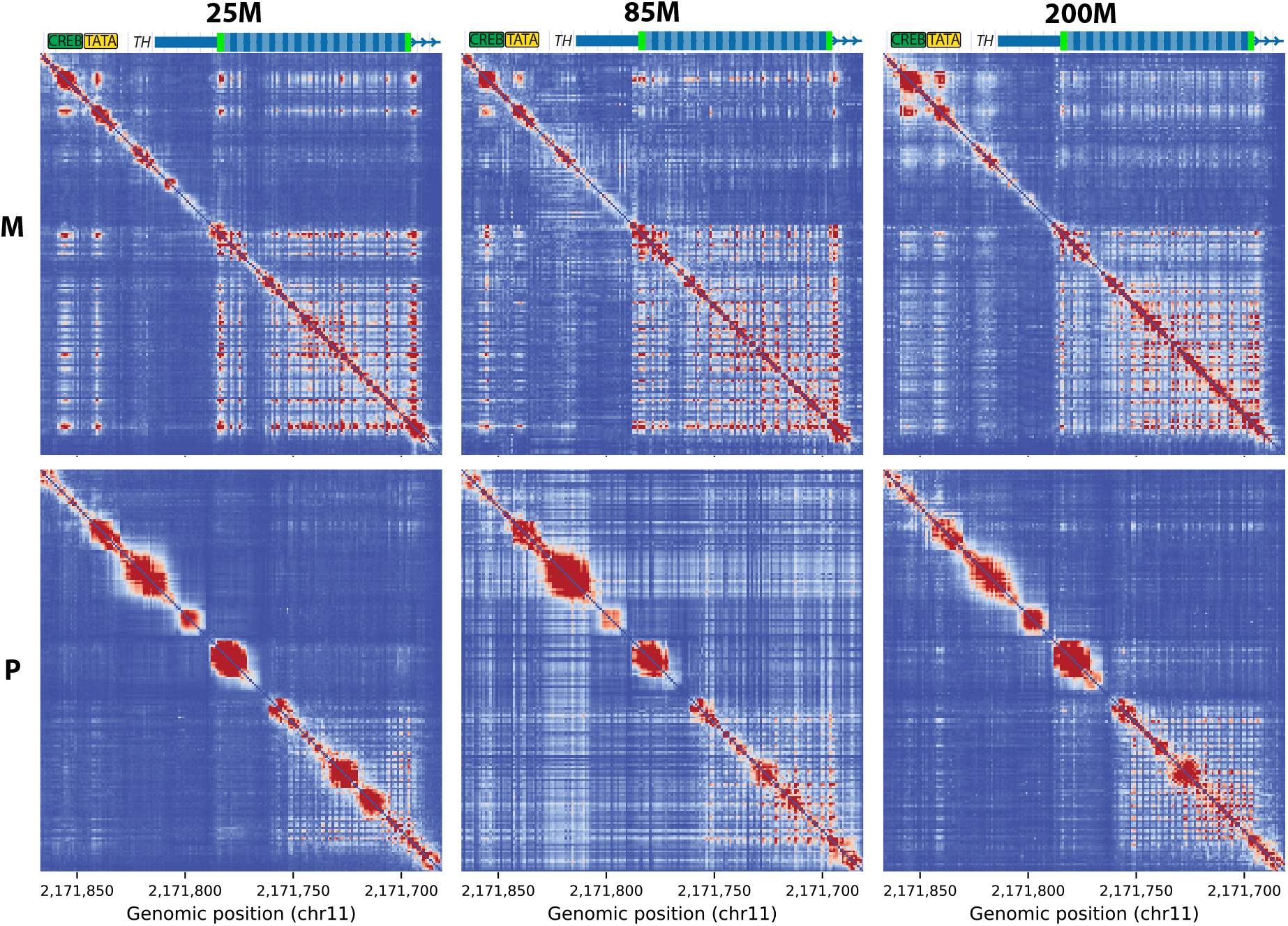
Nucleotide dependency map in *TH* promoter (vertebrate models excluded as these region exceeds its context size). The rows represent models trained on different evolutionary timescales (V: vertebrates, M: mammals, P: primates) and the columns represent different model sizes (number of parameters).

**Supplementary Figure 13:**
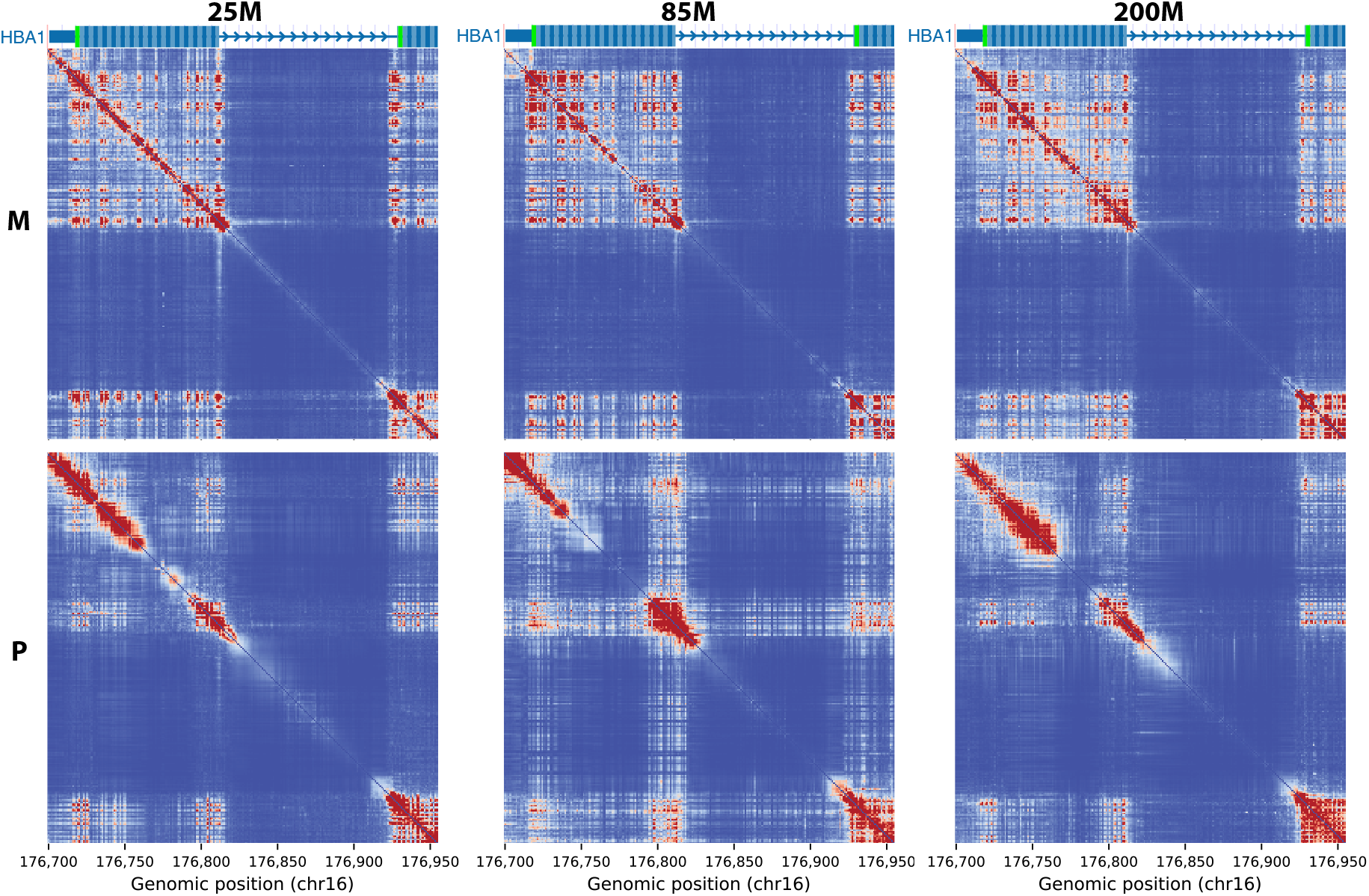
Nucleotide dependency map around the first *HBA1* exons (vertebrate models excluded as these region exceeds its context size). The rows represent models trained on different evolutionary timescales (V: vertebrates, M: mammals, P: primates) and the columns represent different model sizes (number of parameters).

**Supplementary Figure 14:**
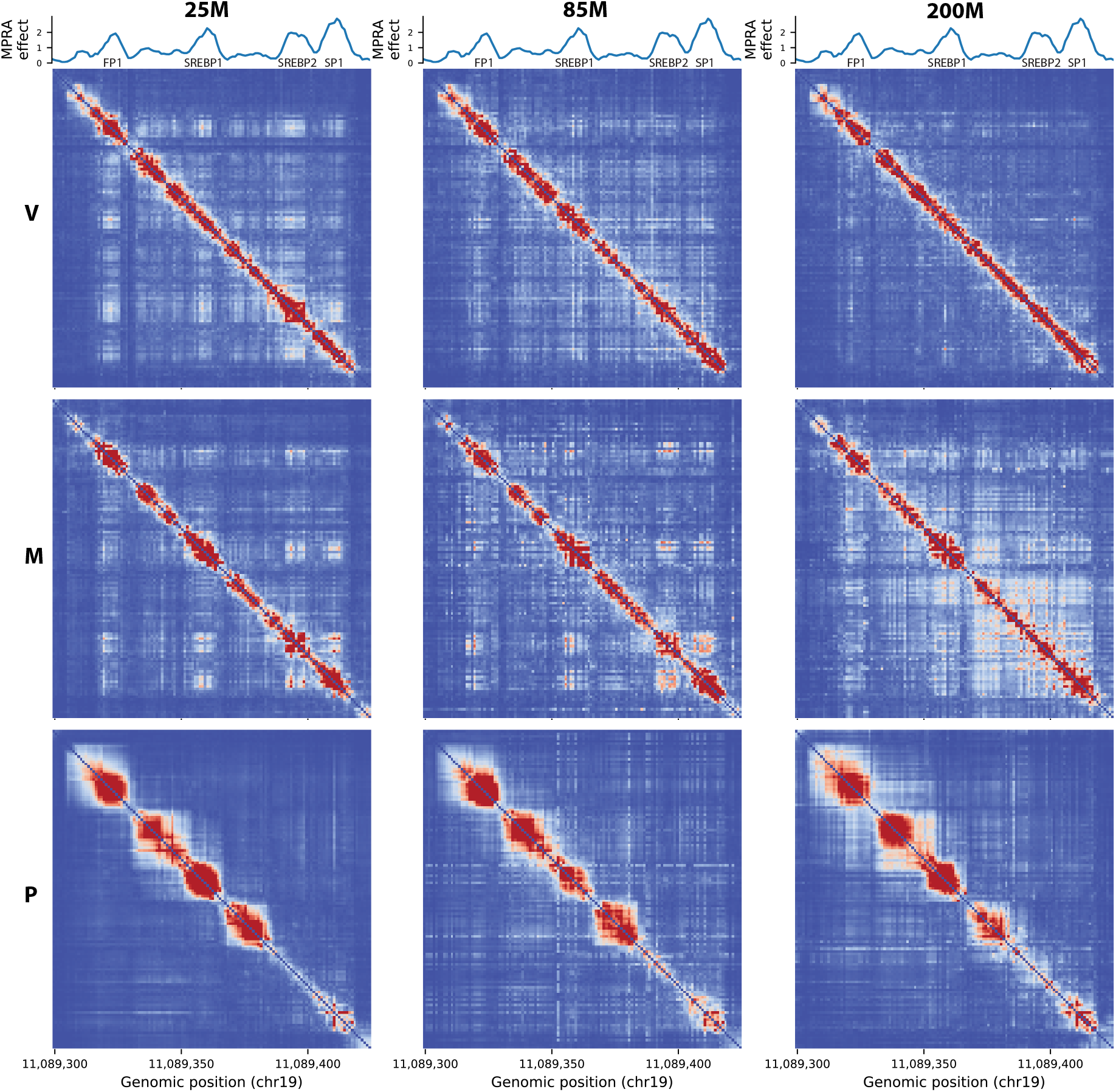
Nucleotide dependency map in *LDLR* promoter (all models). The rows represent models trained on different evolutionary timescales (V: vertebrates, M: mammals, P: primates) and the columns represent different model sizes (number of parameters).

**Supplementary Figure 15:**
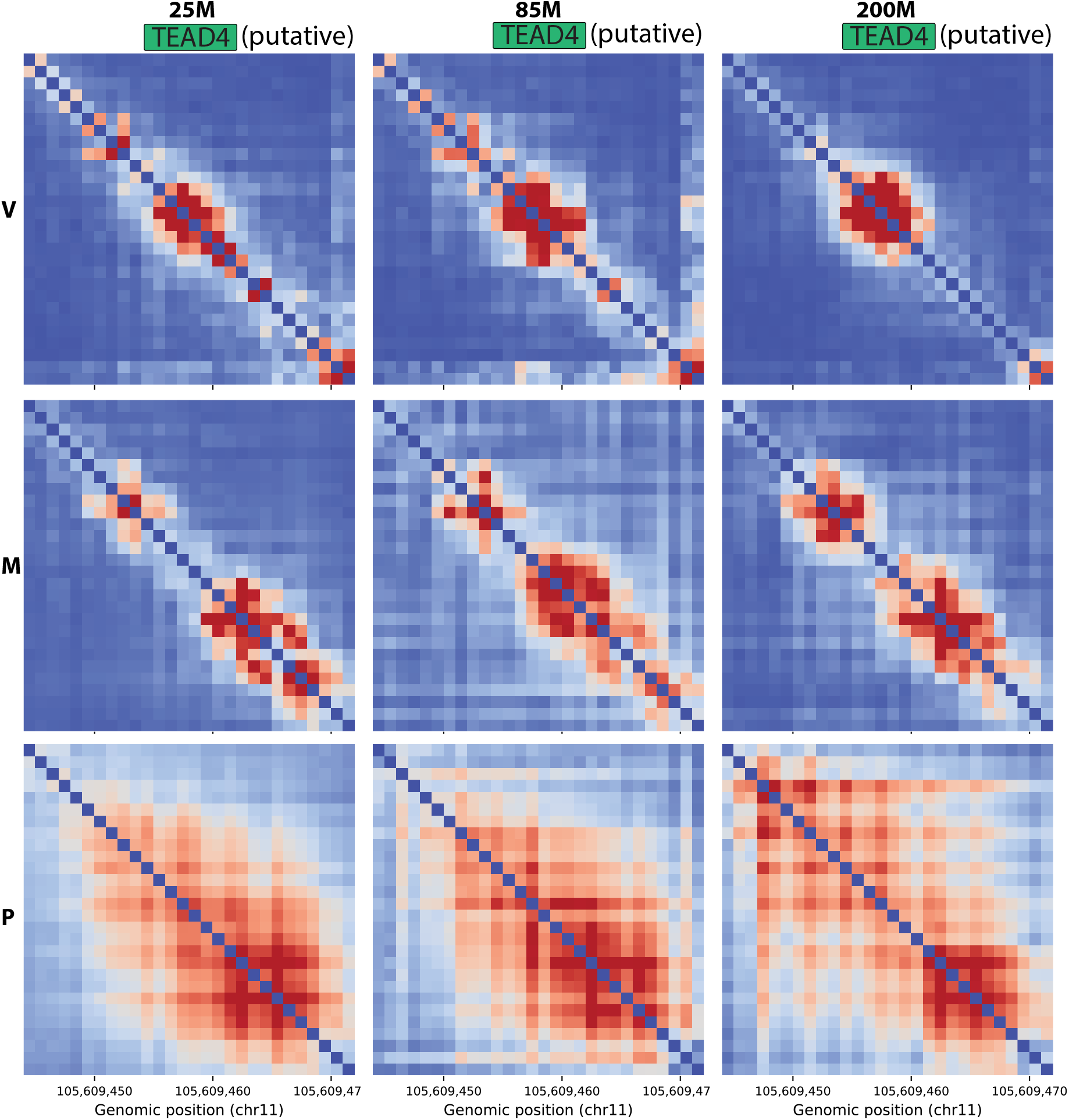
Nucleotide dependency map in a region of putative primate-specific constraint near *GRIA4* (all models). The rows represent models trained on different evolutionary timescales (V: vertebrates, M: mammals, P: primates) and the columns represent different model sizes (number of parameters).

**Supplementary Figure 16:**
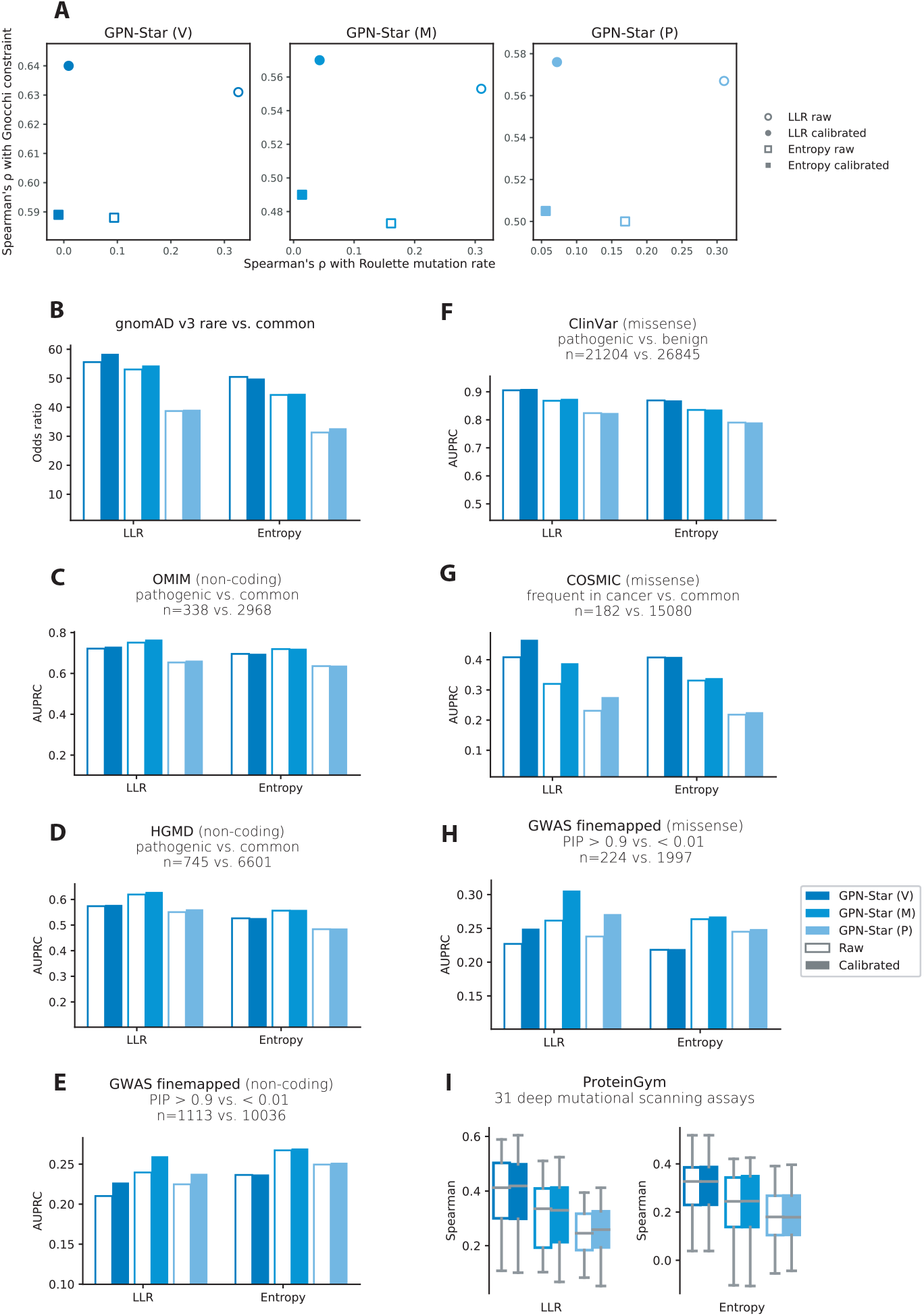
Mutation rate calibration. (A) Spearman correlations between GPN-Star predictions and Roulette mutation rate estimates (*x* axis) and Gnocchi constraint scores (*y* axis) before and after calibration. (B)-(I) Performance of GPN-Star predictions on variant effect prediction benchmarks before and after calibration.

**Supplementary Figure 17:**
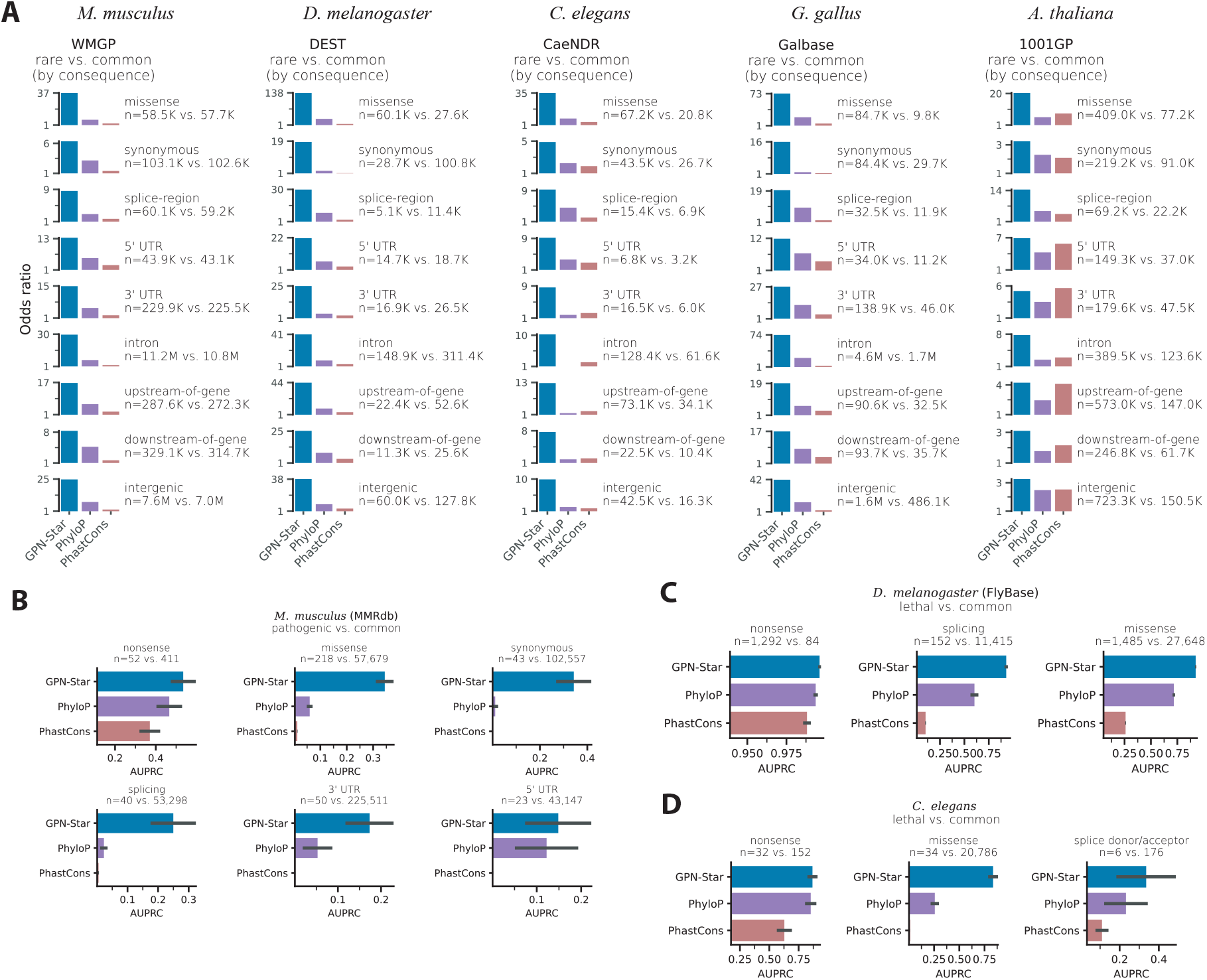
Non-human model evaluations stratified by molecular consequences. (A) Enrichment of rare versus common variants from population genome databases in the tail of deleterious scores (the threshold was chosen such that each score made 30 false discoveries), compared with PhyloP and PhastCons fitted to the same alignments. (B) Classification of MMrdb pathogenic variants versus WMGP common variants in *M. musculus*. (C) Classification of FlyBase lethal variants versus DEST common variants in *D. melanogaster*. (D) Classification of *C. elegans* lethal variants versus CaeNDR common variants.

**Supplementary Figure 18:**
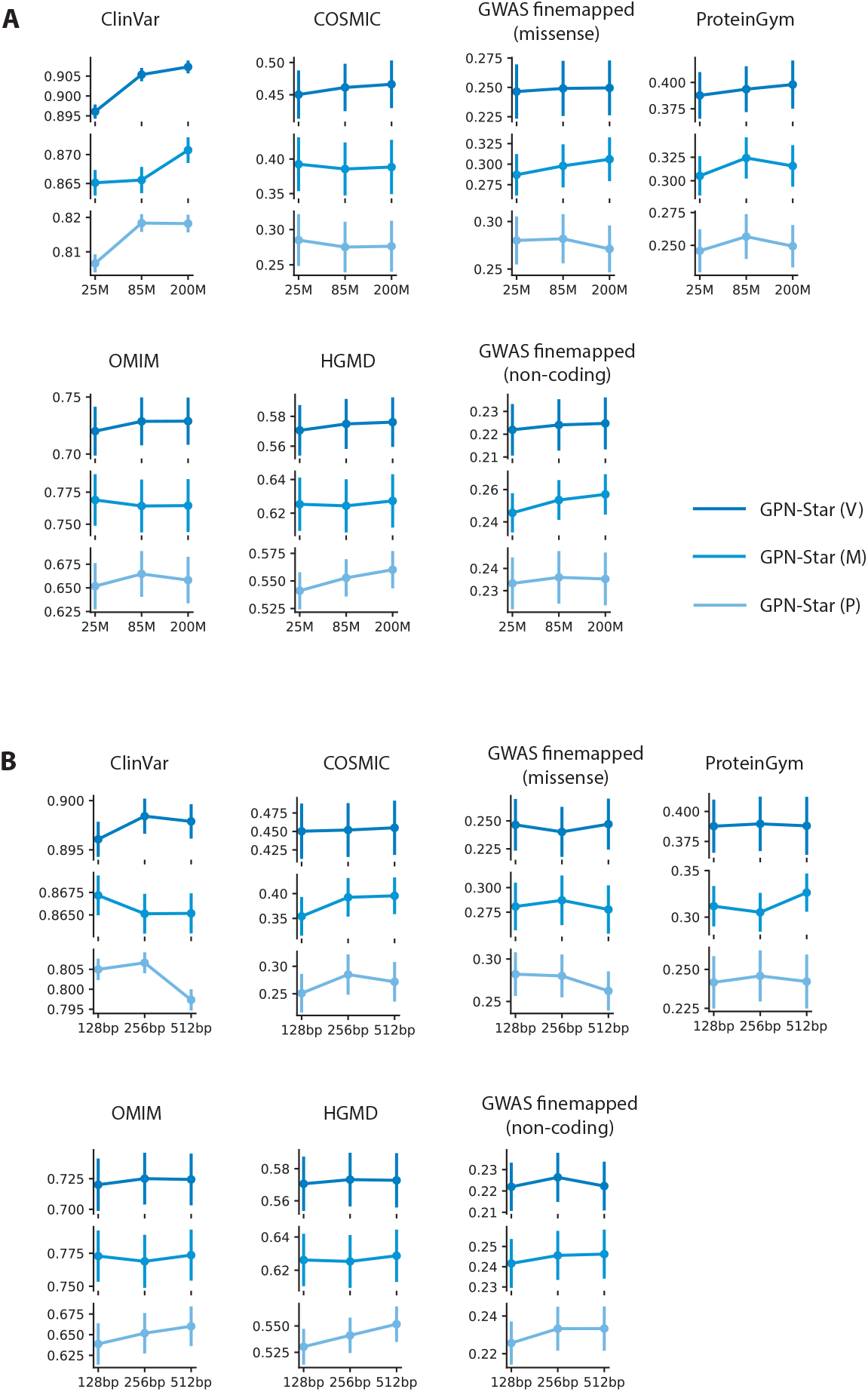
Scaling experiments on model size and context size. (A) Performance of GPN-Star models with 25M, 85M, and 200M parameters across benchmarks. (B) Performance of GPN-Star models with 25M parameters and 128bp, 256bp, and 512bp context sizes across benchmarks. The benchmarks are the same as in Figure 2. For all benchmarks except ProteinGym, the performance metric is the area under precision recall curve (AUPRC) and the error bars represent the standard errors from 1000 bootstrap resamples. For ProteinGym, the dots and error bars represent the mean and standard deviation of Spearman correlations across the DMS assays.

**Supplementary Figure 19:**
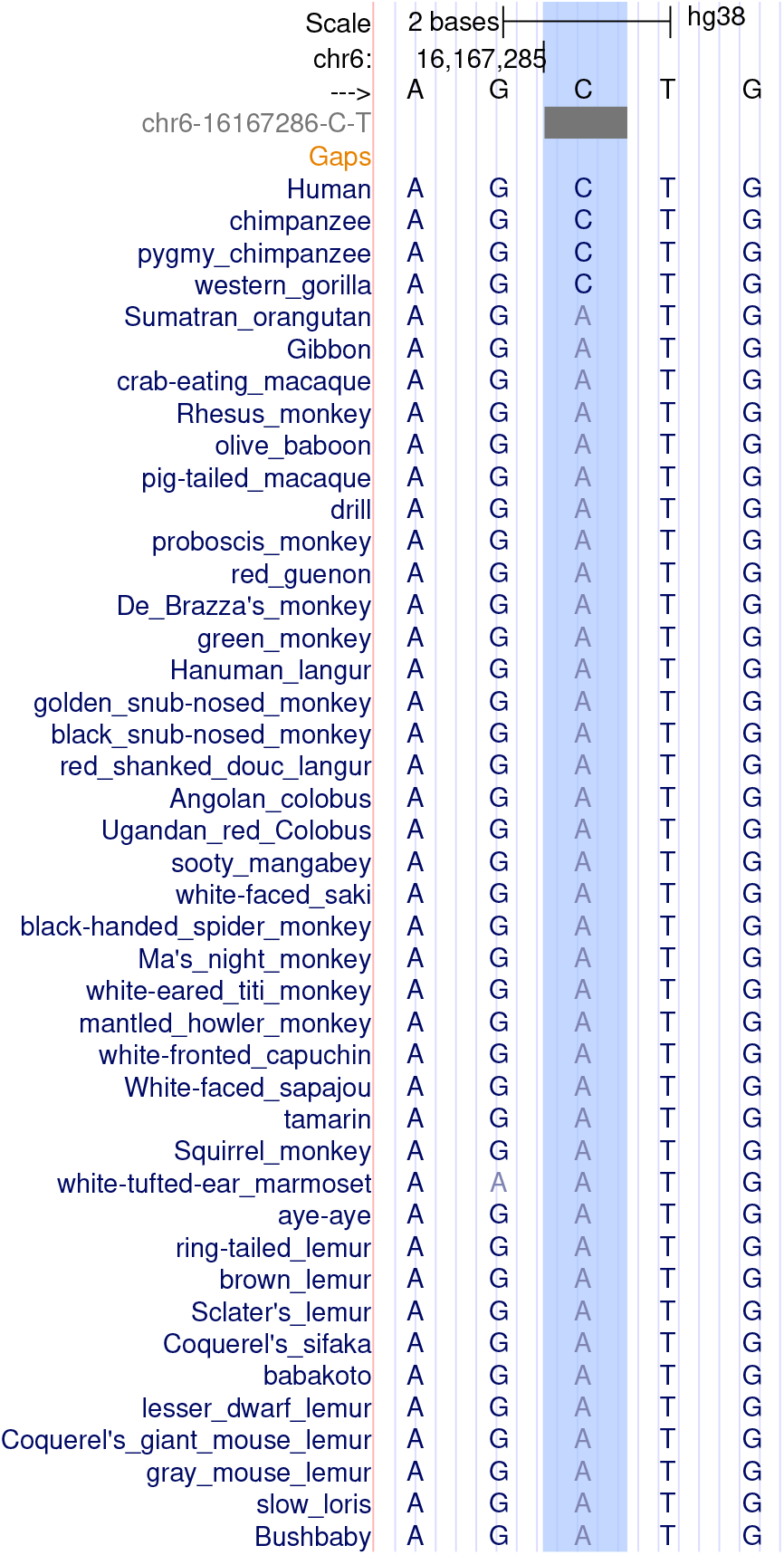
Example putative causal variant (chr6-16167286-C-T) where entropy is more predictive than LLR. The predicted entropy is very low but neither the reference (C) or alternate (T) alleles has a high probability.

**Supplementary Figure 20:**
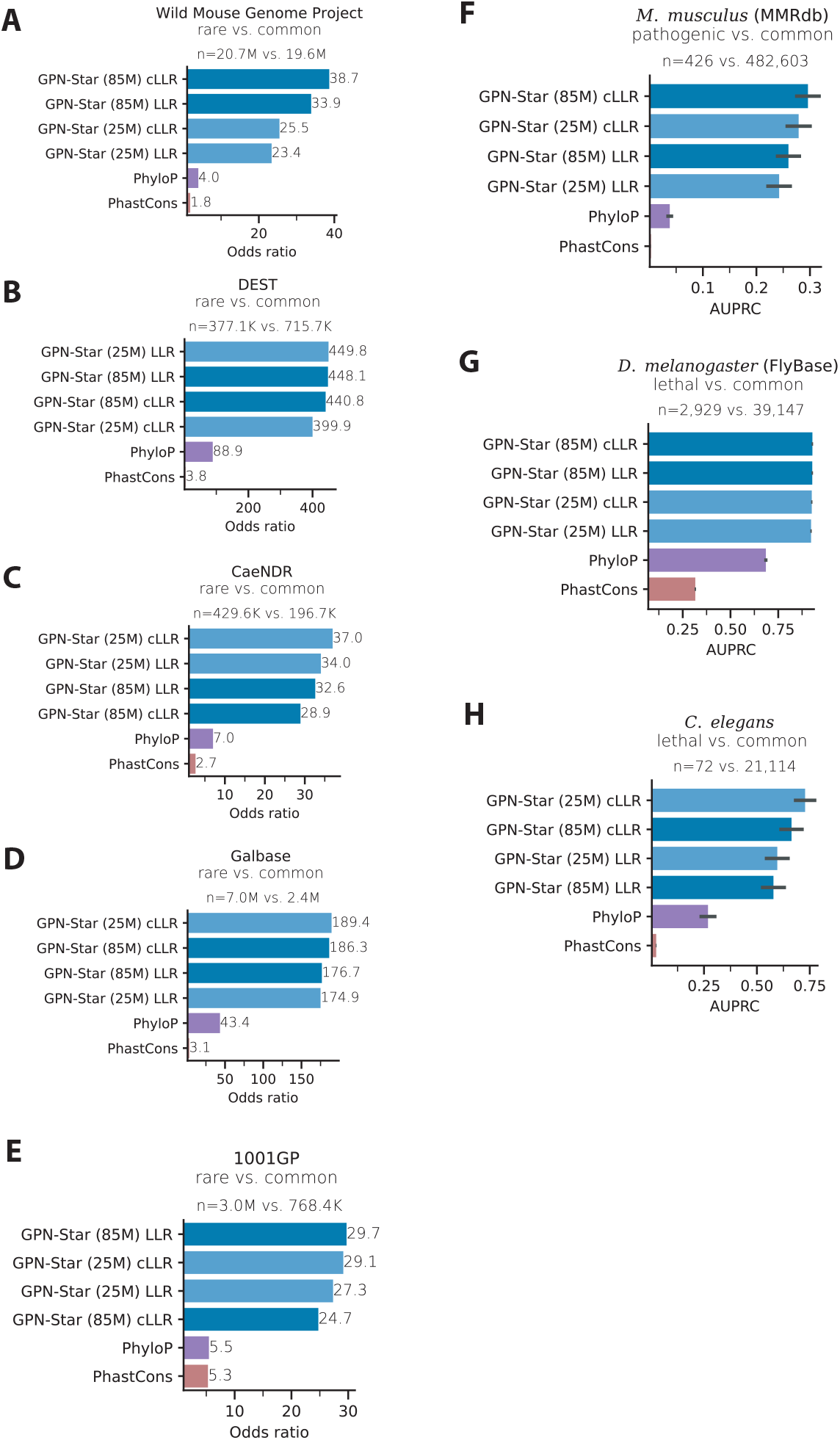
Non-human model evaluations before and after mutation rate calibration. (A)- (E) Enrichment of rare versus common variants from population genome databases in the tail of deleterious scores (the threshold was chosen such that each score made 30 false discoveries), compared with PhyloP and PhastCons fitted to the same alignments. (F) Classification of MMrdb pathogenic variants versus WMGP common variants in *M. musculus*. (G) Classification of FlyBase lethal variants versus DEST common variants in *D. melanogaster*. (H) Classification of *C. elegans* lethal variants versus CaeNDR common variants.

**Supplementary Figure 21:**
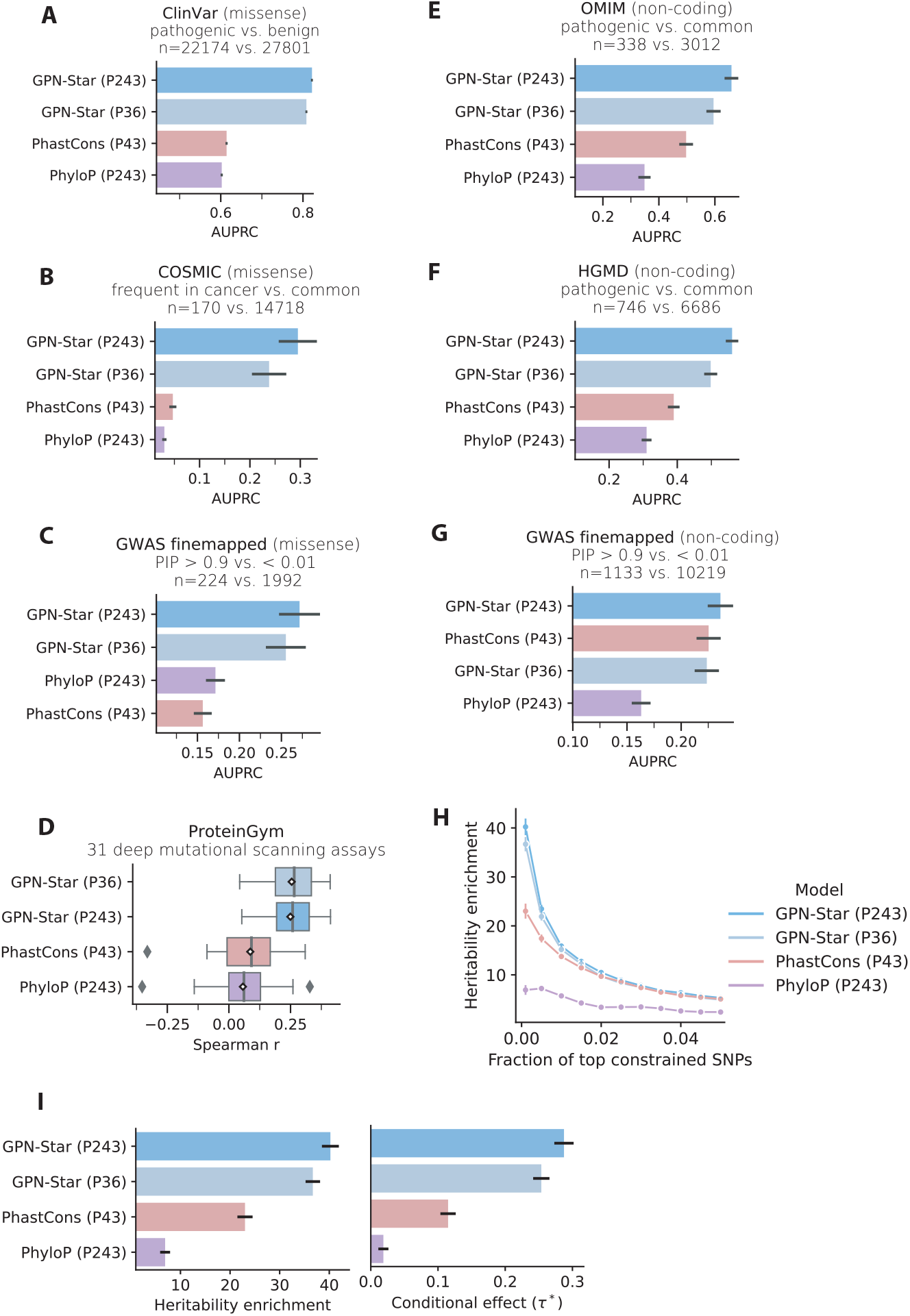
Comparison of the primate models trained with WGAs of different numbers of species. The benchmarks include the variant effect prediction tasks in Figure 2 ((A)-(G)) and complex trait heritability analysis in Figure 3 ((H)-(I)). The numbers in the parentheses in the model names represent the number of primate species in the alignment: P243 denotes the 243 species in the full cactus447way alignment; P43 denotes the 43 species in the cactus241way alignment from the initial Zoonomia release; P36 denotes 36 species in the intersection of the two sets above.

## References

1. Jukes, T. H. & Cantor, C. in Mammalian Protein Metabolism 21–132 (Academic Press, New York, 1969).

2. Dayhoff, M. O., Eck, R. V. & Park, C. M. in Atlas of Protein Sequence and Structure (ed Dayhoff, M.O.) 89–99 (National Biomedical Research Foundation, Washington, DC, 1972).

3. Benegas, G., Ye, C., Albors, C., Li, J. C. & Song, Y. S. Genomic Language Models: Opportunities and Challenges. Trends in Genetics 41, 286–302 (2025).

4. Achiam, J. et al. GPT-4 technical report. arXiv preprint 2303.08774. https://arxiv.org/abs/2303.08774 (2023).

5. Vaswani, A. et al. Attention is All you Need in Advances in Neural Information Processing Systems (eds Guyon, I. et al.) 30 (Curran Associates, Inc., 2017).

6. Benegas, G., Batra, S. S. & Song, Y. S. DNA language models are powerful predictors of genome-wide variant effects. Proceedings of the National Academy of Sciences 120, e2311219120 (2023).

7. Benegas, G., Eraslan, G. & Song, Y. S. Benchmarking DNA Sequence Models for Causal Regulatory Variant Prediction in Human Genetics. bioRxiv preprint. 10.1101/2025.02.11.637758 (2025).

8. Dalla-Torre, H. et al. Nucleotide Transformer: building and evaluating robust foundation models for human genomics. Nature Methods 22, 287–297 (2024).

9. Brixi, G. et al. Genome modeling and design across all domains of life with Evo 2. bioRxiv preprint. 10.1101/2025.02.18.638918 (2025).

10. Jumper, J. et al. Highly accurate protein structure prediction with AlphaFold. Nature 596, 583–589 (2021).

11. Rao, R. M. et al. MSA Transformer in International Conference on Machine Learning (2021), 8844–8856.

12. Frazer, J. et al. Disease variant prediction with deep generative models of evolutionary data. Nature 599, 91–95 (2021).

13. Truong Jr, T. & Bepler, T. PoET: A generative model of protein families as sequences-ofsequences in Advances in Neural Information Processing Systems (eds Oh, A. et al.) 36 (Curran Associates, Inc., 2023), 77379–77415.

14. Ma, C. et al. Retrieved Sequence Augmentation for Protein Representation Learning in Proceedings of the 2024 Conference on Empirical Methods in Natural Language Processing (eds Al-Onaizan, Y., Bansal, M. & Chen, Y.-N.) (Association for Computational Linguistics, Miami, Florida, USA, Nov. 2024), 1738–1767. https://aclanthology.org/2024.emnlp-main.104/.

15. Alamdari, S. et al. Protein generation with evolutionary diffusion: sequence is all you need. bioRxiv, 2023–09 (2023).

16. Li, P., Cheng, X., Song, L. & Xing, E. Retrieval Augmented Protein Language Models for Protein Structure Prediction. bioRxiv, 2024–12 (2024).

17. Sgarbossa, D. & Bitbol, A.-F. RAG-ESM: Improving pretrained protein language models via sequence retrieval. PRX Life 3, 033013 (2025).

18. Yang, K. K. et al. The Dayhoff Atlas: scaling sequence diversity for improved protein generation. bioRxiv, 2025–07 (2025).

19. Akiyama, Y., Zhang, Z., Mirdita, M., Steinegger, M. & Ovchinnikov, S. Scaling down protein language modeling with MSA Pairformer. bioRxiv, 2025–08 (2025).

20. Truong Jr, T. F. & Bepler, T. Understanding protein function with a multimodal retrievalaugmented foundation model. arXiv preprint 2508.04724 (2025).

21. Blanchette, M. et al. Aligning multiple genomic sequences with the threaded blockset aligner. Genome Research 14, 708–715 (2004).

22. Armstrong, J. et al. Progressive Cactus is a multiple-genome aligner for the thousand-genome era. Nature 587, 246–251 (2020).

23. Davydov, E. V. et al. Identifying a high fraction of the human genome to be under selective constraint using GERP++. PLoS computational biology 6, e1001025 (2010).

24. Siepel, A. et al. Evolutionarily conserved elements in vertebrate, insect, worm, and yeast genomes. Genome Research 15, 1034–1050 (2005).

25. Pollard, K. S., Hubisz, M. J., Rosenbloom, K. R. & Siepel, A. Detection of nonneutral substitution rates on mammalian phylogenies. Genome Research 20, 110–121 (2010).

26. Christmas, M. J. et al. Evolutionary constraint and innovation across hundreds of placental mammals. Science 380, eabn3943 (2023).

27. Rhie, A. et al. Towards complete and error-free genome assemblies of all vertebrate species. Nature 592, 737–746 (2021).

28. Stiller, J. et al. Complexity of avian evolution revealed by family-level genomes. Nature 629, 851–860 (2024).

29. Benegas, G., Albors, C., Aw, A. J., Ye, C. & Song, Y. S. A DNA language model based on multispecies alignment predicts the effects of genome-wide variants. Nature Biotechnology. 10.1038/s41587-024-02511-w (2025).

30. Gazal, S. S-LDSC reference files (Zenodo, Jan. 2024). 10.5281/zenodo.10515792.

31. Landrum, M. J. et al. ClinVar: public archive of relationships among sequence variation and human phenotype. Nucleic Acids Research 42, D980–D985 (2014).

32. Tate, J. G. et al. COSMIC: the catalogue of somatic mutations in cancer. Nucleic Acids Research 47, D941–D947 (2019).

33. Karczewski, K. J. et al. The mutational constraint spectrum quantified from variation in 141,456 humans. Nature 581, 434–443 (2020).

34. Notin, P. et al. ProteinGym: Large-Scale Benchmarks for Protein Fitness Prediction and Design in Advances in Neural Information Processing Systems (eds Oh, A. et al.) 36 (Curran Associates, Inc., 2023), 64331–64379. https://proceedings.neurips.cc/paper_files/paper/2023/file/cac723e5ff29f65e3fcbb0739ae91bee-Paper-Datasets_and_Benchmarks.pdf.

35. Cheng, J. et al. Accurate proteome-wide missense variant effect prediction with AlphaMissense. Science 381, eadg7492 (2023).

36. Gao, H. et al. The landscape of tolerated genetic variation in humans and primates. Science 380, eabn8153 (2023).

37. Ghosh, R. et al. Updated recommendation for the benign stand-alone ACMG/AMP criterion. Human mutation 39, 1525–1530 (2018).

38. Avsec, Ž. et al. Effective gene expression prediction from sequence by integrating long-range interactions. Nature Methods 18, 1196–1203 (2021).

39. Linder, J., Srivastava, D., Yuan, H., Agarwal, V. & Kelley, D. R. Predicting RNA-seq coverage from DNA sequence as a unifying model of gene regulation. Nature Genetics. 10.1038/s41588-024-02053-6 (2025).

40. Avsec, Ž. et al. AlphaGenome: advancing regulatory variant effect prediction with a unified DNA sequence model. bioRxiv, 2025–06 (2025).

41. Amberger, J. S., Bocchini, C. A., Schiettecatte, F., Scott, A. F. & Hamosh, A. OMIM.org: Online Mendelian Inheritance in Man (OMIM®), an online catalog of human genes and genetic disorders. Nucleic Acids Research 43, D789–D798 (2015).

42. Stenson, P. D. et al. The Human Gene Mutation Database (HGMD®): optimizing its use in a clinical diagnostic or research setting. Human Genetics 139, 1197–1207 (2020).

43. Avsec, Ž. et al. AlphaGenome: advancing regulatory variant effect prediction with a unified DNA sequence model. bioRxiv. eprint: https://www.biorxiv.org/content/early/2025/07/11/2025.06.25.661532.full.pdf. https://www.biorxiv.org/content/early/2025/07/11/2025.06.25.661532 (2025).

44. Jaganathan, K. et al. Predicting expression-altering promoter mutations with deep learning. Science, eads7373 (2025).

45. Tomaz da Silva, P. et al. Nucleotide dependency analysis of DNA language models reveals genomic functional elements. bioRxiv preprint, 2024–07. https://www.biorxiv.org/content/10.1101/2024.07.27.605418v1 (2024).

46. Kanai, M. et al. Insights from complex trait fine-mapping across diverse populations. medrxiv, 2021–09 (2021).

47. Bomba, L., Walter, K. & Soranzo, N. The impact of rare and low-frequency genetic variants in common disease. Genome biology 18, 77 (2017).

48. Lee, S., Abecasis, G. R., Boehnke, M. & Lin, X. Rare-variant association analysis: study designs and statistical tests. The American Journal of Human Genetics 95, 5–23 (2014).

49. Clarke, B. et al. Integration of variant annotations using deep set networks boosts rare variant association testing. Nature Genetics, 1–10 (2024).

50. Backman, J. D. et al. Exome sequencing and analysis of 454,787 UK Biobank participants. Nature 599, 628–634 (2021).

51. Karczewski, K. J. et al. Systematic single-variant and gene-based association testing of thousands of phenotypes in 394,841 UK Biobank exomes. Cell genomics 2 (2022).

52. Finucane, H. K. et al. Partitioning heritability by functional annotation using genome-wide association summary statistics. Nature Genetics 47, 1228–1235 (2015).

53. Weissbrod, O. et al. Functionally informed fine-mapping and polygenic localization of complex trait heritability. Nature Genetics 52, 1355–1363 (2020).

54. Márquez-Luna, C. et al. Incorporating functional priors improves polygenic prediction accuracy in UK Biobank and 23andMe data sets. Nature Communications 12, 6052 (2021).

55. Sullivan, P. F. et al. Leveraging base-pair mammalian constraint to understand genetic variation and human disease. Science 380, eabn2937 (2023).

56. Kuderna, L. F. et al. Identification of constrained sequence elements across 239 primate genomes. Nature 625, 735–742 (2024).

57. O’Connor, L. J. & Sella, G. Principled measures and estimates of trait polygenicity. bioRxiv, 2025–07 (2025).

58. Dyer, S. C. et al. Ensembl 2025. Nucleic Acids Research 53, D948–D957 (2025).

59. Moore, J. E. et al. An Expanded Registry of Candidate cis-Regulatory Elements for Studying Transcriptional Regulation. bioRxiv (2024).

60. Karollus, A., Mauermeier, T. & Gagneur, J. Current sequence-based models capture gene expression determinants in promoters but mostly ignore distal enhancers. Genome Biology 24, 56 (2023).

61. Fabiha, T. et al. A consensus variant-to-function score to functionally prioritize variants for disease. bioRxiv, 2024–11 (2024).

62. Finucane, H. K. et al. Heritability enrichment of specifically expressed genes identifies diseaserelevant tissues and cell types. Nature Genetics 50, 621–629 (2018).

63. Zhang, Z. et al. Protein language models learn evolutionary statistics of interacting sequence motifs. Proceedings of the National Academy of Sciences 121, e2406285121 (2024).

64. Verbeek, M. M. et al. Mutations in the cyclic adenosine monophosphate response element of the tyrosine hydroxylase gene. Annals of Neurology 62, 422–426 (2007).

65. Dong, H.-Y., Feng, J.-Y., Yue, X.-J., Shan, L. & Jia, F.-Y. Dopa-responsive dystonia caused by tyrosine hydroxylase deficiency: three cases report and literature review. Medicine 99, e21753 (2020).

66. Ribasés, M. et al. A homozygous tyrosine hydroxylase gene promoter mutation in a patient with dopa-responsive encephalopathy: clinical, biochemical and genetic analysis. Molecular Genetics and Metabolism 92, 274–277 (2007).

67. Stamelou, M. et al. Myoclonus-dystonia syndrome due to tyrosine hydroxylase deficiency. Neurology 79, 435–441 (2012).

68. Kircher, M. et al. Saturation mutagenesis of twenty disease-associated regulatory elements at single base-pair resolution. Nature Communications 10 (2019).

69. Khamis, A. et al. Functional analysis of four LDLR 5 UTR and promoter variants in patients with familial hypercholesterolaemia. European Journal of Human Genetics 23, 790– 795 (2015).

70. Bennett, M. K., Ngo, T. T., Athanikar, J. N., Rosenfeld, J. M. & Osborne, T. F. Costimulation of promoter for low density lipoprotein receptor gene by sterol regulatory elementbinding protein and Sp1 is specifically disrupted by the yin yang 1 protein. Journal of Biological Chemistry 274, 13025–13032 (1999).

71. Seplyarskiy, V. et al. A mutation rate model at the basepair resolution identifies the mutagenic effect of Polymerase III transcription. Nature Genetics, 1–8 (2023).

72. Consortium, E. P. et al. An integrated encyclopedia of DNA elements in the human genome. Nature 489, 57–74 (2012).

73. Lonsdale, J. et al. The genotype-tissue expression (GTEx) project. Nature Genetics 45, 580– 585 (2013).

74. Song, B., Buckler, E. S. & Stitzer, M. C. New whole-genome alignment tools are needed for tapping into plant diversity. Trends in Plant Science 29, 355–369 (2024).

75. Öztürk-Çolak, A. et al. FlyBase: updates to the Drosophila genes and genomes database. Genetics 227, iyad211 (2024).

76. Qin, Z. et al. Genomic identification and functional characterization of essential genes in Caenorhabditis elegans. G3: Genes, Genomes, Genetics 8, 981–997 (2018).

77. Small, S., Blair, A. & Levine, M. Regulation of even-skipped stripe 2 in the Drosophila embryo. The EMBO journal 11, 4047–4057 (1992).

78. Bothma, J. P. et al. Dynamic regulation of eve stripe 2 expression reveals transcriptional bursts in living Drosophila embryos. Proceedings of the National Academy of Sciences 111, 10598–10603 (2014).

79. Rives, A. et al. Biological structure and function emerge from scaling unsupervised learning to 250 million protein sequences. Proceedings of the National Academy of Sciences 118, e2016239118 (2021).

80. Lin, Z. et al. Evolutionary-scale prediction of atomic-level protein structure with a language model. Science 379, 1123–1130 (2023).

81. Of Life Project Consortium, D. T. Sequence locally, think globally: the Darwin Tree of Life Project. Proceedings of the National Academy of Sciences 119, e2115642118 (2022).

82. Lewin, H. A. et al. The earth BioGenome project 2020:Starting the clock 2022.

83. Miles, A. et al. zarr-developers/zarr-python: v3.0.7 version v3.0.7. Apr. 2025. 10.5281/zenodo.15255977.

84. Kent, W. J. et al. The human genome browser at UCSC. Genome Research 12, 996–1006 (2002).

85. A comparative genomics multitool for scientific discovery and conservation. Nature 587, 240–245 (2020).

86. Albors, C., Li, J. C., Benegas, G., Ye, C. & Song, Y. S. A Phylogenetic Approach to Genomic Language Modeling in Research in Computational Molecular Biology (ed Sankararaman, S.) (Springer Nature Switzerland, Cham, 2025), 99–117. isbn: 978-3-031-90252-9.

87. Tian, F., Yang, D.-C., Meng, Y.-Q., Jin, J. & Gao, G. PlantRegMap: charting functional regulatory maps in plants. Nucleic Acids Research 48, D1104–D1113 (2020).

88. Hubisz, M. J., Pollard, K. S. & Siepel, A. PHAST and RPHAST: phylogenetic analysis with space/time models. Briefings in Bioinformatics 12, 41–51 (2011).

89. Su, J. et al. Roformer: Enhanced transformer with rotary position embedding. arXiv preprint 2104.09864 (2021).

90. Li, S. et al. Functional Interpolation for Relative Positions improves Long Context Transformers in The Twelfth International Conference on Learning Representations (2024). https://openreview.net/forum?id=rR03qFesqk.

91. Aggarwala, V. & Voight, B. F. An expanded sequence context model broadly explains variability in polymorphism levels across the human genome. Nature Genetics 48, 349–355 (2016).

92. Tarailo-Graovac, M. & Chen, N. Using RepeatMasker to Identify Repetitive Elements in Genomic Sequences. Current Protocols in Bioinformatics 25, 4.10.1–4.10.14. eprint: https://currentprotocols.onlinelibrary.wiley.com/doi/pdf/10.1002/0471250953.bi0410s25. https://currentprotocols.onlinelibrary.wiley.com/doi/abs/10.1002/0471250953.bi0410s25 (2009).

93. Open2C et al. Bioframe: operations on genomic intervals in Pandas dataframes. Bioinformatics 40, btae088 (2024).

94. McInnes, L., Healy, J. & Melville, J. UMAP: Uniform manifold approximation and projection for dimension reduction. arXiv preprint 1802.03426 (2018).

95. Schubach, M., Maass, T., Nazaretyan, L., Röner, S. & Kircher, M. CADD v1.7: using protein language models, regulatory CNNs and other nucleotide-level scores to improve genome-wide variant predictions. Nucleic Acids Research 52, D1143–D1154 (2024).

96. Chen, K. M., Wong, A. K., Troyanskaya, O. G. & Zhou, J. A sequence-based global map of regulatory activity for deciphering human genetics. Nature Genetics 54, 940–949 (2022).

97. Bycroft, C. et al. The UK Biobank resource with deep phenotyping and genomic data. Nature 562, 203–209 (2018).

98. Chen, S. et al. A genome-wide mutational constraint map quantified from variation in 76,156 human genomes. bioRxiv, 2022–03 (2022).

99. Brandes, N., Goldman, G., Wang, C. H., Ye, C. J. & Ntranos, V. Genome-wide prediction of disease variant effects with a deep protein language model. Nature Genetics. issn: 1546-1718. 10.1038/s41588-023-01465-0 (Aug. 2023).

100. Halligan, D. L. et al. Contributions of protein-coding and regulatory change to adaptive molecular evolution in murid rodents. PLoS genetics 9, e1003995 (2013).

101. Davies, R. W. Factors influencing genetic variation in wild mice PhD thesis (University of Oxford, 2015).

102. Harr, B. et al. Genomic resources for wild populations of the house mouse, Mus musculus and its close relative Mus spretus. Scientific data 3, 1–14 (2016).

103. Phifer-Rixey, M. et al. The genomic basis of environmental adaptation in house mice. PLoS Genetics 14, e1007672 (2018).

104. Payseur, B. A. & Jing, P. Genomic targets of positive selection in giant mice from Gough Island. Molecular Biology and Evolution 38, 911–926 (2021).

105. Fujiwara, K. et al. Insights into Mus musculus population structure across Eurasia revealed by whole-genome analysis. Genome biology and evolution 14, evac068 (2022).

106. Morgan, A. P. et al. Population structure and inbreeding in wild house mice (Mus musculus) at different geographic scales. Heredity 129, 183–194 (2022).

107. Lawal, R. A. et al. Taxonomic assessment of two wild house mouse subspecies using wholegenome sequencing. Scientific reports 12, 20866 (2022).

108. Chen, S., Zhou, Y., Chen, Y. & Gu, J. fastp: an ultra-fast all-in-one FASTQ preprocessor. Bioinformatics 34, i884–i890 (2018).

109. Hickey, G. et al. Pangenome graph construction from genome alignments with MinigraphCactus. Nature biotechnology 42, 663–673 (2024).

110. Ferraj, A. et al. Resolution of structural variation in diverse mouse genomes reveals chromatin remodeling due to transposable elements. Cell Genomics 3 (2023).

111. Sirén, J. et al. Pangenomics enables genotyping of known structural variants in 5202 diverse genomes. Science 374, abg8871 (2021).

112. Garrison, E. et al. Variation graph toolkit improves read mapping by representing genetic variation in the reference. Nature biotechnology 36, 875–879 (2018).

113. Poplin, R. et al. A universal SNP and small-indel variant caller using deep neural networks. Nature biotechnology 36, 983–987 (2018).

114. Lin, M. F. et al. GLnexus: joint variant calling for large cohort sequencing. BioRxiv, 343970 (2018).

115. Fairfield, H. et al. Mutation discovery in mice by whole exome sequencing. Genome biology 12, 1–12 (2011).

116. Fairfield, H. et al. Exome sequencing reveals pathogenic mutations in 91 strains of mice with Mendelian disorders. Genome research 25, 948–957 (2015).

117. Perez, G. et al. The UCSC Genome Browser database: 2025 update. Nucleic Acids Research 53, D1243–D1249 (2025).

118. Nunez, J. C. et al. Footprints of worldwide adaptation in structured populations of D. melanogaster through the expanded DEST 2.0 genomic resource. bioRxiv, 2024–11 (2024).

119. Crombie, T. A. et al. CaeNDR, the Caenorhabditis natural diversity resource. Nucleic Acids Research 52, D850–D858 (2024).

120. Fu, W. et al. Galbase: a comprehensive repository for integrating chicken multi-omics data. BMC genomics 23, 364 (2022).

121. Alonso-Blanco, C. et al. 1,135 genomes reveal the global pattern of polymorphism in Arabidopsis thaliana. Cell 166, 481–491 (2016).

122. Gazal, S. et al. Linkage disequilibrium–dependent architecture of human complex traits shows action of negative selection. Nature Genetics 49, 1421–1427 (2017).

123. 1000 Genomes Project Consortium et al. A global reference for human genetic variation. Nature 526, 68–74 (2015).

